# Innovation within stability: balancing selection preserves multigenic S-haplotype architecture in apple

**DOI:** 10.1101/2025.10.28.683340

**Authors:** Aurélie Mesnil, Vincent Castric, Gianluca Lombardi, Thomas Blein, Jia-Long Yao, Charles Ampomah-Dwamena, Marie Monniaux, Somia Saidi, Xilong Chen, William Marande, Caroline Callot, Nadine Gautier, Johann Confais, Anne-Laure Fuchs, Enrique Dapena, Anthony Venon, Takashi Tsuchimatsu, Alessandra Carbone, Xavier Vekemans, Amandine M. Cornille

## Abstract

Self-incompatibility (SI) systems prevent self-fertilization, thereby maintaining genetic diversity in flowering plants. Among them, collaborative non-self recognition (CNSR) is the most widespread, yet the genomic organization and evolutionary maintenance of its multigenic recognition system remain poorly understood. Using 27 haplotype-resolved genomes from wild and domesticated apples (*Malus* spp.), we dissected the structure and evolution of the S-locus. We identified 17 S-RNase alleles and 500 pollen-expressed S-locus F-box brother (SFBB) genes across 18 families. The S-locus shows extensive structural divergence among alleles and transposable element accumulation, consistent with long-term restricted effective recombination. Despite this divergence, haplotypes carrying the same S-RNase allele retain remarkably conserved SFBB repertoires and gene organization, even across species boundaries, indicating that long-term balancing selection preserves highly conserved S-haplotype architectures associated with specific S-RNase lineages. Tandem duplication, positive selection, and signatures consistent with gene conversion contribute to the diversification of pollen-expressed SFBB genes while S-RNase-associated SFBB repertoires remain conserved across haplotypes carrying the same S-RNase allele. Our results reveal how a structurally dynamic yet evolutionarily constrained genomic region can sustain long-term S-allele diversity and preserve complex multigenic haplotype architectures in flowering plants.

## Main

Recognition systems are fundamental to biological interactions, enabling organisms to distinguish self from non-self across contexts as diverse as immunity, mating, and symbiosis. A central evolutionary challenge for such systems is generating novel recognition specificities while preserving compatibility with existing partners. Understanding how recognition systems remain evolvable without losing functionality remains a major question in evolutionary biology.

Self-incompatibility (SI) systems provide one of the best natural models for addressing this question. By preventing self-fertilization, SI promotes outcrossing and maintains genetic diversity over long evolutionary timescales^1–3^. In the Maleae tribe of the Rosaceae family, including apple (*Malus* spp.), SI is controlled by an S-RNase-based collaborative non-self recognition (CNSR) system in which a pistil-expressed S-RNase acts as a toxin and multiple pollen-expressed S-locus F-box brothers (SFBBs) function collectively as detoxification factors^4–7^. Unlike single-gene recognition systems, CNSR relies on the coordinated action of numerous linked pollen determinants, creating a multigenic recognition architecture. Regulatory elements beyond canonical coding determinants may also contribute to S-locus function; non-coding sequences of S-RNase genes in *Fragaria* have been shown to produce lncRNAs, hinting at additional regulatory layers^8^.

The multigenic nature of CNSR creates a fundamental evolutionary paradox. New pollen-recognition specificities must continually emerge to accommodate the diversification of S-RNases, yet recognition repertoires must remain sufficiently complete to maintain compatibility with most potential mates^9–11^. Theoretical studies predict that balancing selection should favor S-haplotypes carrying nearly complete repertoires of pollen determinants capable of detoxifying most non-self S-Rnases^9–11^. However, recent models suggest that the gain, loss, and exchange of recognition genes may generate substantial variation among haplotypes that carry the same S-RNase lineage^11^. Consequently, it remains unclear whether balancing selection primarily preserves individual SI determinants or larger multigenic recognition architectures. This question is particularly relevant because S-loci often exhibit genomic features associated with regions of long-term restricted recombination, including extensive structural polymorphism, accumulation of transposable elements, and reduced effective recombination^12–15^. Such genomic environments may facilitate diversification while simultaneously increasing the risk of degeneration. The mechanisms that allow multigenic recognition systems to remain functional despite these opposing evolutionary forces are poorly understood^16,17^. Previous work in *Petunia* and *Antirrhinum* has shown that duplication, gene conversion, and sequence diversification contribute to the evolution of pollen determinants^16,18,19^. However, whether balancing selection preserves individual recognition genes or entire co-adapted recognition modules remains unresolved. Distinguishing among these alternatives requires haplotype-resolved analyses that simultaneously capture gene content, structural organization, and evolutionary relationships across complete S-haplotypes.

Here, using 27 phased S-haplotypes from cultivated apple and its wild relatives, we reconstruct the genomic architecture and evolutionary history of the apple S-locus. We identify 17 S-RNase lineages and 500 SFBB genes across 18 families, and show that S-RNase lineages remain tightly associated with highly conserved SFBB repertoires and gene organization. Despite extensive sequence divergence, structural rearrangements, transposable element accumulation, and ongoing diversification through duplication, positive selection, and sequence exchange, balancing selection preserves remarkably stable multigenic recognition modules across species boundaries. These results reveal how long-term balancing selection can maintain complex recognition architectures while allowing continual innovation of their constituent genes.

## Results

### Long-term balancing selection preserves S-haplotype architecture

To understand how collaborative non-self recognition systems remain evolutionarily stable despite continual diversification of recognition specificities, we reconstructed and compared 27 complete Shaplotypes from 14 *M. domestica* (12 dessert, two cider) and 13 wild relatives (*M. sylvestris, M. sieversii, M. orientalis*, Supplementary Table 1). This haplotype-resolved dataset allowed us to investigate how balancing selection, structural variation, and gene turnover collectively shape the architecture and evolution of the apple S-locus. At the S-locus on chromosome 17, each haplotype contained a single S-RNase gene. Combined analysis of our 27 sequences with public database sequences (Supplementary Table 2) identified 17 clearly distinct allelic lineages (Supplementary Table 3, Supplementary Fig. 1A). A 100% amino acid identity was typically observed between gene copies within alleles (Supplementary Table 4, Supplementary Fig. 1B), suggesting strong purifying selection. One exception, haplotype h32 from *M. orientalis*, exhibited only 89.1% similarity with two MdS9 allele sequences from *M. domestica*. In contrast, pairwise identity between S-RNase sequences of different alleles averaged 65.8% (range 53.2–94.8%, n = 334 pairs), consistent with diversifying selection maintaining distinct S-specificities. Controlled pollinations are necessary to confirm phenotypic recognition equivalence. Several wild haplotypes carried previously uncharacterized alleles (h24, h25, h29, h31). The *S*-RNase phylogeny (Supplementary Fig. 2A) was incongruent with the species tree (based on chromosome 17 single-copy genes), showing extensive interspecific allele sharing. Several S-RNase lineages were found across multiple Malus species, suggesting either long-term trans-species polymorphism, as expected under balancing selection^20^, or adaptive introgression among species^1,21^.

We next characterized the pollen component of the SI system. Across the 27 S-haplotypes, we identified 500 intron-less SFBB genes (17–20 per haplotype; ∼1.2 kb each), all encoding a conserved N-terminal F-box domain and a C-terminal FBA domain (Supplementary Note 1, Supplementary Table 8). Phylogenetic analyses grouped these genes into 18 distinct SFBB families (SFBB1-SFBB18; Supplementary Note 1, Supplementary Table 7, Supplementary Fig. 3).

We then examined how SFBB repertoires varied among S-haplotypes (Fig. 1A, B). Each haplotype carried a characteristic combination of SFBB families that was strongly associated with its S-RNase allele (Mantel test, *p*-value = 0.024, Fig. 1B) despite belonging to different *Malus* species. Pairwise comparisons further confirmed this pattern. Haplotypes carrying the same S-RNase allele exhibited significantly more similar SFBB repertoires than haplotypes carrying different S-RNase alleles (Wilcoxon test, *p* = 1.16 × 10^−6^; Fig. 1D), irrespective of species identity. This result provides direct quantitative support for the association between S-RNase identity and SFBB repertoire composition. For example, the *M. orientalis* h32 SFBB repertoire was almost indistinguishable from that of the two *M. domestica MdS9* haplotypes, differing only by a duplication of SFBB3. These observations indicate that S-RNase identity is a strong predictor of SFBB repertoire composition across species boundaries.

**Fig. 1:**
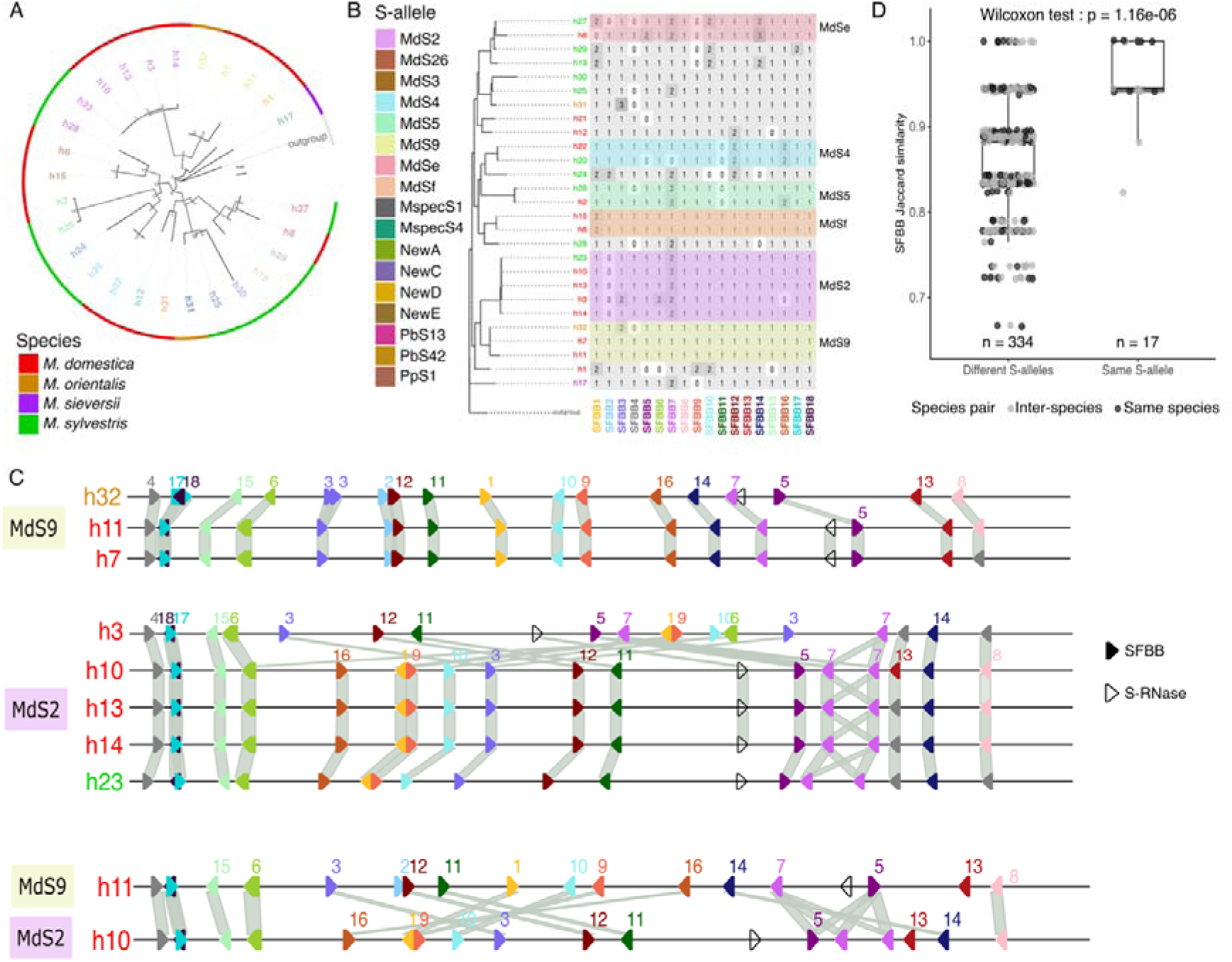
Balancing selection preserves complete S-haplotype architectures. (**A**) Maximum-likelihood circular phylogenetic tree of S-RNase protein sequences, rooted with Eriobotrya japonica S6 S-RNase. Branch width is proportional to ultrafast bootstrap support (UFBoot; IQ-TREE2). The 27 Malus haplotypes analyzed in this study are coloured by species: M. domestica (red), M. sylvestris (green), M. orientalis (orange), and M. sieversii (purple). (**B**) SFBB gene family copy number per haplotype. Rows correspond to the 27 Malus haplotypes ordered as in (A); columns correspond to SFBB gene families 1–18 (SFBB1–SFBB18), each identified by a unique color. Cell fill and number indicate the copy count carried by each haplotype (white = 0, grey ≥ 1). (**C**) Gene organization at the S-locus for representative haplotypes from two S-allele groups (MdS9, top; MdS2, bottom). Filled arrows represent SFBB genes (colors indicate orthologous families); open arrows represent the S-RNase gene. Syntenic connections between haplotypes are shown by shaded ribbons; scale bar indicates genomic distance (kb). (**D**) SFBB pairwise amino acid similarity as a function of S-RNase identity. Each point represents a haplotype pair, colored by comparison type (inter-species or same species). Haplotype pairs sharing the same S-RNase (right) show significantly higher SFBB similarity than pairs with different S-RNases (left; Wilcoxon test, *p* = 1.16 × 10^−6^), suggesting co-evolution between S-RNase identity and SFBB repertoire within each S-allele group.

This conservation extended beyond gene content. The SFBB gene order was highly conserved among haplotypes carrying the same S-RNase allele (Fig. 1C), with only minor local differences that may reflect assembly quality (e.g., h3 vs. other *MdS2* haplotypes), consistent with a lower quality assembly index (LAI) for this haplotype (Supplementary Fig. 5). Whole-locus alignments revealed extensive synteny and low structural divergence among haplotypes sharing the same S-RNase allele, even when sampled from different species (Fig. 1C).

By contrast, haplotypes carrying different *S*-RNases exhibited markedly divergent architectures (Fig. 1C and 1D). Although a few SFBB blocks (e.g., SFBB4 to SFBB6 and SFBB5 to SFBB14) remained conserved near the locus boundaries, central regions showed frequent inversions and unaligned segments (Supplementary Fig. 6). Consistent with this pattern, haplotypes sharing the same S-RNase allele displayed significantly longer syntenic blocks (*p* < 2.2e−16), whereas haplotypes carrying different alleles exhibited significantly larger inversions (*p* = 1.3e−4).

We next investigated whether this conservation of S-RNase–SFBB modules extends to the broader genomic architecture of the S-locus^5^. We identified the limits of the S-locus using conserved flanking genes^15^, including an upstream NF-Y gene and two downstream PI3/PI4 kinase family genes. The locus averaged 1.35 Mb (0.92 to 1.87 Mb). It contained about 42 annotated genes, approximately three times fewer than observed in matched control regions (Fig. 2A, Supplementary Fig. 7). The S-locus was also highly enriched in TEs, particularly LTRs (Fig. 2B, ∼85% *vs*. 70% genome-wide; Supplementary Fig. 8; Wilcoxon-Mann-Whitney test, *p*-value = 2.9e−10). *S*-locus annotated genes were grouped into 55 orthologous clusters using Syntenet^22^ and classified into four categories (Supplementary Fig. 9, Supplementary Table 9). Apart from SFBB and *S*-RNase genes, 38 clusters encoded diverse functions (e.g., kinases, transcription factors, transporters), suggesting possible recruitment or retention driven by structural dynamics or regulation. Gene content clustering across haplotypes revealed both conserved and variable clusters (Supplementary Fig. 9), with high TE content and variable gene presence, reinforcing the understanding of the *S*-locus as a structurally dynamic region.

**Fig. 2:**
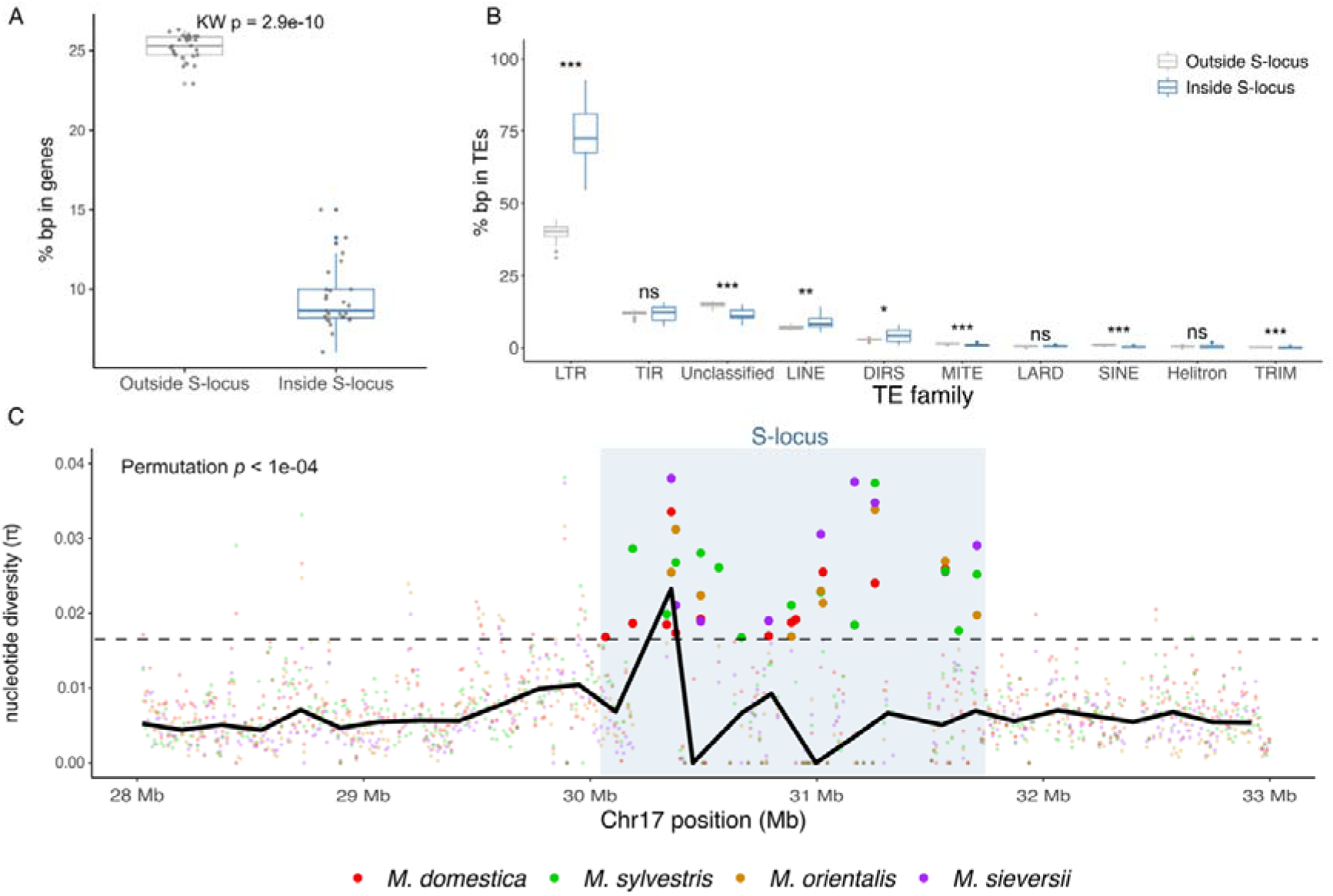
Characteristics of the self-incompatibility locus on chromosome 17. (**A**) Percentage of base pairs covered by annotated genes inside versus outside the S-locus, across all haplotypes. (**B**) Percentage of base pairs covered by transposable elements (TEs) inside versus outside the Slocus, separated by TE order. For panels C and D, boxes show the interquartile range and median across haplotypes; dots represent individual haplotypes. Grey = outside S-locus, blue = inside Slocus. Significance of inside/outside differences was assessed with a Kruskal-Wallis test (* p < 0.05, ** p < 0.01, *** p < 0.001, ns = not significant). (**C**) Nucleotide diversity (π) along chromosome 17 (28 to 33 Mb) across the S-locus region, calculated in non-overlapping windows for each species. The S-locus is highlighted in blue. Points show individual window values; larger, more opaque points denote windows in the top 1% of π for that species (genome-wide threshold). Tick marks below each panel indicate the position of these top 1% windows. The dashed line shows the local 99th percentile of π across 50 bins spanning the region. Permutation test (10,000 permutations) results indicate whether top 1% windows are significantly enriched within the S-locus for each species.

Genetic diversity (*π*)^23,24^ computed in sliding windows peaked in the *S*-locus region in all four species, especially in *M. sylvestris* and *M. sieversii*, and was significantly enriched in the top 1% π windows (Fig. 2C, *p* < 0.05; Supplementary Fig. 10), supporting long-term balancing selection previously documented around plant self-incompatibility loci^25^.

Comparing across species, with different CNSR systems, the number of SFBB genes and families in *Malus* (∼18) was similar to that in *Petunia* (∼18 SLFs)^16^ but lower than in *Antirrhinum* (∼32 SLFs)^19^ (Table 1).

**Table 1:** Comparative architecture of the S-locus across flowering plants with CNSR SI systems. Summary of key genomic features of the S-locus across species with collaborative non-self recognition (CNSR)-based self-incompatibility. Shown are the number of pollen-expressed genes (SFBBs or SLFs), S-locus size (Mb), number of gene families (from sequence/function-based clustering), chromosomal position (when known), and references. Data are drawn from published studies and the present work (*Malus*). In species like *Petunia* and *Citrus*, the S-locus was inferred from BACs or partial assemblies.

| Model | Plant family | Number of pollen genes | Mean size of the S-locus (Mb) | Number of pollen gene families | Location | References |
| --- | --- | --- | --- | --- | --- | --- |
| Apple (*Malus*) | Rosaceae | 16–20 SFBBs | 1.35 | 18 | Peri-telomeric, long arm of chromosome 17 (17 chromosomes) | This study |
| Snapdragon (*Antirrhinum*) | Plantaginaceae | 32 SLF/SFB | 1.20 | 31 | Peri-telomeric, short arm of chromosome 8 (8 chromosomes) | <sup>19,26</sup> |
| Potatoes | Solanaceae | 12 SLFs | 13 | 8 | Chromosome 1 (12 chromosomes) | <sup>27</sup> |
| *Petunia* | Solanaceae | 16–20 SLFs | Unknown | 18 | Sub-centromeric, chromosome 3 (7 chromosomes) | <sup>16,28</sup> |
| *Coffea* | Rubiaceae | Unknown | Unknown | Unknown | Chromosome 5 (11 chromosomes) | <sup>29</sup> |
| *Citrus* | Rutaceae | 11–17 SLFs | 0.198 to 0.370 (BAC) | 12 | Unknown | <sup>17,30</sup> |
| *Pyrus* | Rosaceae | 17–20<br>SFBBs | 1.67 | 22 | Peri-telomeric,<br>long arm of<br>chromosome<br>17 (17 chro-<br>mosomes) | <sup>31</sup> |
| Pineapple<br>(*Ananas*) | Bromeliaceae<br>(Monocot) | Unknown | Unknown | Unknown | LG02 (25<br>chromosomes) | <sup>32</sup> |

Together, these results suggest that balancing selection preserves complete S-haplotype architectures rather than individual SI genes. Despite extensive sequence divergence, structural rearrangements, and accumulation of TEs, S-haplotypes sharing the same S-RNase maintain remarkably conserved SFBB repertoires and organization, indicating strong evolutionary constraints on functional compatibility.

### Multiple evolutionary processes generate SFBB diversity while preserving recognition modules

A major question regarding S-locus architecture is how new recognition specificities are generated and maintained by multiple pollen determinant genes and how they spread across S-haplotypes with different *S-RNase* alleles^9–11^. We obtained several lines of evidence suggesting that multiple evolutionary forces shape the diversity of *SFBB* genes within the *Malus* S-locus.

First, tandem and proximal duplications were found to be the dominant modes contributing to SFBB expansion, in contrast to dispersed duplications that dominated elsewhere on chromosome 17 (paired Wilcoxon signed-rank tests across haplotypes, n = 27; Tandem: p.adj < 0.0001; Proximal: p.adj < 0.001; Dispersed: p.adj < 0.0001, Fig. 3A). These local duplications likely provided the raw material for the functional diversification of S-alleles. Secondly, signatures of positive selection (dN/dS > 1, Supplementary Table 10 and 11) were detected in SFBBs, with CodeML identifying codons under strong positive selection (Fig. 3B). The proportion of positively selected codons was highest in the FBA domain, lowest (=0) in the F-box domain, and intermediate in regions outside both functional domains, a pattern consistent across nearly all haplotypes regardless of S-allele identity (Fig. 3B). Several of these positively selected sites occurred at conserved positions within a given SFBB family but varied between families, suggesting family-specific diversification (Table 2, Supplementary Table 12). These amino acids (aa) clustered in regions of predicted surface interaction based on Boltz-1 structural modeling^33^ (Supplementary Fig. 11) and exhibited high MuLAN local attention scores, suggesting functional relevance^34^ (Supplementary Fig. 12). Multivariate analysis of aa positions across families supported this pattern of divergence (Supplementary Fig. 13).

**Fig. 3:**
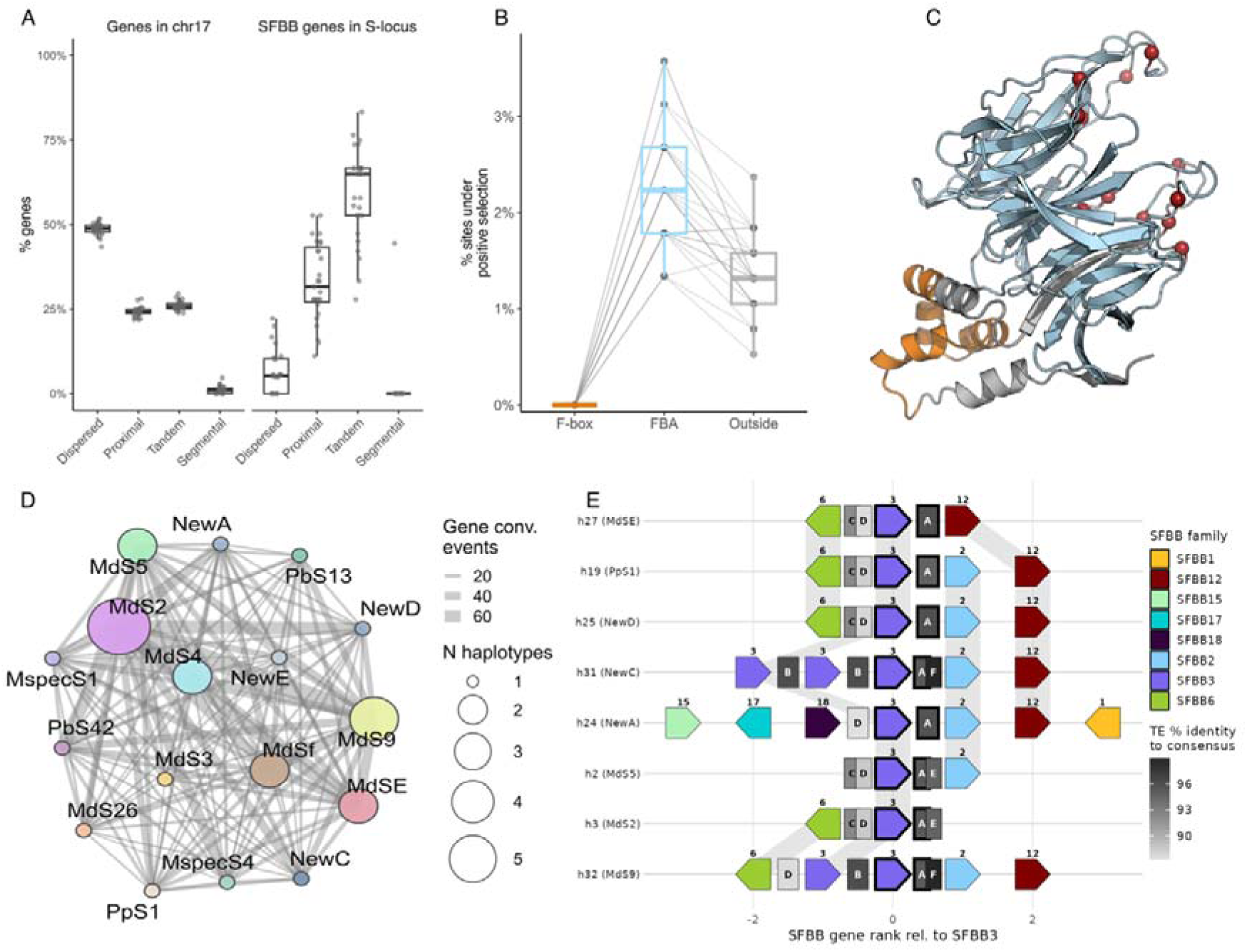
Duplication, selection, and sequence exchange generate SFBB diversity. (**A**) Proportions of genes classified by duplication mode (Dispersed, Proximal, Tandem, Segmental) in chromosome 17 as a whole versus within the S-locus, across all haplotypes. Boxes show the interquartile range and median; dots represent individual haplotypes. (**B**) Percentage of alignment sites under positive selection (BEB posterior probability > 0.99, codeml/PAML) within the F-box (yellow), FBA (blue), and Outside (grey)regions, with one point per haplotype. (**C**) Three-dimensional structural model of the SFBB2 protein. Sites under positive selection (BEB posterior probability > 0.99) are highlighted in red, as in (B); the FBA domain is in blue, and the F-box domain is in yellow. (**D**) Gene conversion network inferred with GENECONV across SFBB families. Nodes represent Salleles; node size is proportional to the number of haplotypes carrying that allele. Edge width is proportional to the number of gene conversion events detected between pairs of SFBB. Details are provided in Supplementary Table 16. (**E**) SFBB gene cluster synteny centered on SFBB3 across haplotypes with different S-alleles, ordered by S-RNase phylogeny. Arrows represent SFBB genes, coloured and numbered by family; SFBB3 (thick outline) serves as the anchor gene. Grey boxes indicate TE insertions within 15 kb of SFBB3; shading intensity reflects percentage identity to the TE consensus sequence. Letters code the TE consensus: A = MDOM_G18346 (significantly associated with SFBB3, permutation test *p* ≤ 0.05), B = MSYD_G11277, C = MSYL_G13944, D = MDOM_G14134, E = MSI2_G8771, F = de novo MDO_R5841 (B–F: local context only, not significant). Grey ribbons connect orthologous SFBB genes between adjacent haplotypes. The x-axis represents gene rank relative to SFBB3, not physical scale.

**Table 2:** Amino acid sites associated with SFBB gene families in Malus S-haplotypes. Summary of amino acid (aa) positions showing significant associations with SFBB gene families across 27 phased S-haplotypes. For each position, the number of aa variants across all families or within a given family, and the average number of families per aa are shown. p-values from multinomial regression are given with or without FDR correction. FBA: F-box–associated domain. Positively associated sites highlight candidates for SFBB diversification and allele-specific recognition. Full data in Supplementary Tables 12 and 13.

| Amino acid position | SFBB domain | Number of aa variants across families | Number of aa variants within families | Average number of families per aa | Multinomial regression |  |
| --- | --- | --- | --- | --- | --- | --- |
|  |  |  |  |  | p-value | Adjusted p-value |
| 297 | – | 10 | 1.33 | 2.40 | 1.1e–244 | 5.7e–244 |
| 299 | – | 10 | 1.44 | 2.60 | 3.0e–188 | 4.2e–188 |
| 336 | FBA | 11 | 2.22 | 3.64 | 5.0e–98 | 5.0e–98 |
| 356 | FBA | 7 | 1.50 | 3.86 | 4.9e–215 | 1.6e–214 |
| 358 | FBA | 11 | 1.28 | 2.09 | 2.2e–263 | 2.2e–262 |
| 421 | FBA | 11 | 1.83 | 3.00 | 9.0e–191 | 1.5e–190 |
| 429 | FBA | 10 | 1.39 | 2.50 | 3.5e–199 | 7.0e–199 |
| 476 | FBA | 7 | 1.39 | 3.57 | 3.8e–203 | 9.5e–203 |
| 543 | – | 12 | 2.00 | 3.00 | 8.1e–175 | 1.0e–174 |
| 549 | – | 8 | 1.61 | 3.63 | 3.6e–166 | 4.0e–166 |

In parallel, we observed strong sequence conservation within SFBB families despite their occurrence on divergent S-haplotypes (Supplementary Table 13), consistent with signatures of inter-haplotype gene conversion (Fig. 3D). A quasi-Poisson model identified allele pairs with significantly elevated conversion rates (MdS2–MspecS1, MdS2–MdSe, MdS2–MdSf), independent of their phylogenetic distance (Supplementary Fig. 14). Recurrent associations between specific SFBB families and nearby TEs provide an additional pattern consistent with sequence exchange among S-haplotypes (Fig. 3E, Supplementary Fig. 15 and 16, Supplementary Table 14). These SFBB–TE pairs were significantly more frequent than expected under random-association models (Supplementary Fig. 15, permutation test, p ≤ 0.01) and were observed across multiple S-haplotypes that carry distinct S-RNase alleles. The TE consensus sequences enriched for such associations do not belong to any specific classification (Supplementary Table 14), but all of these consensus sequences correspond to incomplete copies with a match in RepBase^35^ (Supplementary Table 15). Such conserved SFBB–TE cooccurrences, along with similar distances to TE consensus and similar insertion ages (Supplementary Fig. 16A–B), are consistent with sequence exchanges between haplotypes, potentially involving gene conversion events. Importantly, SFBB–TE associations remained correlated with specific S-RNase alleles, suggesting that only compatible combinations are retained by balancing selection.

### Candidate molecular determinants of recognition specificity

*S*-RNase polymorphisms drive SI specificity and show pronounced hypervariability (Supplementary Fig. 17). Codon-based analyses identified eight positively selected codons conserved within each *S*-RNase allele and distributed across the coding region, not confined to hypervariable segments. Several sites lie outside previously predicted recognition regions^36,37^, yet structural modeling indicated they were exposed on the protein surface (Fig. 4A; Supplementary Fig. 18). Their integrative prioritization, combining positive-selection scores and MuLAN per-residue attention scores, revealed that most S-RNase positively selected sites do not attain high attention values (Fig. 4C; Supplementary Fig. 19), suggesting that sequence-based models alone may not fully capture functionally relevant recognition sites on the pistil side.

**Fig. 4:**
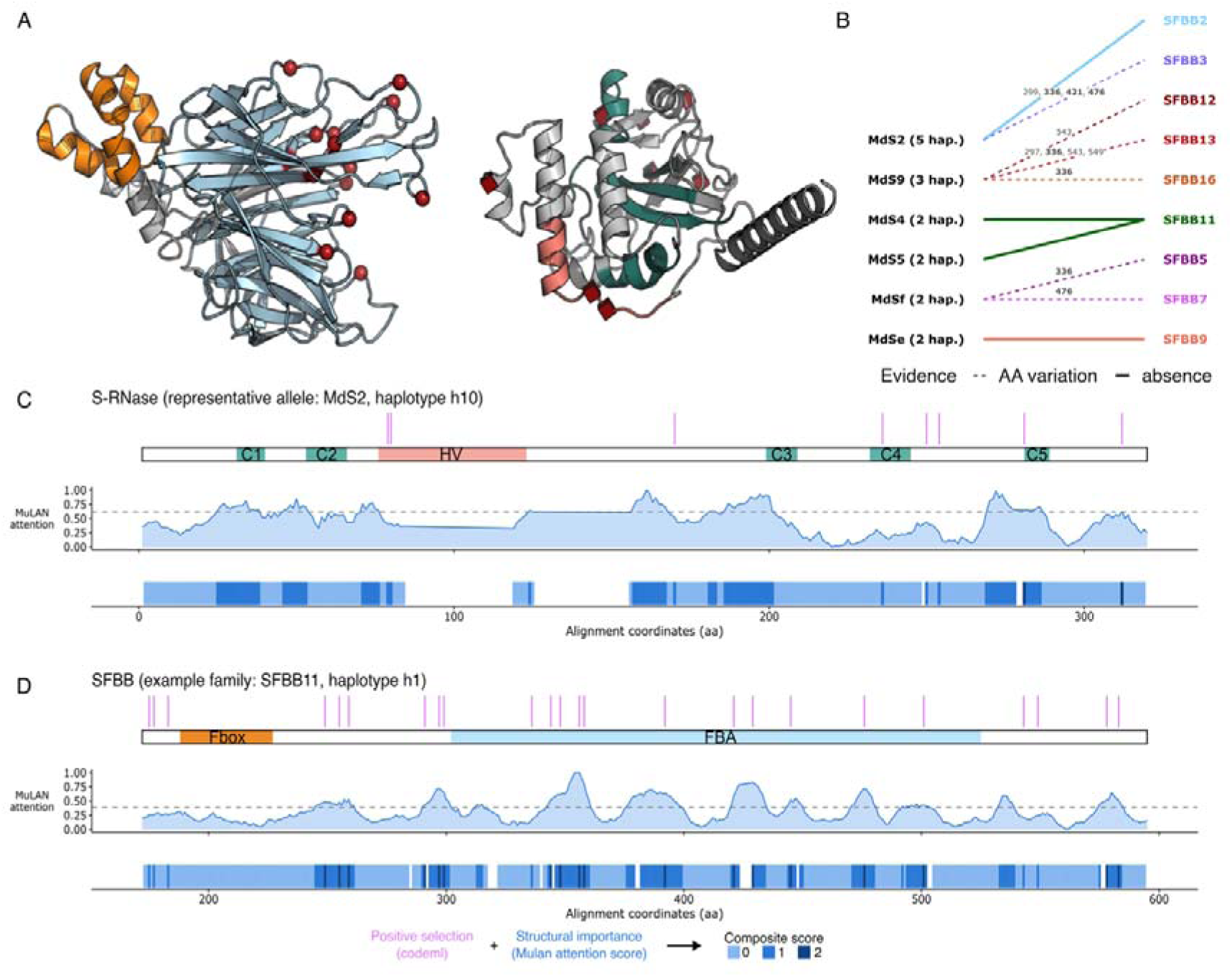
Candidate molecular determinants of recognition specificity. (**A**) Three-dimensional structural models of SFBB2 (left) and MdS2 S-RNase (right). In SFBB2, the F-box domain is shown in yellow and the FBA domain in light blue; in S-RNase, conserved regions C1– C5 are shown in dark green and the hypervariable region (HV) in salmon. Red spheres indicate positively selected sites identified by codeml (SFBB2) or by the intersection of FEL and codeml M2a/M8 analyses (S-RNase; Supplementary Fig. 17, 18). (**B**) Candidate detoxification relationships inferred from complete family absence across all haplotypes of an S-allele (solid lines) or consistent amino-acid variation at FBA-domain sites (dashed lines, bold site numbers). SFBB2 was absent from all MdS2 haplotypes; SFBB11 was absent from both MdS4 and MdS5 haplotypes, consistent with a role for these families in detoxification of the corresponding S-RNases (Supplementary Table 17). (**C**) Integrative prioritization of candidate recognition residues in S-RNase (representative allele MdS2, haplotype h10). From top to bottom: positive-selection sites (codeml, magenta) mapped onto the domain track (C1-C5, HV); MuLAN per-residue attention score (dashed line: top 30% per protein); composite score (0-2), summing the positive-selection and high-attention criteria at each position. Most S-RNase positive-selection sites do not reach high attention (Supplementary Fig. 19). (**D**) Same integrative analysis for a representative SFBB protein (family SFBB11, haplotype h1), with F-box and FBA domains indicated. Several SFBB11 sites combine positive selection and high attention (composite score = 2), concentrated in the FBA domain (Supplementary Fig. 12).

For SFBB proteins, positively selected codons were detected mainly in the FBA domain and interdomain regions (Fig. 3B), and selected residues mapped to predicted surface regions in structural models (Fig. 3C; Supplementary Fig. 11), with high MuLAN local attention scores at several sites (Fig. 4D; Supplementary Fig. 12).

To identify candidate detoxification relationships, we examined SFBB family presence– absence patterns across S-haplotypes sharing the same S-RNase (Fig. 4B). SFBB2 was absent from all MdS2 haplotypes (h14, h13, h3, h10, h23, Supplementary Table 17), SFBB11 was absent from both MdS4 (h20, h22) and MdS5 (h2, h26) haplotypes (Supplementary TableT17), consistent with the hypothesis that these families may participate in the detoxification of the corresponding S-RNases^18^. Family-specific amino acid substitutions at positively selected sites in SFBB genes further supported functional divergence among SFBB families—for example, SFBB12 in MdS2 haplotypes consistently showed a non-consensus residue at position 299 (Supplementary TableT12). SFBB18 was present in all haplotypes and lacked family-specific substitutions at selected sites, suggesting a more conserved function, or a role in detoxifying S-RNases not represented in the studied sample.

Despite these patterns, structural predictions alone did not reliably distinguish interacting from non-interacting SFBB–S-RNase pairs. Boltz-1 generated generally confident models for individual proteins (Supplementary Fig. 20), but confidence scores for predicted complexes showed extensive overlap among the expected, unsupported, and unknown interaction classes (Supplementary Fig. 21). Applying the same framework to Petunia, where several SLF–S-RNase interactions have been experimentally validated, yielded higher overall confidence scores and lower variance among predictions (Supplementary Fig. 20–25), yet interaction classes remained largely overlapping.

These results suggest that structural prediction tools currently capture general folding properties of SI proteins but remain insufficient to reliably infer recognition specificity in CNSR systems.

### Pollen-specific lncRNAs as candidate accessory components of the S-locus

Beyond S-RNase and SFBB, we examined whether the S-locus harbors additional candidate regulators by profiling gene expression across leaf, pistil, and pollen tissues in three reference haplotypes (Gala haplotype 1, Gala haplotype 2, and Honeycrisp). S-RNase genes showed pistil-biased expression, whereas SFBB genes were predominantly expressed in pollen, consistent with their expected roles in the SI response (Supplementary Fig. 27, Supplementary Table 19). Among 83 S-locus lncRNAs expressed in pollen, six showed strict pollen-specific expression and were absent from both pistil and leaf tissues (Supplementary Table 18). These transcripts were highly expressed (TPM = 284– 7,215) and distributed throughout the S-locus region (Supplementary Fig. 28), identifying them as candidate reproductive lncRNAs that may act as accessory components of the S-locus. Structural cofolding predictions incorporating these pollen-specific lncRNAs into S-RNase–SFBB complexes did not provide strong evidence of stable direct interactions, although one lncRNA consistently increased predicted interface confidence for the MdS2/SFBB2 pair (Supplementary Fig. 29). Full methods and results for this analysis are reported in Supplementary Note 2.

## Discussion

### Long-term balancing selection preserves complete S-haplotype architectures despite extensive haplotype divergence

The most striking result of our study is that balancing selection preserves complete S-haplotype architectures rather than individual self-incompatibility determinants. Haplotypes carrying the same S-RNase allele retained highly similar SFBB repertoires and gene organization, even across different *Malus* species, despite extensive structural divergence among alternative haplotypes. This finding extends the classical view of trans-species S-allele persistence under balancing selection^20,21^ by showing that long-term maintenance also applies to multigenic S-haplotype architectures. Such conservation, also reported for multiple copies of the S17 haplotype of *Pyrus*^31^ is particularly remarkable given the dynamic genomic environment of the S-locus, characterized by structural rearrangements, transposable element accumulation, and ongoing gene turnover. Together, these results confirm predictions that functional compatibility should impose strong constraints on S-haplotype evolution in CNSR systems^9–11^, allowing diversification of individual recognition genes over long evolutionary time^20^ while maintaining stable recognition architectures over evolutionary timescales^21^.

### A structurally dynamic locus shaped by restricted recombination

The apple S-locus shares several genomic hallmarks with other regions of long-term restricted recombination, including tight linkage among functionally interacting genes, extensive structural divergence among alternative haplotypes, and long-term balancing selection maintaining distinct haplotypic configurations^12–15^. The high density of transposable elements further suggests long-term restricted effective recombination. These features are consistent with theoretical expectations for regions of suppressed recombination^12–14^ and indicate that the S-locus of Rosaceae experiences many of the same evolutionary constraints. Moreover, the accumulation of TEs suggests reduced efficacy of purifying selection due to Hill–Robertson interference^38^, a process commonly associated with recombination suppression in such regions^12,13^. Yet unlike some regions of restricted recombination that progressively degenerate, the S-locus must simultaneously preserve functional diversity across multiple genes. The coexistence of mutational accumulation and strong balancing selection highlights an inherent evolutionary tension between maintaining long-term polymorphism and limiting genomic deterioration. Understanding how SI systems resolve this tension may provide broader insights into the evolution of other complex recognition systems maintained by balancing selection.

### Evolutionary maintenance of complete recognition repertoires

A central prediction of theoretical models of CNSR evolution is that selection should favor haplotypes carrying nearly complete repertoires of pollen determinants capable of detoxifying all non-self S-RNases^9–11^. Our findings provide empirical support for this prediction. Across the 27 phased Malus S-haplotypes, individual haplotypes carried representatives of 15 to 18 recognized SFBB families (mean = 16.7), missing at most three families relative to the full genus-wide repertoire. Strikingly, five haplotypes (h6, h7, h11, h15, h30) appeared to carry a member of all 18 families. We interpret this as evidence that the self-recognizing SFBB is not necessarily absent from the genome, but may instead persist as a degraded, pseudogenized copy retaining enough sequence similarity to be classified within its family despite being non-functional. A directly comparable pattern was experimentally validated in *Petunia*, where each S-haplotype carries 16–20 of the SLFs required to recognize the vast majority of non-self S-RNases, and haplotypes were shown by transformation assays to specifically lack the SLF conferring recognition of their own S-RNase^16^. In *Antirrhinum*, haplotypes similarly carry a large repertoire of pollen determinants (32 SLFs per haplotype, clustering into 31 types across sampled haplotypes)^19^, although whether this reflects the same one-family-missing pattern has not been directly assessed. Together, these observations indicate that haplotypes across CNSR systems typically encode pollen determinants against nearly the entire non-self recognition spectrum, consistent with the strict self/non-self discrimination required for CNSR.

The number of SFBB families we identified in *Malus* (18) differs somewhat from the 22 groups recently defined by Gu et al. (2025)^31^ across combined *Pyrus* and *Malus* sequences, likely reflecting differences in taxonomic scope and clustering approach rather than a true disagreement on SFBB diversity.

At the same time, complete recognition does not necessarily require a strict one-to-one relationship between pollen determinants and S-RNases. The broader recognition patterns observed for SFBB11 suggest that some SFBBs may recognize multiple S-RNases, consistent with theoretical expectations that promiscuous recognition can facilitate the emergence and maintenance of new Shaplotypes^11^. Such flexibility may explain how more than 30 S-alleles can be maintained in *M. domestica*^39^ despite the presence of only ∼18 SFBB families per haplotype. Together, these observations suggest that balancing selection optimizes recognition repertoires through a trade-off among specificity, redundancy, and the genomic constraints imposed by restricted recombination^9–11^.

### Diversification of pollen determinants through duplication and sequence exchange

Maintaining complete recognition repertoires requires a continual supply of novel pollen specificities^9–11^. Our analyses suggest that diversification and conservation operate simultaneously within the *Malus* S-locus. SFBB expansion was primarily driven by tandem and proximal duplications, in contrast to the duplication patterns observed elsewhere in chromosome 17. These local duplications likely provide the raw material for neofunctionalization and the emergence of novel recognition functions.

Positive selection acting predominantly on the FBA domain further indicates adaptive diversification of substrate recognition surfaces. At the same time, the high sequence similarity observed among homologous SFBB families across divergent S-haplotypes, together with signatures consistent with gene conversion, suggests that sequence exchange contributes to the spread of functional variants. Rather than representing opposing evolutionary forces, duplication and sequence exchange may jointly promote innovation while preserving compatibility with resident S-RNases. This dynamic could help reconcile the paradox of CNSR evolution: generating new recognition specificities without disrupting existing mating compatibility.

### Beyond canonical SI determinants: candidate accessory regulators and limits of current interaction models

In addition to the canonical S-RNase and SFBB determinants, the pollen-specific lncRNAs identified within the S-locus (Supplementary Note 3) represent candidate accessory regulators of the SI response, although their biological role remains unknown. Similar regulatory layers have been implicated in reproductive signaling pathways in other plant systems^8^, raising the possibility that the molecular architecture of the S-locus extends beyond the classical toxin–antitoxin framework.

Our structural analyses further illustrate the current limitations of computational approaches for deciphering CNSR specificity. Although Boltz-1 and MuLAN successfully captured general structural features of SI proteins, they failed to reliably distinguish expected from unsupported SFBB–S-RNase interactions. These findings suggest that current AI-based structural prediction methods remain insufficient to infer recognition specificity in CNSR systems. Future experimental approaches— including protein interaction assays, CRISPR-mediated perturbations, and functional complementation experiments—will be necessary to establish the molecular basis of recognition and determine whether accessory transcripts contribute to S-locus function.

### Outlook

Our results reveal how multigenic recognition systems can remain evolutionarily stable despite continual turnover of their individual components. By combining haplotype-resolved genomics, evolutionary analyses, and structural predictions, we show that innovation and conservation coexist within the apple S-locus: new SFBB variants arise through duplication, selection, and sequence exchange^9–11,16,17^, whereas balancing selection preserves higher-order recognition architectures^1,20,21^. More broadly, these findings suggest that balancing selection may act not only on individual alleles but also on complex multigenic modules, providing a general framework for understanding the longterm maintenance of adaptive polymorphisms across biological systems^1,21^.

Taken together, our results show that the apple S-locus combines evolutionary stability at the haplotype level with continual turnover of individual recognition genes.

## Methods

### Dataset description

Twenty-seven assembled haplotypes (Supplementary Table 1) from *Malus* species, including both cultivated apple (*M. domestica*, 16 genotypes, comprising 14 dessert varieties and two cider varieties) and wild relatives (11 genotypes from *M. sylvestris*, one *M. sieversii*, and two *M. orientalis*) were examined in this study. Among the individuals used to produce this dataset, 17 were diploids. In most cases, two haplotype assemblies were obtained per individual (Gala and Fuji cultivars, the cider individual, the *M. orientalis* individual, and the four *M. sylvestris* individuals grown at IDEEV). For the Red Delicious, Honeycrisp, Brown Snout, Costard, Bardsey Island, and Newton cultivars, and for *M. sieversii* and *M. sylvestris*, only one haplotype was kept from previous studies due to the unavailability of the second haplotype or insufficient assembly quality. For Golden Delicious and Hanfu, the haplotypes were derived from a double haploid individual, the GDHH13 *M. domestica* reference genome^40^, and a tri-haploid individual^41^. This dataset combines 15 already available haplotype assemblies and 12 new sequenced haplotypes from which only chromosome 17 was retained for this study, as this chromosome hosts the SI locus in *Malus*^42^ (Fig. 1).

The 27 haplotypes displayed high assembly completeness, with genome sizes ranging from ∼600 Mb to 660 Mb, scaffold N50 values of ∼9–39 Mb (average N50 = 33.09 Mb ± 6.86), and BUSCO^43^ completeness scores generally > 95%. Repetitive content varied moderately between haplotypes (65–76%). Details about the genome assemblies and gene annotations are provided in Supplementary Table 1 and Supplementary Note 1. For convenience, all haplotypes were identified with an ID from h1 to h32. To avoid bias due to heterogeneous annotations, all haplotypes were reannotated using a single, standardized gene annotation pipeline.

### Construction of an S-RNase and SFBB gene database

An S-locus gene database, consisting of all *S-RNase* and *SFBB* sequences available in public repositories, was built as a raw material to investigate the genomic architecture of these genes in the genome assemblies described above. Public *S-RNase* DNA sequences^44–47^ and *SFBB* DNA sequences^4,48^ were collected. The homologous sequences of the *S-RNase* and *SFBB* genes were then recovered using BLASTN searches on the GenBank database with the parameter –e-value 0.001 and added to complete the S-locus gene database. The *S-RNase* database was split between exon 1 and 2 sequences to avoid mis-mapping due to high polymorphism of the intron between individuals. The accession numbers of the genes are provided in Supplementary Tables 2 and 6.

### S-RNase assignment of each haplotype

Using the database of publicly available *S-RNase* sequences constructed above (Supplementary Table 2), the Maloideae *S-RNase* sequences in the database were aligned to *S-RNase* sequences from the haplotypes using Clustal Omega v1.2.4^49^ to assign a given *S-RNase* allele to each haplotype (Supplementary Table 3 and Supplementary Fig. 1).

### Haplotype assembly and gene annotation

Twenty-seven haplotype assemblies sourced from publicly available data and newly sequenced genotypes were constructed using PacBio HiFi sequences. Each haplotype was defined as a set of chromosomes transmitted together (Supplementary Table 1 and Supplementary Note 2).

Among these, 15 haplotype assemblies were obtained from the literature. Twelve haplotype assemblies were from 10 *M. domestica* cultivars: double haploid Golden Delicious (one haplotype available, GDDH13), Fuji (two haplotypes), Gala (two haplotypes), Red Delicious (one haplotype available), Honeycrisp (one haplotype), tri-haploid Hanfu (one haplotype available), Brown Snout (one haplotype available), Costard, Bardsey Island (one haplotype available), and Newtons (one haplotype available). Additionally, three haplotype assemblies were used, with one from *M. sieversii* and two from *M. sylvestris*.

The dataset was enriched with six newly sequenced genomes, representing 12 haplotypes. These new genomes represented one M. domestica genotype (Xuanina, used for cider production), one *M. orientalis* genotype from Armenia, and four *M. sylvestris* samples, one each from Austria, Denmark, France, and Romania. Each newly sequenced genome contributes two parental haplotypes, resulting in a total of 12 supplementary haplotypes. Details about DNA extraction, sequencing, and assembly of the seven newly sequenced genomes are summarized in Supplementary Table 1 and Supplementary Note 2. Briefly, PacBio sequencing and assembly of the resulting reads generated an average assembly size of 642 Mb, similar to publicly available assemblies (average of 636.5 Mb), with an average scaffold N50 value of 28.12 Mb, which was lower than the average scaffold N50 of publicly available assemblies (average of 37 Mb).

### Characterization of SFBB gene families

To cluster SFBB genes from all haplotypes into families, all complete SFBB protein sequences were aligned using Clustal Omega v1.2.4^49^. The alignment was used as input for IQ-tree^50^ with ModelFinder^51^ for a first run to determine the best-fitting substitution model using 1,000 bootstrap replicates. Using the resulting phylogenetic tree, the hierarchical clustering method of Ward^52^ was used with the cophenetic distance matrix, using R^53^ and the software packages ape v5.7-1^54^, FactoClass v1.2.8^55^, and stats v4.3.1^53^. Additionally, OrthoFinder^56^ was run on SFBB protein sequence alignments derived from each haplotype. To validate the defined SFBB families, identity percentages were computed from pairwise SFBB alignments (from the same families or different families) using the pairwiseAlignment command from the R package Biostrings v2.68.1^57^.

The Malus S-locus is located in the peri-telomeric region on the long arm of chromosome 17^42^. Therefore, chromosome 17 contigs and annotated genes were extracted using the Helixer/Eggnog pipeline (Supplementary Note 2), followed by analysis of the SI regions to define their size and gene content.

### Phylogenetic analysis of S-RNases

To determine the haplotypes of the S-alleles, S-RNase protein sequences were aligned to reference sequences listed in Supplementary Table 2 using Clustal Omega v1.2.4^49^. S-RNase sequences associated with S-alleles from previous studies were used to identify S-alleles in the S-haplotype dataset. Specifically, phylogenetic relationships among all S-RNase sequences were examined using IQ-tree^50^ with ModelFinder^51^ in the first run to determine the best-fitting substitution model using 1,000 bootstrap replicates.

## Sequences and gene synteny within S-loci

To examine the synteny across the S-loci from the 27 Malus haplotypes, regions encompassing 500 kb upstream and downstream of S-locus sequences were aligned using the nucmer tool in MUMmer^58^. SyRI (Synteny and Rearrangement Identifier) v1.6.3^59^ was then used to identify the syntenic path (longest set of collinear regions), structural rearrangements (inversions, translocations, and duplications), local variations (single nucleotide polymorphisms [SNPs], insertion/deletions [InDels], copy-number variation [CNV]) within syntenic blocks and structural rearrangements, and unaligned regions. SyRI takes alignment blocks identified by the initial aligner and identifies larger regions consisting of multiple consecutive one-to-one alignment blocks. Finally, PlotSR^60^ was used to visualize synteny using the output from SyRI.

To generate gene synteny plots, S-loci annotation gff files from Apollo^61^ were converted to bed files using agat_convert_sp_gff2bed.pl from AGAT^62^, which were used as input for the R package gggenomes v0.9.9.9000^63^. Identity between S-locus genes was computed using Diamond^64^ based on their corresponding amino acid (aa) sequences.

Syntenic relationships across several S-loci were examined using the network-based approach with the R package Syntenet v1.9.1^22^, which treats anchor pairs (duplicates retained from a large-scale duplication event) from synteny comparisons as connected nodes of an undirected unweighted graph^65^.

### Nucleotide diversity analysis with Pixy

Nucleotide diversity (π) across the genome was calculated using Pixy v1.2.6.beta1 software^23^, which accurately estimates π from VCF files that include invariant sites and accounts for missing data. π was computed in non-overlapping 10-kb windows using the species information for each sample in the dataset.

The input VCF file was originally generated in Chen et al., (2026)^66^, and includes multiple phased haploid genomes from several Malus species. From this dataset, a subset of 41 samples was extracted, representing four species relevant to our study: M. domestica (n=10), M. orientalis (n=9), *M. sieversii* (n=10), and *M. sylvestris* (n=12). The analysis was restricted to chromosome 17, which harbors the S-locus. The coordinates of the S-locus in the GDDH13 reference genome^40^, corresponding to haplotype h1 in our study, were defined as 30,032,275 bp and 31,732,190 bp, and windows were classified as either inside or outside the S-locus.

To identify outlier windows with significantly higher π than the remaining genomes, the windows with the top 1% highest-π values within each species were defined independently. To determine whether the S-locus was enriched for high-diversity windows, a permutation-based enrichment test was performed using R^53^. The empirical *p*-value was calculated as the proportion of permutations in which the number of outlier windows in the S-locus was equal to or greater than the observed count. A significant *p*-value (*p* < 0.05) indicates that high-diversity windows are non-randomly concentrated in the S-locus, consistent with a signal of balancing selection.

### Signatures of selection at SI genes

To determine whether the *SFBB* genes evolved neutrally or by natural selection (positive or purifying), the codeml program in the PAML package v4.9^71^ was used. Codeml computes the ratio of nonsynonymous (dN) to synonymous (dS) substitutions for various models of sequence evolution. The most likely model explaining the data is chosen based on a likelihood ratio test computed among models. Three pairs of site-specific models allowed us to test for recurrent, diversifying selection (M0 and M3, M1a and M2a, and M7 and M8). One SFBB protein sequence alignment for each haplotype was used. To assess whether aa identity at each aligned position was significantly associated with the *SFBB* gene family, a multinomial regression model was fitted for each position that showed at least two different aa and was present in at least two families. For each position, a null model (aa ∼ 1) was compared to a full model (aa ∼ Family) using a likelihood ratio test. *P*-values were computed from the χ^2^ distribution and adjusted for multiple testing using the Benjamini–Hochberg procedure to control the false discovery rate (FDR). Multiple correspondence analysis (MCA), which is specifically designed for categorical data and better captures variation in aa composition across discrete sites, was performed to investigate the clustering of *SFBB* genes based on aa identity at key positions. For each gene, aa residues at conserved positions were extracted and treated as categorical variables. The MCA was conducted using the *FactoMineR* package^72^ in R^53^, with the SFBB family provided as a supplementary qualitative variable. Individuals (genes) were projected onto the first two dimensions, and family labels were used for colouring.

For S-RNases, 27 protein sequences were aligned using Clustal Omega v1.2.4^49^. To identify codon positions likely involved in allele-specific recognition in the S-RNase sequences, evidence from multiple complementary approaches was combined. First, the Fixed Effects Likelihood (FEL) method available on the Datamonkey web platform^73^ was applied for codon alignment restricted to one representative sequence per S-allele to focus on substitutions that occurred between allelic groups rather than within them. In parallel, the site models implemented in codeml (PAML^71^) were used to detect codon sites evolving under positive selection based on likelihood ratio tests comparing models, allowing for *ω* > 1 (e.g., M2a, M8) against their respective null models. The results from both methods were compared to retain only robust candidates. To assess whether these positively selected sites were conserved within allelic groups, the full codon alignment was examined, including all available sequences per allele. For each candidate site, intra-allelic variation was quantified by counting the number of distinct codons observed within each group. Positions showing no variation within any allele, i.e., monomorphic within-group but polymorphic between groups, were considered strong candidates for functional divergence related to allele-specific interactions.

### Search for duplicated genes

To examine how multiple *SFBB* genes were generated and proliferated, the classify_gene_pairs command in the R/Bioconductor package *doubletrouble*^74^ was used with the standard scheme (binary = FALSE). In short, intra-haplotypes protein similarity searches were performed using DIAMOND^64^ (sensitive mode, e-value = 1e−10, top hits = 5) to identify the entire set of paralogous gene pairs for each haplotype. Paralogous gene pairs in the same syntenic blocks were classified as arising from segmental duplication. A tandem duplication is a segmental duplication that results from an unequal crossing over^75,76^. A few genes separate proximal gene pairs^76^ and can originate from ancient tandem duplication events disrupted by the insertion of other genes^77^. Dispersed duplication occurs through unpredictable and random patterns by mechanisms that remain unclear, generating two gene copies that are neither neighboring nor colinear^78^.

### Signatures of gene conversion in SFBB genes

Intra-*SFBB* family gene conversion signatures were detected using GENECONV^79^, which scans aligned sequences and identifies segments (tracts) that are (1) identical or nearly identical, (2) longer than expected by chance, and (3) shared between sequences that are otherwise divergent. GENECONV tests gene conversion using 10,000 permutations and assesses significant gene alignment using Bonferroni correction as well as the Karlin-Altschul score. To identify potential preferential gene conversion between specific S-allele pairs, a global table of gene conversion events was compiled, with each event associated with the S-alleles of the two haplotypes involved. For each event, the corresponding S-allele pair was annotated. Because S-alleles were unevenly represented across the 27 S-haplotypes, each S-allele was assigned a weight equal to the inverse of its frequency across the set of assemblies. To enable robust statistical testing, a bootstrap procedure was implemented. In each replicate, 10 S-haplotypes were randomly selected, and only gene conversion events between S-alleles present in that subset were retained. The number of events per S-allele pair was then counted and associated with the corresponding mean weight. After repeating this procedure 500 times, a dataset was obtained with multiple observations per pair. A quasi-Poisson regression model using the glm function from the R package *stats*^53^ was then used to test whether some S-allele pairs exhibited significantly higher or lower gene conversion frequencies than expected. The model was fitted using the number of conversion events as the response variable, the S-allele pair as a predictor, and the mean weight as a statistical weight to account for allelic representation bias.

### TE distribution and enrichment analysis around SFBB genes

TEs within the S-locus were annotated using a pan-genomic TE library built from a representative set of apple genomes. The TE consensus libraries were first generated using the TEdenovo pipeline from REPET v3.0 using Tedenovo^80^, and then using these libraries the TEs of the following genomes were annotated using TEannot^81^: the reference cultivar GDDH13^40^, one haplotype from a cider variety (unpublished data), one haplotype from each European Malus sylvestris population (unpublished data), one haplotype of M. orientalis (unpublished data), and one haplotype of M. sieversii^82^. These libraries were concatenated, and redundancy was removed using PASTEC^83^ with thresholds of at least 98% identity and 95% coverage. Sequences showing high similarity to SFBB genes (>90%) or classified as potential host genes (PHGs) by PASTEC were excluded. The resulting curated database was then used to determine shared and non-shared TE insertions across genomes in S-locus regions using panREPET^84^.

To identify potentially recurrent associations between TEs and *SFBB* gene families independently of the *S-RNase* allele, a permutation test was performed. For each TE consensus–*SFBB* family pair, the number of distinct S-alleles (nS-alleles) observed among the haplotypes where the association occurred was counted. TE–*SFBB* proximity was restricted to ≤ 4 kb as a conservative threshold. Inter-*SFBB* distances within the S-locus are generally orders of magnitude larger (commonly tens of kb; Supplementary Fig. 30), so a 4-kb window targets elements that are genuinely colocated with an *SFBB* gene, encompassing likely regulatory flanks and the range of plausible short recombination/gene-conversion footprints while limiting noise from distant TEs. This number was then compared to a null distribution obtained by randomly permuting the S-alleles across haplotypes. At each iteration, S-alleles were shuffled across haplotypes while preserving the TE–*SFBB* associations. An empirical *p*-value was calculated as the proportion of permutations in which the number of S-alleles was equal to or greater than the observed value. This approach allows us to detect TE–*SFBB* associations that recur across multiple S-alleles more frequently than expected by chance, suggesting possible gene conversion or recombination events within the S-locus. These associations were visualized as heatmaps using the R package *ggplot2*^85^.

### Protein–protein interactions

The sequences from the different families were analysed using MuLAN^34^, a sequence-based deep-learning method designed to predict changes in binding affinity induced by mutations affecting protein sequences. In addition, MuLAN can be run on single sequences to compute scores for each residue based on a local attention mechanism, normalized in the range [0, 1], which correlates with interacting or functional regions. In this work, we applied MuLAN to single sequences. Finally, the sequences in each family were aligned. The score profiles at the aligned positions were plotted, highlighting the experimentally determined positive selection sites (Supplementary Fig. 12, Supplementary Fig. 19, and Supplementary Fig. 25). Predicted structures for protein complexes were obtained with Boltz-1^33^, using the 1.0.0 version of the package. Structures were rendered in PyMOL (v3.1.6.1)^86^. For each complex, multiple sequence alignments were automatically retrieved using the MMseqs2 server, and five alternative structures were generated, setting an initial random seed of 42. For each case, we then selected the predicted structure with the highest associated confidence score. We did not employ additional steering potentials as no steric clashes were observed in the model’s outputs.

In the comparison of model structures of complexes generated from *Malus* and *Petunia* sequences, we evaluated structural differences with different confidence measures of the predicted protein–protein interaction: pLDDT (predicted local distance difference test) and i-pLDDT (interface pLDDT), which measure local confidence averaged over residues, and pTM (predicted template modeling score) and i-pTM (interface pTM), which measure overall accuracy of the structure and relative orientation of the subunits.

### lncRNA Identification, and S-Locus Expression Analysis

RNA-seq analyses were performed using a modular Bash/R workflow on three haplotype-resolved apple references (h2 and h3 from Gala cultivar, and h10 from Honeycrisp cultivar). Reads were quality-filtered and trimmed with fastp^87^, aligned to haplotype-specific genomes using STAR^88^, and processed with samtools^89^. Per-sample transcript assemblies were generated with StringTie^90^ and merged at the reference level, followed by transcript classification with gffcompare^91^. Candidate lncRNAs were selected from non-reference transcript classes, structurally filtered, evaluated for coding potential using CPC2^92^, CPAT v3.0.5^93^, PLEK2 v2.1^94^ and RNAplonc v1.0^95^, and further filtered using Pfam-domain screening to retain high-confidence noncoding transcripts.

For expression quantification, SALMON^96^ was used in two complementary modes: (i) a global transcriptome mode based on combined mRNA + lncRNA transcript references, generating lncRNA transcript-level TPM matrices and global gene-level TPM matrices; and (ii) an S-locus-targeted mode based on restricted transcriptomes containing S-locus genes and lncRNAs. S-locus features were classified as SI_gene (SFBB and S-RNase), other_gene, or lncRNA.

S-locus lncRNA sequences were clustered across haplotypes using CD-HIT-EST^97^ to define sequence-based lncRNA clusters and cluster membership. Differential expression analyses were performed with DESeq2^98^ across tissue contrasts. Significantly DE lncRNAs were further annotated with genomic coordinates, lncRNA cluster membership, and proximity/overlap relative to reference-specific SFBB and S-RNase BED annotations, including nearest SFBB family assignment. Custom R scripts were used to generate S-locus heatmaps and summary tables, including log2 fold-change summaries for S-locus coding genes annotated as SFBB, S-RNase, or other genes.

## Data availability

Chromosome 17 data are available in the PRJNA1511955 Bioproject. The coding sequences and protein sequences of SFBB and S-RNase genes are available in the ZENODO repository: DOI https://doi.org/10.5281/zenodo.17226708. Source data are provided with this paper.

## Code availability

All analysis scripts and custom code are available at https://github.com/CornilleEclecticLab/Apple-SI/tree/main.

## Acknowledgements

The authors thank INRA-CNRGV (http://cnrgv.toulouse.inra.fr/) for HMW DNA extraction and longread sequencing. We are grateful to the Genotoul bioinformatics platform Toulouse Occitanie (Bioinfo Genotoul, https://bioinfo.genotoul.fr) and to NYUAD Jubail High Performance Computing (HPC) for providing help, computing, and/or storage resources. This work was carried out using the URGI facilities (https://doi.org/10.15454/1.5572414581735654E12) and was supported by the French government through the National Research Agency (ANR), as part of the France 2030 program for research infrastructure (EQUIPEX+), reference ANR-21-ESRE-0048. We thank J. Flowers, K. Alix and T. Giraud for insightful discussions.

## Author contributions

AMC, VC, and XV designed the study. AMC obtained funding. AV, WM, XC, AM, and AMC generated, assembled, and anchored the genome sequences. AM, GL, SS, JC, ALF, AC, and AMC analyzed the data. All co-authors discussed the results. AM, AMC, VC, MM, XV, and AC wrote the manuscript with critical input from other co-authors.

## References

1. Charlesworth, D. Balancing Selection and Its Effects on Sequences in Nearby Genome Regions. PLoS Genet. 2, e64 (2006).

2. Glémin, S. & Galtier, N. Genome Evolution in Outcrossing Versus Selfing Versus Asexual Species. in Evolutionary Genomics (ed. Anisimova, M..) vol. 855 311–335 (Humana Press, Totowa, NJ, 2012).

3. Wright, S. I., Ness, R. W., Foxe, J. P. & Barrett, S. C. H. Genomic Consequences of Outcrossing and Selfing in Plants. Int. J. Plant Sci. 169, 105–118 (2008).

4. Minamikawa, M. et al. Apple S locus region represents a large cluster of related, polymorphic and pollen-specific F-box genes. Plant Mol. Biol. 74, 143–154 (2010).

5. Sassa, H., et al. *S Locus F-Box Brothers*: Multiple and Pollen-Specific F-Box Genes With *S* Haplotype-Specific Polymorphisms in Apple and Japanese Pear. Genetics 175, 1869–1881 (2007).

6. Fujii, S., Kubo, K. & Takayama, S. Non-self– and self-recognition models in plant self-incompatibility. Nat. Plants 2, 16130 (2016).

7. Williams, J. S., Wu, L., Li, S., Sun, P. & Kao, T.-H. Insight into S-RNase-based self-incompatibility in Petunia: recent findings and future directions. Front. Plant Sci. 6, (2015).

8. Du, J. et al. Molecular characteristics of *S-RNase* alleles as the determinant of self-incompatibility in the style of *Fragaria viridis*. Hortic. Res. 8, (2021).

9. Bod’ová, K., Priklopil, T., Field, D. L., Barton, N. H. & Pickup, M. Evolutionary Pathways for the Generation of New Self-Incompatibility Haplotypes in a Nonself-Recognition System. Genetics 209, 861–883 (2018).

10. Harkness, A., Goldberg, E. E. & Brandvain, Y. Diversification or Collapse of Self-Incompatibility Haplotypes as a Rescue Process. Am. Nat. 197, E89–E109 (2021).

11. Erez, K., Jangid, A., Feldheim, O. N. & Friedlander, T. The role of promiscuous molecular recognition in the evolution of RNase-based self-incompatibility in plants. Nat. Commun. 15, 4864 (2024).

12. Jay, P. et al. Supergene Evolution Triggered by the Introgression of a Chromosomal Inversion. Curr. Biol. 28, 1839–1845.e3 (2018).

13. Jay, P., Jeffries, D., Hartmann, F. E., Véber, A. & Giraud, T. Why do sex chromosomes progressively lose recombination? Trends Genet. 40, 564–579 (2024).

14. Kay, T., Helleu, Q. & Keller, L. Iterative evolution of supergene-based social polymorphism in ants. Philos. Trans. R. Soc. B Biol. Sci. 377, 20210196 (2022).

15. Goubet, P. M. et al. Contrasted Patterns of Molecular Evolution in Dominant and Recessive Self-Incompatibility Haplotypes in Arabidopsis. PLoS Genet. 8, e1002495 (2012).

16. Kubo, K. et al. Gene duplication and genetic exchange drive the evolution of S-RNase-based self-incompatibility in Petunia. Nat. Plants 1, 14005 (2015).

17. Liang, M. et al. Evolution of self-compatibility by a mutant Sm-RNase in citrus. Nat. Plants 6, 131–142 (2020).

18. Kubo, K. et al. Collaborative Non-Self Recognition System in S-RNase–Based Self-Incompatibility. Science 330, 796–799 (2010).

19. Zhu, S. et al. The Snapdragon Genomes Reveal the Evolutionary Dynamics of the *S* –Locus Supergene. Mol. Biol. Evol. 40, msad080 (2023).

20. Vekemans, X. & Slatkin, M. Gene and allelic genealogies at a gametophytic self-incompatibility locus. Genetics 137, 1157–1165 (1994).

21. Castric, V., Bechsgaard, J., Schierup, M. H. & Vekemans, X. Repeated Adaptive Introgression at a Gene under Multiallelic Balancing Selection. PLoS Genet. 4, e1000168 (2008).

22. Almeida-Silva, F., Zhao, T., Ullrich, K. K., Schranz, M. E. & Van De Peer, Y. syntenet: an R/Bioconductor package for the inference and analysis of synteny networks. Bioinformatics 39, (2023).

23. Korunes, K. L. & Samuk, K. PIXY: Unbiased estimation of nucleotide diversity and divergence in the presence of missing data. Mol. Ecol. Resour. 21, 1359–1368 (2021).

24. Nei, M. & Li, W. H. Mathematical model for studying genetic variation in terms of restriction endonucleases. Proc. Natl. Acad. Sci. 76, 5269–5273 (1979).

25. Roux, C. et al. Recent and Ancient Signature of Balancing Selection around the S-Locus in Arabidopsis halleri and A. lyrata. Mol. Biol. Evol. 30, 435–447 (2013).

26. Yang, Q., Zhang, D., Li, Q., Cheng, Z. & Xue, Y. Heterochromatic and genetic features are consistent with recombination suppression of the self incompatibility locus in *Antirrhinum*. Plant J. 51, 140–151 (2007).

27. Jing, X. et al. Genetic diversity of the self-incompatibility locus in diploid potato. J. Integr. Agric. S2095311924004209 (2024) doi:10.1016/j.jia.2024.12.011.

28. Harbord, R. M., Napoli, C. A. & Robbins, T. P. Segregation Distortion of T-DNA Markers Linked to the Self-Incompatibility (S) Locus in Petunia hybrida. Genetics 154, 1323–1333 (2000).

29. Vázquez, N. et al. BDBM 1.0: A Desktop Application for Efficient Retrieval and Processing of High-Quality Sequence Data and Application to the Identification of the Putative Coffea S-Locus. Interdiscip. Sci. Comput. Life Sci. 11, 57–67 (2019).

30. Cao, Z.-H. et al. An *S* –locus F-box protein as pollen *S* determinant targets non-self S-RNase underlying self-incompatibility in *Citrus*. J. Exp. Bot. 75, 3891–3902 (2024).

31. Gu, C. et al. Long-read genome sequencing reveals the sequence characteristics of pear self-incompatibility locus. Mol. Hortic. 5, 13 (2025).

32. Chen, L.-Y. et al. The bracteatus pineapple genome and domestication of clonally propagated crops. Nat. Genet. 51, 1549–1558 (2019).

33. Wohlwend, J. et al. Boltz-1 Democratizing Biomolecular Interaction Modeling. Preprint at 10.1101/2024.11.19.624167 (2024).

34. Lombardi, G. & Carbone, A. MuLAN: Mutation-driven Light Attention Networks for investigating protein-protein interactions from sequences. Preprint at 10.1101/2024.08.24.609515 (2024).

35. Jurka, J. et al. Repbase Update, a database of eukaryotic repetitive elements. Cytogenet. Genome Res. 110, 462–467 (2005).

36. Ishimizu, T. et al. Identification of regions in which positive selection may operate in S RNase of Rosaceae: Implication for *S* allele specific recognition sites in S RNase. FEBS Lett. 440, 337–342 (1998).

37. Ushijima, K. et al. Cloning and characterization of cDNAs encoding S-RNases from almond (Prunus dulcis): primary structural features and sequence diversity of the S-RNases in Rosaceae. Mol. Gen. Genet. MGG 260, 261–268 (1998).

38. Branco, S. et al. Multiple convergent supergene evolution events in mating-type chromosomes. Nat. Commun. 9, 2000 (2018).

39. Kim, H.-T. et al. Identification and characterization of *S-RNase* genes in apple rootstock and the diversity of *S-RNases* in *Malus* species. J. Plant Biotechnol. 43, 49–57 (2016).

40. Daccord, N. et al. High-quality de novo assembly of the apple genome and methylome dynamics of early fruit development. Nat. Genet. 49, 1099–1106 (2017).

41. Qin, S. et al. A chromosome-scale genome assembly of Malus domestica, a multi-stress resistant apple variety. Genomics 115, 110627 (2023).

42. Aguiar, B. et al. Convergent Evolution at the Gametophytic Self-Incompatibility System in Malus and Prunus. PLOS ONE 10, e0126138 (2015).

43. Simão, F. A., Waterhouse, R. M., Ioannidis, P., Kriventseva, E. V. & Zdobnov, E. M. BUSCO: assessing genome assembly and annotation completeness with single-copy orthologs. Bioinformatics 31, 3210–3212 (2015).

44. Dreesen, R. S. G. et al. Analysis of Malus S-RNase gene diversity based on a comparative study of old and modern apple cultivars and European wild apple. Mol. Breed. 26, 693–709 (2010).

45. Larsen, B., Ørgaard, M., Toldam-Andersen, T. B. & Pedersen, C. A high-throughput method for genotyping S-RNase alleles in apple. Mol. Breed. 36, 24 (2016).

46. Ma, X., Cai, Z., Liu, W., Ge, S. & Tang, L. Identification, genealogical structure and population genetics of S-alleles in Malus sieversii, the wild ancestor of domesticated apple. Heredity 119, 185–196 (2017).

47. Vieira, J., Ferreira, P. G., Aguiar, B., Fonseca, N. A. & Vieira, C. P. Evolutionary patterns at the RNase based gametophytic self –incompatibility system in two divergent Rosaceae groups (Maloideae and Prunus). BMC Evol. Biol. 10, 200 (2010).

48. Okada, K., Moriya, S., Haji, T. & Abe, K. Isolation and characterization of multiple F-box genes linked to the S 9 – and S 10 –RNase in apple (Malus × domestica Borkh.). Plant Reprod. 26, 101– 111 (2013).

49. Sievers, F. & Higgins, D. G. Clustal Omega. Curr. Protoc. Bioinforma. 48, (2014).

50. Minh, B. Q., et al. Corrigendum to: IQ-TREE 2: New Models and Efficient Methods for Phylogenetic Inference in the Genomic Era. Mol. Biol. Evol. 37, 2461–2461 (2020).

51. Kalyaanamoorthy, S., Minh, B. Q., Wong, T. K. F., Von Haeseler, A. & Jermiin, L. S. ModelFinder: fast model selection for accurate phylogenetic estimates. Nat. Methods 14, 587–589 (2017).

52. Ward, J. H. Hierarchical Grouping to Optimize an Objective Function. J. Am. Stat. Assoc. 58, 236–244 (1963).

53. R Core Team. R: A language and environment for statistical computing. R Foundation for Statistical Computing, Vienna, Austria (2021).

54. Paradis, E. & Schliep, K. ape 5.0: an environment for modern phylogenetics and evolutionary analyses in R. Bioinformatics 35, 526–528 (2019).

55. Campo Elias Pardo et al. FactoClass: Combination of Factorial Methods and Cluster Analysis. The R Foundation 10.32614/cran.package.factoclass (2008).

56. Emms, D. M. & Kelly, S. OrthoFinder: phylogenetic orthology inference for comparative genomics. Genome Biol. 20, (2019).

57. H. Pagès, P. A. Biostrings. Bioconductor 10.18129/B9.BIOC.BIOSTRINGS (2017).

58. Marçais, G. et al. MUMmer4: A fast and versatile genome alignment system. PLOS Comput. Biol. 14, e1005944 (2018).

59. Goel, M., Sun, H., Jiao, W.-B. & Schneeberger, K. SyRI: finding genomic rearrangements and local sequence differences from whole-genome assemblies. Genome Biol. 20, 277 (2019).

60. Goel, M. & Schneeberger, K. plotsr: visualizing structural similarities and rearrangements between multiple genomes. Bioinformatics 38, 2922–2926 (2022).

61. Dunn, N. A. et al. Apollo: Democratizing genome annotation. PLOS Comput. Biol. 15, e1006790 (2019).

62. Jacques Dainat et al. NBISweden/AGAT: AGAT v1.5.0. Zenodo 10.5281/ZENODO.3552717 (2025).

63. Hackl, T., Ankenbrand, M., van Adrichem, B., Wilkins, D. & Haslinger, K. gggenomes: effective and versatile visualizations for comparative genomics. Preprint at 10.48550/ARXIV.2411.13556 (2024).

64. Buchfink, B., Xie, C. & Huson, D. H. Fast and sensitive protein alignment using DIAMOND. Nat. Methods 12, 59–60 (2015).

65. Zhao, T. & Schranz, M. E. Network approaches for plant phylogenomic synteny analysis. Curr. Opin. Plant Biol. 36, 129–134 (2017).

66. Chen, X. et al. Gene flow from the European wild apple and selection shaped the domesticated apple genome. Curr. Biol. 36, 2104–2118.e7 (2026).

67. Cingolani, P. et al. A program for annotating and predicting the effects of single nucleotide polymorphisms, SnpEff: SNPs in the genome of Drosophila melanogaster strain w1118; iso-2; iso-3. Fly (Austin) 6, 80–92 (2012).

68. Pedersen, B. S. & Quinlan, A. R. cyvcf2: fast, flexible variant analysis with Python. Bioinformatics 33, 1867–1869 (2017).

69. McKinney, W. Data Structures for Statistical Computing in Python. in 56–61 (Austin, Texas, 2010). doi:10.25080/Majora-92bf1922-00a.

70. Hunter, J. D. Matplotlib: A 2D Graphics Environment. Comput. Sci. Eng. 9, 90–95 (2007).

71. Yang, Z. PAML 4: Phylogenetic Analysis by Maximum Likelihood. Mol. Biol. Evol. 24, 1586– 1591 (2007).

72. Lê, S., Josse, J. & Husson, F. **FactoMineR**: An *R* Package for Multivariate Analysis. J. Stat. Softw. 25, (2008).

73. Weaver, S. et al. Datamonkey 2.0: A Modern Web Application for Characterizing Selective and Other Evolutionary Processes. Mol. Biol. Evol. 35, 773–777 (2018).

74. Almeida-Silva, F. & Van De Peer, Y. doubletrouble:an R/Bioconductor package for the identification, classification, and analysis of gene and genome duplications. Preprint at 10.1101/2024.02.27.582236 (2024).

75. Hahn, M. W. Distinguishing Among Evolutionary Models for the Maintenance of Gene Duplicates. J. Hered. 100, 605–617 (2009).

76. Wang, Y., Wang, X. & Paterson, A. H. Genome and gene duplications and gene expression divergence: a view from plants. Ann. N. Y. Acad. Sci. 1256, 1–14 (2012).

77. Freeling, M. Bias in Plant Gene Content Following Different Sorts of Duplication: Tandem, Whole-Genome, Segmental, or by Transposition. Annu. Rev. Plant Biol. 60, 433–453 (2009).

78. Wang, Y., Ficklin, S. P., Wang, X., Feltus, F. A. & Paterson, A. H. Large-Scale Gene Relocations following an Ancient Genome Triplication Associated with the Diversification of Core Eudicots. PLOS ONE 11, e0155637 (2016).

79. Sawyer, S. Statistical tests for detecting gene conversion. Mol. Biol. Evol. https://doi.org/10.1093/oxfordjournals.molbev.a040567 (1989) doi:10.1093/oxfordjournals.molbev.a040567.

80. Flutre, T., Duprat, E., Feuillet, C. & Quesneville, H. Considering Transposable Element Diversification in De Novo Annotation Approaches. PLoS ONE 6, e16526 (2011).

81. Quesneville, H. et al. Combined Evidence Annotation of Transposable Elements in Genome Sequences. PLoS Comput. Biol. 1, e22 (2005).

82. Sun, X. et al. Phased diploid genome assemblies and pan-genomes provide insights into the genetic history of apple domestication. Nat. Genet. 52, 1423–1432 (2020).

83. Hoede, C. et al. PASTEC: An Automatic Transposable Element Classification Tool. PLoS ONE 9, e91929 (2014).

84. Saidi, S., Blaison, M., Del Pilar Rodríguez-Ordóñez, M., Confais, J. & Quesneville, H. A reference-free pipeline for detecting shared transposable elements from pan-genomes to retrace their dynamics in a species. Genome Biol. 27, 117 (2026).

85. Wickham, H. ggplot2. *WIREs Comput*. Stat. 3, 180–185 (2011).

86. Schrödinger, L. The PyMOL Molecular Graphics System. (2026).

87. Chen, S. Ultrafast one pass FASTQ data preprocessing, quality control, and deduplication using fastp. iMeta 2, e107 (2023).

88. Dobin, A. et al. STAR: ultrafast universal RNA-seq aligner. Bioinformatics 29, 15–21 (2013).

89. Danecek, P. et al. Twelve years of SAMtools and BCFtools. GigaScience 10, giab008 (2021).

90. Pertea, M. et al. StringTie enables improved reconstruction of a transcriptome from RNA-seq reads. Nat. Biotechnol. 33, 290–295 (2015).

91. Pertea, G. & Pertea, M. GFF Utilities: GffRead and GffCompare. F1000Research 9, 304 (2020).

92. Kang, Y.-J. et al. CPC2: a fast and accurate coding potential calculator based on sequence intrinsic features. Nucleic Acids Res. 45, W12–W16 (2017).

93. Wang, L. et al. CPAT: Coding-Potential Assessment Tool using an alignment-free logistic regression model. Nucleic Acids Res. 41, e74–e74 (2013).

94. Li, A. et al. PLEKv2: predicting lncRNAs and mRNAs based on intrinsic sequence features and the coding-net model. BMC Genomics 25, 756 (2024).

95. Negri, T. D. C. et al. Pattern recognition analysis on long noncoding RNAs: a tool for prediction in plants. Brief. Bioinform. 20, 682–689 (2019).

96. Patro, R., Duggal, G., Love, M. I., Irizarry, R. A. & Kingsford, C. Salmon provides fast and bias-aware quantification of transcript expression. Nat. Methods 14, 417–419 (2017).

97. Fu, L., Niu, B., Zhu, Z., Wu, S. & Li, W. CD-HIT: accelerated for clustering the next-generation sequencing data. Bioinformatics 28, 3150–3152 (2012).

98. Love, M. I., Huber, W. & Anders, S. Moderated estimation of fold change and dispersion for RNA-seq data with DESeq2. Genome Biol. 15, 550 (2014).

