## Supplementary note 1 for "Innovation within stability: balancing selection preserves multigenic S-haplotype architecture in apple"

**Supplementary Note 1: SFBB families’ determination and validation**

To determine SFBB families of the 500 complete SFBB sequences, the amino acid sequences were aligned with sequences from the database (Supplementary Table 5), and phylogeny was constructed with 1000 bootstraps. *Eriobotya japonica* (Loquat) Fbox sequence (accession AGM21636.1) was used as an outgroup. Then, Ward hierarchical clustering (Ward, 1963) based on cophenetic distances between tree nodes (R package *stats,* R Core Team, 2021) as well as orthogroups determination using Orthofinder (Emms & Kelly, 2019) were applied. Ward clustering segregated SFBB sequences into 18 groups, while Orthofinder defined 16 orthogroups (Supplementary Fig. 3). The main difference is that a big Orthofinder orthogroup contains 3 Ward groups.

We took advantage of publicly available SFBB gene accessions, for which the family they belong to is known, to determine the family of SFBB genes identified from the assemblies we used. In most cases, public accessions tagged with identical SFBB families clustered together, allowing us to attribute the SFBB family to all the clusters. But in some cases, we found that public accessions tagged with identical SFBB families belong to several groups in our phylogeny. For instance, public accessions tagged with the SFBB3 family are found in two distant groups (Supplementary Fig. 3). Public accessions tagged with the SFBB1 family clustered together in our phylogenetic tree (Supplementary Fig. 3); however, hierarchical clustering and Orthofinder allowed us to split the group they form into two groups. The same conclusions can be made for public accessions tagged with the SFBB8 family (Supplementary Fig. 3).

To validate that these families should indeed be divided into two distinct groups, we compared the pairwise similarity distribution among sequences, considering both inter- and intra-group sequence similarity. We observed that sequences within each proposed subgroup displayed significantly higher similarity to one another compared to sequences from the other subgroup, supporting the hypothesis of functional divergence between these groups. Furthermore, when examining gene structure and conserved motifs, we identified differences that were consistent with the proposed division. This evidence collectively supports the reclassification of the SFBB1, SFBB3, and SFBB8 families into two distinct families each: SFBB1 and SFBB15, SFBB3 and SFBB4, and SFBB8 and SFBB13 (Supplementary Fig. 4, A, B and C).

On the contrary, families SFBB4 and SFBB12 can be brought together. Computing intra-family identity percentages considering that these two families form only one, showed that all intra-superfamily identity percentages are higher than 90% (Supplementary Fig. 4, D). For families SFBB9 and SFBB10, Ward hierarchical clustering and Orthofinder defined two groups (Supplementary Fig. 3), but their limits are not the same as the original SFBB families. The best thing to do is probably to redefine the SFBB9 and SFBB10 boundaries, as the Ward groups and orthogroups are supported by a branch bootstrap of 100%. Finally, some new families were created because no SFBBs with a family association were present in some groups. Eighteen updated SFBB families were finally defined (Supplementary Table 7, Supplementary Fig. 3). SFBB genes annotated on *Malus* haplotypes, coordinates, and families are summarized in Supplementary Table 8.

Considering these updated SFBB families, their presence was checked in each haplotype (figure 1, B). In each haplotype, 0 to 2 genes from each SFBB family were identified. Since some S alleles are represented in this dataset by multiple haplotypes, it was possible to compare them, and observe that some SFBB families are absent from all haplotypes with the same S allele. For instance, SFBB11 is absent from all MdS4 (h20 and h22) and MdS5 (h2 and h26) haplotypes. SFBB2 family is absent from all MdS2 haplotypes except h23 (h3, h10, h13 and h14). Absent SFBB families’products could target these S-RNases. Concerning haplotypes from S allele MdSf, all identified SFBB families are present, suggesting that the SFBB family whose products target this S-RNase was not identified from our dataset, or that one of the SFBB families identified contain unfunctional products (although the protein sequences are complete).

References:

Emms, D. M., & Kelly, S. (2019). OrthoFinder: Phylogenetic orthology inference for comparative genomics. *Genome Biology*, *20*(1). https://doi.org/10.1186/s13059-019-1832-y

R Core Team. (2021). *R: A language and environment for statistical computing.* [Computer software]. R Foundation for Statistical Computing, Vienna, Austria. https://www.R-project.org/

Ward, J. H. (1963). Hierarchical Grouping to Optimize an Objective Function. *Journal of the American Statistical Association*, *58*(301), 236–244. https://doi.org/10.1080/01621459.1963.10500845
