## Supplementary note 2 for "Innovation within stability: balancing selection preserves multigenic S-haplotype architecture in apple"

Supplementary Note 2: lncRNA annotation and tissue-specific expression in the *Malus* S-locus

### RNA-seq alignment to reference genomes

Paired-end RNA-seq reads from three biological replicates of pollen, pistil, and leaf tissues from two apple genotypes (Gala and Honeycrisp) were aligned independently to each of the three reference assemblies (h2: Gala haplotype 1, h3: Gala haplotype 2 (Sun et al. 2020), and h10: Honeycrisp (Khan et al. 2022) using STAR v2.7.11b (Dobin et al. 2013) with default parameters and paired-end mode (--outSAMtype BAM Unsorted). Alignments to chromosome 17 were then extracted from each full-genome BAM file using samtools view (samtools v1.21; Danecek et al. 2021).

### lncRNA discovery and annotation

Transcripts were assembled independently for each sample using StringTie v3.0.0 (Pertea et al. 2015) in reference-guided mode. Per-sample assemblies were merged using stringtie --merge with the reference gene annotation provided as a guide. The merged assembly was compared to the reference annotation using Gffcompare v0.12.10 (Pertea and Pertea 2020). Transcripts assigned class codes "u" (intergenic), "x" (antisense to an annotated gene), or "i" (intronic) were retained as lncRNA candidates, thereby excluding transcripts overlapping known annotated genes on the same strand.

Candidate transcripts were required to have a spliced sequence length ≥ 200 bp. Transcript sequences were extracted from the reference genome using gffread v0.12.7 (Pertea and Pertea 2020).

Coding potential was assessed using four independent tools: CPC2 (web server; https://cpc2.gao-lab.org/; Kang et al. 2017), CPAT v3.0.5 (coding probability cutoff = 0.44 ; Wang et al. 2013), PLEK2 v2.1 (Li et al. 2024), and RNAplonc v1.0 (Negri et al. 2019). A transcript was classified as non-coding if at least three out of four tools predicted it to be non-coding (majority-vote ensemble).

### Transcript quantification in the S-locus region

Transcript-level quantification within the S-locus was performed using Salmon v1.10.3 (Patro et al. 2017). A dedicated reference transcriptome was constructed for each haplotype by combining the sequences of all annotated S-locus lncRNAs with those of known S-locus protein-coding genes (SFBB family members and S-RNase). Salmon indices were built with a k-mer size of 31 (-k 31). Quantification was run in quasi-mapping mode with --validateMappings, and the library type was auto-detected (-l A). Transcript-per-million (TPM) values were aggregated into per-reference expression matrices for downstream analyses.

### Identification of tissue-specific lncRNAs

Pollen-specific and pistil-specific lncRNAs were identified from the S-locus TPM matrices using a stringent expression-based criterion. A lncRNA was classified as tissue-specific if it satisfied all of the following conditions in the target tissue: (i) mean TPM across replicates ≥ 1.0; (ii) at least 2 out of 3 replicates had TPM ≥ 1.0 in pollen ; and (iii) mean TPM ≤ 1.0 in each of the two other tissues. Analyses were performed independently for each of the three reference haplotypes, and results were combined into a single table.

The genomic coordinates of tissue-specific lncRNAs were extracted from the lncRNA annotation GTF files and expressed relative to the boundaries of the S-locus region defined for each haplotype. Positions of S-RNase and SFBB family genes within the S-locus were retrieved from the reference gene annotation and displayed alongside the lncRNA positions.

### Sequence similarity analysis

Transcript sequences of pollen-specific and pistil-specific lncRNAs were pooled across the three reference haplotypes. Pairwise sequence similarity was evaluated using three complementary approaches: (1) k-mer Pearson correlation. For each transcript, 4-mer frequencies were computed, and z-score normalized across all sequences (mean-centered and scaled by standard deviation per k-mer), following the SEEKR framework (Li et al. 2024). Pairwise Pearson correlation coefficients between normalized k-mer profiles were computed and reported as a pairwise similarity matrix. (2) BLAST all-vs-all. Pairwise nucleotide alignments were performed using BLASTN v2.16.0 (Ye et al. 2006) (-dust no, e-value threshold 1×10⁻²). Self-hits were removed, and results were annotated with query and subject coverage. (3) minimap2 all-vs-all. Pairwise alignments were additionally computed using minimap2 v2.28 (Li 2018) with the asm20 preset, which tolerates up to ~20% sequence divergence, allowing detection of more divergent homologs across haplotypes.

### Structural prediction of S-RNase–SFBB–lncRNA complexes

To investigate whether pollen-specific S-locus lncRNAs modulate the interaction between S-RNase proteins and SFBB (S-locus F-box brother) proteins in Malus domestica, co-folding predictions were performed using Boltz v2.2.1 (Passaro et al. 2025). Two biologically motivated S-allele pairs were tested following the non-self recognition model: MdS5 S-RNase sequences against SFBB11 copies, and MdS2 S-RNase sequences against SFBB2 copies. For each S-RNase × SFBB pair, two types of complexes were modeled: binary complexes comprising the S-RNase (chain A) and the SFBB protein (chain B), and ternary complexes additionally including one pollen-specific lncRNA identified at the S-locus (chain C). To ensure biological relevance, lncRNA sequences were matched to the haplotype of the SFBB being tested, reflecting co-occurrence in the same pollen grain, using the pollen-specific S-locus lncRNAs identified in our three reference haplotypes: MSTRG.2343.1 for Gala H1 (h2), MSTRG.2154.1 for Gala H2 (h3), and MSTRG.2350.1, MSTRG.2329.1, MSTRG.2384.1 and MSTRG.2363.1 for Honeycrisp (h10). The goal was to compare binary and ternary predictions within matched haplotype pairs. In total, 24 predictions were run (Supplementary Table 20). Predictions were run in single-sequence mode without multiple sequence alignment (MSA), which reduces computation time while preserving the ability to compare binary and ternary complexes under identical conditions. Interface predicted TM-score (ipTM) was extracted from the JSON confidence output and used to compare binary and ternary predictions for each matched pair.

**Results**

### RNA-seq alignment quality

RNA-seq data from nine samples per reference genome (three biological replicates each for leaf, pollen and pistil) were aligned to three *Malus* reference assemblies. Across all samples, the unique mapping rate was consistently high (Supplementary Fig. 26, Supplementary Table 21). For Gala haplotype 1, unique mapping rates ranged from 86.2% to 88.9% (mean: 87.6%; total input reads per sample: 41.9–55.8 M). For Gala haplotype 2, rates were slightly lower, ranging from 78.3% to 80.1% (mean: 79.5%), consistent with the expected incomplete coverage of a haplotype-resolved assembly when using Gala reads. For Honeycrisp, unique mapping rates were the highest, ranging from 90.9% to 95.0% (mean: 93.6%; 40.2–63.7 M reads), reflecting the use of its own reference assembly. After extraction of chromosome 17 reads and chromosome identifier harmonization, BAM files were used for all downstream lncRNA analyses.

### lncRNA discovery and annotation on S-loci

The annotation pipeline was applied independently to each of the three reference haplotypes. After transcript assembly with StringTie and comparison to the reference annotation with Gffcompare, transcripts assigned class codes "u" (intergenic), "x" (antisense) or "i" (intronic) were retained. Following the length filter (≥ 200 bp), 31, 42 and 32 candidate transcripts were obtained for Gala haplotype 1, Gala haplotype 2 and Honeycrisp, respectively.

Coding potential was assessed with four tools (CPC2, CPAT, PLEK2, RNAplonc, Supplementary Table 22). The fraction of transcripts predicted as non-coding varied across tools but was consistently high (Supplementary Table 22). After applying the majority-vote ensemble filter (≥ 3 out of 4 tools predicting non-coding), 25, 34 and 24 lncRNA transcripts were retained for Gala haplotype 1, Gala haplotype 2 and Honeycrisp, respectively.

### lncRNA quantification in the S-locus region

Salmon quantification was restricted to the S-locus region, using a dedicated transcriptome combining S-locus lncRNAs with known genes. The S-locus transcriptomes comprised 74, 86 and 64 transcripts for Gala haplotype 1, Gala haplotype 2 and Honeycrisp, respectively, of which 25, 34 and 24 were lncRNAs (83 lncRNA transcripts in total across the three haplotypes).

### Tissue-specific lncRNAs in the S-locus

Applying the tissue-specificity criteria (mean TPM ≥ 1.0 in pollen, mean TPM ≤ 1.0 in each of leaf and pistil, ≥ 2/3 replicates above threshold), 6 pollen-specific lncRNAs were identified across the three haplotypes: 1 in Gala haplotype 1 (MSTRG.2343.1), 1 in Gala haplotype 2 (MSTRG.2154.1), and 4 in Honeycrisp (MSTRG.2329.1, MSTRG.2350.1, MSTRG.2363.1, MSTRG.2384.1), representing 7.2% of all quantified S-locus lncRNAs (Supplementary Fig. 27).

Applying the same criteria with pistil as the target tissue, no pistil-specific lncRNAs were identified in any of the three haplotypes under the current TPM thresholds, suggesting that S-locus lncRNA expression is not enriched in pistil relative to the other tissues assayed, or that pistil-enriched lncRNAs do not pass the stringent specificity threshold applied here.

The six pollen-specific lncRNAs were mapped to their positions within the S-locus boundaries (Supplementary Table 23; Supplementary Fig. 28). In Honeycrisp, the four positions spanned the full length of the S-locus, from 2.8% to 90.4% of the total S-locus length (Supplementary Fig. 28). No clustering of pollen-specific lncRNAs around S-RNase or any particular SFBB family was apparent.

The six pollen-specific lncRNA sequences were compared pairwise using three complementary approaches. K-mer Pearson correlation (k = 4) computed across all 15 pairwise combinations yielded low similarity values (maximum r = 0.105, between Honeycrisp MSTRG.2329.1 and MSTRG.2350.1; range: −0.47 to 0.11), indicating divergent sequence composition across pollen-specific lncRNAs. To assess whether pollen-specific lncRNAs share partial sequence similarity, pairwise local alignments were performed using BLASTN with progressively sensitive settings (word_size = 11 and 7, e-value ≤ 0.01, dust masking disabled). Both searches identified a single cross-haplotype alignment: a 15-bp perfect match between Gala haplotype 2 MSTRG.2154.1 and Honeycrisp MSTRG.2350.1 (e-value = 0.004, bitscore = 28.8), which is below the length threshold generally considered meaningful for lncRNA homology inference (Li et al. 2023). Together, these results indicate that pollen-specific lncRNAs identified across haplotypes do not share detectable sequence homology, suggesting independent transcriptional origins at each locus rather than conservation of a common pollen-expressed lncRNA.

### Structural modeling of S-RNase–SFBB interactions with pollen-specific lncRNAs

To assess whether pollen-specific S-locus lncRNAs could modulate the interaction between S-RNase and SFBB proteins, we performed Boltz-2 co-folding predictions for binary (S-RNase + SFBB) and ternary (S-RNase + SFBB + lncRNA) complexes across two S-allele pairs (MdS2/SFBB2 and MdS5/SFBB11). Predicted interface quality was evaluated using the interface predicted TM-score (ipTM). For the MdS2 S-RNase × SFBB2 pair, addition of the pollen-specific lncRNA MSTRG.2343.1 (Gala Hap1) consistently increased ipTM across all five MdS2 S-RNase variants tested (mean binary ipTM = 0.115 ± 0.008 vs mean ternary ipTM = 0.181 ± 0.002; mean Δ = +0.067). This effect was remarkably uniform regardless of the S-RNase haplotype origin (Gala Hap2 (h3), Honeycrisp (h10), h13, h14, h23), suggesting that MSTRG.2343.1 consistently promotes a higher-confidence predicted arrangement of the MdS2 S-RNase–SFBB2 complex (Supplementary Fig. 29). For the MdS5 S-RNase × SFBB11 pair, the effect of lncRNAs was more variable. Binary complexes showed a mean ipTM of 0.143 ± 0.021. Among the five lncRNAs tested, only MSTRG.2350.1 (Honeycrisp) showed a consistent positive effect (mean Δ = +0.026), while MSTRG.2329.1 had a slightly negative effect (mean Δ = −0.013). The remaining lncRNAs showed effects close to zero (Δ ranging from −0.009 to +0.010), with no consistent directional trend (Supplementary Table 20).

Overall, all ipTM values remained below 0.20, well below the threshold generally associated with confident structural predictions in AlphaFold-like models (ipTM > 0.5). These results therefore do not provide evidence of stable direct interactions between S-RNase and SFBB proteins in the configurations tested. Nevertheless, the systematic increase in ipTM observed for MdS2 × SFBB2 in the presence of MSTRG.2343.1 suggests that this lncRNA may influence the structural compatibility of this complex, warranting further investigation.

**References**

Danecek P, Bonfield JK, Liddle J, Marshall J, Ohan V, Pollard MO, Whitwham A, Keane T, McCarthy SA, Davies RM, et al. 2021. Twelve years of SAMtools and BCFtools. *GigaScience* 10:giab008.

Dobin A, Davis CA, Schlesinger F, Drenkow J, Zaleski C, Jha S, Batut P, Chaisson M, Gingeras TR. 2013. STAR: ultrafast universal RNA-seq aligner. *Bioinformatics* 29:15–21.

Kang Y-J, Yang D-C, Kong L, Hou M, Meng Y-Q, Wei L, Gao G. 2017. CPC2: a fast and accurate coding potential calculator based on sequence intrinsic features. *Nucleic Acids Res.* 45:W12–W16.

Khan A, Carey SB, Serrano A, Zhang H, Hargarten H, Hale H, Harkess A, Honaas L. 2022. A phased, chromosome-scale genome of ‘Honeycrisp’ apple (Malus domestica). *Gigabyte* 2022:1–15.

Li A, Zhou Haotian, Xiong S, Li J, Mallik S, Fei R, Liu Y, Zhou Hongfang, Wang X, Hei X, et al. 2024. PLEKv2: predicting lncRNAs and mRNAs based on intrinsic sequence features and the coding-net model. *BMC Genomics* 25:756.

Li H. 2018. Minimap2: pairwise alignment for nucleotide sequences.Birol I, editor. *Bioinformatics* 34:3094–3100.

Li S, Eberhard QE, Ni L, Calabrese JM. 2024. Improved functions for nonlinear sequence comparison using SEEKR. *RNA* 30:1408–1421.

Li Z, Liu L, Feng C, Qin Y, Xiao J, Zhang Z, Ma L. 2023. LncBook 2.0: integrating human long non-coding RNAs with multi-omics annotations. *Nucleic Acids Res.* 51:D186–D191.

Negri TDC, Alves WAL, Bugatti PH, Saito PTM, Domingues DS, Paschoal AR. 2019. Pattern recognition analysis on long noncoding RNAs: a tool for prediction in plants. *Brief. Bioinform.* 20:682–689.

Passaro S, Corso G, Wohlwend J, Reveiz M, Thaler S, Somnath VR, Getz N, Portnoi T, Roy J, Stark H, et al. 2025. Boltz-2: Towards Accurate and Efficient Binding Affinity Prediction. Available from: http://biorxiv.org/lookup/doi/10.1101/2025.06.14.659707

Patro R, Duggal G, Love MI, Irizarry RA, Kingsford C. 2017. Salmon provides fast and bias-aware quantification of transcript expression. *Nat. Methods* 14:417–419.

Pertea G, Pertea M. 2020. GFF Utilities: GffRead and GffCompare. *F1000Research* 9:304.

Pertea M, Pertea GM, Antonescu CM, Chang T-C, Mendell JT, Salzberg SL. 2015. StringTie enables improved reconstruction of a transcriptome from RNA-seq reads. *Nat. Biotechnol.* 33:290–295.

Sun X, Jiao C, Schwaninger H, Chao CT, Ma Y, Duan N, Khan A, Ban S, Xu K, Cheng L, et al. 2020. Phased diploid genome assemblies and pan-genomes provide insights into the genetic history of apple domestication. *Nat. Genet.* 52:1423–1432.

Wang L, Park HJ, Dasari S, Wang S, Kocher J-P, Li W. 2013. CPAT: Coding-Potential Assessment Tool using an alignment-free logistic regression model. *Nucleic Acids Res.* 41:e74–e74.

Ye J, McGinnis S, Madden TL. 2006. BLAST: improvements for better sequence analysis. *Nucleic Acids Res.* 34:W6–W9.
