## Supplementary note 3 for "Innovation within stability: balancing selection preserves multigenic S-haplotype architecture in apple"

### Supplementary Note 3: Details on dataset used, DNA extraction, PacBio sequencing, assembly of the new genomes, TEs and gene annotation

Our dataset included 15 already available haplotype assemblies and 12 newly sequenced haplotypes, for this study, only chromosome 17 was kept, as this chromosome hosts the SI loci in *Malus* (Aguiar et al., 2015). All high-quality assembled apple genomes available in 2022 were retrieved from public repositories. One haplotype from the *M. domestica* double-haploid GDDH13 reference genome (Daccord et al., 2017), two phased haplotypes of a Gala cultivar (Sun et al., 2020), and two available phased haplotypes of a Fuji cultivar (Li et al., 2024) were used. For cultivars Red Delicious (Han et al., 2017), Honeycrisp (Khan et al., 2022), Hanfu (Qin et al., 2023), Brown Snout, Costard, Bardsey Island, and Newtons (Könyves et al., 2022), only one phased haplotype was available and was used in this study. Similarly, for individuals of *M. sieversii* native to Central Asia (Sun et al., 2020), *M. sylvestris* native to Germany (Sun et al., 2020), and native to England (Ruhsam et al., 2022), only one phased haplotype was available and collected for this study. In total, 12 *M. domestica,* two *M. sylvestris* and one *M. sieversii* haplotypes were therefore retrieved from public repositories (Supplementary Table 1).

To complete this dataset, *M. sylvestris* individuals from Austria, Denmark, France and Romania, one *M. orientalis* individual native to Armenia, and one Spanish Cider *M. domestica* cultivar were grown in IDEEV facilities in order to sequence their genomes using PacBio Sequel II. Assembly pipeline used allowed us to have the two parental haplotypes for these genotypes, so twelve supplementary haplotypes. Ultimately, 16 *M. domestica*, 11 *M. sylvestris*, one *M. sieversii*, and two *M. orientalis* haplotypes, in total, 27 haplotypes, were available for this study.

Most individuals used to make this dataset are diploids and, in most cases, two assembled haplotypes were obtained per individual (double-haploid Golden Delicious, tri-haploid Hanfu, Gala and Fuji cultivars, the cider individual, *the M. orientalis* individual, the *M. sieversii* individual, and the four *M. sylvestris* individuals grown at IDEEV). For eight diploid genotypes (double-haploid Golden Delicious, Hanfu, Red Delicious, Honeycrisp, Brown Snout, Costard, Bardsey Island, and Newtons cultivars, *M. sieversii* and *M. sylvestris* from previous studies), we only have one assembled haplotype. In total, 14 haplotypes of *M. domestica*, 10 haplotypes of *M. sylvestris*, one haplotype of *M. sieversii* and two haplotypes of *M. orientalis* were used for further analyses (Supplementary Table 1). For convenience, all haplotypes were identified with id from h1 to h32. Finally, two haplotypes were collected from haploid individuals: the double-haploid Golden Delicious individual used to construct the GDDH13 *M. domestica* reference genome (Daccord et al., 2017), and the *M. domestica* cultivar Hanfu individual (Qin et al., 2023), which is a tri-haploid.

#### DNA extraction, library construction and sequencing

For h21 to h32, Genomic DNA was extracted from whole plant material using the QIAGEN Genomic-tips 100/G kit (Cat No./ID: 10243) following the tissue protocol. Briefly, ~1 g of plant tissue was frozen and ground in liquid nitrogen with a mortar and pestle. After 2 h of lysis at 50°C and a single centrifugation step, DNA was bound to the column, washed, and subsequently eluted. The eluate was desalted and concentrated by alcohol precipitation, and DNA was finally resuspended in EB buffer.

DNA concentration and purity were assessed using a NanoDrop One spectrophotometer (Thermo Scientific) and a Qubit 3 Fluorometer using the Qubit dsDNA BR assay (Invitrogen), respectively. DNA integrity and fragment size distribution were evaluated using a Femto Pulse system (Agilent, Santa Clara, CA, USA). Then, DNA was submitted to Leiden Genome Technology Center (Netherlands) for PacBio® HiFi library construction and sequencing.

#### Genome assemblies and scaffold anchoring

Raw PacBio reads were assembled with hifiasm (Cheng et al., 2021), with default parameters, which allowed obtaining two phased haplotypes. TIDK (Brown et al., 2025) was used to detect telomere repeats on resulting contigs. Contigs were anchored to chromosomes with ALLMAPS JCVI (Tang et al., 2015), which allows combining several genetic maps to order and orient contigs, and using RagTag scaffold (Alonge et al., 2022), which allows using whole-genome alignments to a reference assembly to scaffold query sequences. The comparison of AGP files resulting from ALLMAPS and RagTag was performed using homemade Perl scripts available on <https://github.com/CornilleEclecticLab/ART_pipeline/tree/main>, and AGPTOOLS (Ricemeyer et al., 2025). Briefly, Perl scripts allow selecting contigs associated with a specific chromosome in several AGP files and making a final AGP file synthesizing all results. Then, this final agp file was used as input with AGPTOOLS.

Concerning ALLMAPS anchoring, integrated genetic maps from Di Pierro et al., (2016) and from Howard et al., (2021) with SNP markers from the IRSC pear Infinium® II 1 K array (Montanari et al., 2019) and the apple 20K SNP array (Bianco et al., 2014) were used. Concerning RagTag scaffold, GDDH13 genome (Daccord et al., 2017) was used as reference to scaffold assembly contigs. Only chromosome 17 was then kept for further analyses.

#### Gene and transposable element annotation

To eliminate bias in genome annotation caused by different pipelines, gene annotation was performed using a single pipeline. First, TE libraries were constructed using REPET TEdenovo (Flutre et al., 2011) using 8 assemblies from several *Malus* species (GDDH13, cider, MSYL_AUT, MSYL_DNK, MSYL_FRA, MSYL_ROU, Msiev, MORI_ARM), in order to capture maximal diversity of TE content. Then, these TE libraries were merged, and redundancy was removed using PASTEC (Hoede et al., 2014). Finally, TEs were annotated in all assemblies using the constructed TEs panlibrary, and REPET TEannot (Quesneville et al., 2005). TE sequences were softmasked from assemblies using BEDOPS (Neph et al., 2012), to convert gff files to bed files and BEDTOOLS maskfasta (Quinlan & Hall, 2010) to softmask TE sequences.

The quality of repetitive genomic regions was assessed using the LTR Assembly Index (LAI) (Quesneville et al., 2005): (1) LTRharvest in GenomeTools v1.6.2 (Ellinghaus et al., 2008) was used to de novo predict the candidate LTR-RTs (full-length LTRs retrotransposon) in the haplotypes assembly sequences for chromosome 17, and (2) LTR_retriever v2.9.5 (Ou & Jiang, 2018) was then used to combine and refactor all the candidates to get the final full-length LTR-RTs. LAI was calculated based on the formula: LAI = (intact LTR-RT length/total LTR-RT length) × 100.

LAI values in the S-loci vary among haplotypes but are mostly above 10 (Supplementary Fig. 5), except in Gala haplotypes (h2 and h3), where assembly quality may not be sufficient and care must be taken with conclusions made on these haplotypes.

To predict genes, Helixer (Stiehler et al., 2021) was run on softmasked haplotype fasta files using land_plant as the –lineage model. Gffread (Pertea & Pertea, 2020) was used to get CDS and protein sequences for predicted genes. Finally, gene functional annotation was performed from protein sequences, using eggnog-mapper (Cantalapiedra et al., 2021; Huerta-Cepas et al., 2016, 2017).

To complete SFBB and S-RNase gene annotation, manual annotation was performed using Apollo (Dunn et al., 2019) to identify supplementary SFBB and S-RNase genes and to check SFBB and S-RNase gene structures.

#### References

Aguiar, B., Vieira, J., Cunha, A. E., Fonseca, N. A., Iezzoni, A., Van Nocker, S., & Vieira, C. P. (2015). Convergent Evolution at the Gametophytic Self-Incompatibility System in Malus and Prunus. *PLOS ONE*, *10*(5), e0126138. https://doi.org/10.1371/journal.pone.0126138

Alonge, M., Lebeigle, L., Kirsche, M., Jenike, K., Ou, S., Aganezov, S., Wang, X., Lippman, Z. B., Schatz, M. C., & Soyk, S. (2022). Automated assembly scaffolding using RagTag elevates a new tomato system for high-throughput genome editing. *Genome Biology*, *23*(1), 258. https://doi.org/10.1186/s13059-022-02823-7

Bianco, L., Cestaro, A., Sargent, D. J., Banchi, E., Derdak, S., Di Guardo, M., Salvi, S., Jansen, J., Viola, R., Gut, I., Laurens, F., Chagné, D., Velasco, R., Van De Weg, E., & Troggio, M. (2014). Development and Validation of a 20K Single Nucleotide Polymorphism (SNP) Whole Genome Genotyping Array for Apple (Malus × domestica Borkh). *PLoS ONE*, *9*(10), e110377. https://doi.org/10.1371/journal.pone.0110377

Brown, M. R., Manuel Gonzalez De La Rosa, P., & Blaxter, M. (2025). tidk: A toolkit to rapidly identify telomeric repeats from genomic datasets. *Bioinformatics*, *41*(2), btaf049. https://doi.org/10.1093/bioinformatics/btaf049

Cantalapiedra, C. P., Hernández-Plaza, A., Letunic, I., Bork, P., & Huerta-Cepas, J. (2021). eggNOG-mapper v2: Functional Annotation, Orthology Assignments, and Domain Prediction at the Metagenomic Scale. *Molecular Biology and Evolution*, *38*(12), 5825–5829. https://doi.org/10.1093/molbev/msab293

Cheng, H., Concepcion, G. T., Feng, X., Zhang, H., & Li, H. (2021). Haplotype-resolved de novo assembly using phased assembly graphs with hifiasm. *Nature Methods*, *18*(2), 170–175. https://doi.org/10.1038/s41592-020-01056-5

Daccord, N., Celton, J.-M., Linsmith, G., Becker, C., Choisne, N., Schijlen, E., van de Geest, H., Bianco, L., Micheletti, D., Velasco, R., Di Pierro, E. A., Gouzy, J., Rees, D. J. G., Guérif, P., Muranty, H., Durel, C.-E., Laurens, F., Lespinasse, Y., Gaillard, S., … Bucher, E. (2017). High-quality de novo assembly of the apple genome and methylome dynamics of early fruit development. *Nature Genetics*, *49*(7), 1099–1106. https://doi.org/10.1038/ng.3886

Di Pierro, E. A., Gianfranceschi, L., Di Guardo, M., Koehorst-van Putten, H. J., Kruisselbrink, J. W., Longhi, S., Troggio, M., Bianco, L., Muranty, H., Pagliarani, G., Tartarini, S., Letschka, T., Lozano Luis, L., Garkava-Gustavsson, L., Micheletti, D., Bink, M. C., Voorrips, R. E., Aziz, E., Velasco, R., … Van De Weg, W. E. (2016). A high-density, multi-parental SNP genetic map on apple validates a new mapping approach for outcrossing species. *Horticulture Research*, *3*(1), 16057. https://doi.org/10.1038/hortres.2016.57

Dunn, N. A., Unni, D. R., Diesh, C., Munoz-Torres, M., Harris, N. L., Yao, E., Rasche, H., Holmes, I. H., Elsik, C. G., & Lewis, S. E. (2019). Apollo: Democratizing genome annotation. *PLOS Computational Biology*, *15*(2), e1006790. https://doi.org/10.1371/journal.pcbi.1006790

Ellinghaus, D., Kurtz, S., & Willhoeft, U. (2008). LTRharvest, an efficient and flexible software for de novo detection of LTR retrotransposons. *BMC Bioinformatics*, *9*(1), 18. https://doi.org/10.1186/1471-2105-9-18

Flutre, T., Duprat, E., Feuillet, C., & Quesneville, H. (2011). Considering Transposable Element Diversification in De Novo Annotation Approaches. *PLoS ONE*, *6*(1), e16526. https://doi.org/10.1371/journal.pone.0016526

Han, M., Sun, Q., Zhou, J., Qiu, H., Guo, J., Lu, L., Mu, W., & Sun, J. (2017). Insertion of a solo LTR retrotransposon associates with spur mutations in ‘Red Delicious’ apple (Malus × domestica). *Plant Cell Reports*, *36*(9), 1375–1385. https://doi.org/10.1007/s00299-017-2160-x

Hoede, C., Arnoux, S., Moisset, M., Chaumier, T., Inizan, O., Jamilloux, V., & Quesneville, H. (2014). PASTEC: An Automatic Transposable Element Classification Tool. *PLoS ONE*, *9*(5), e91929. https://doi.org/10.1371/journal.pone.0091929

Howard, N. P., Troggio, M., Durel, C.-E., Muranty, H., Denancé, C., Bianco, L., Tillman, J., & Van De Weg, E. (2021). Integration of Infinium and Axiom SNP array data in the outcrossing species Malus × domestica and causes for seemingly incompatible calls. *BMC Genomics*, *22*(1), 246. https://doi.org/10.1186/s12864-021-07565-7

Huerta-Cepas, J., Forslund, K., Coelho, L. P., Szklarczyk, D., Jensen, L. J., von Mering, C., & Bork, P. (2017). Fast Genome-Wide Functional Annotation through Orthology Assignment by eggNOG-Mapper. *Molecular Biology and Evolution*, *34*(8), 2115–2122. https://doi.org/10.1093/molbev/msx148

Huerta-Cepas, J., Szklarczyk, D., Forslund, K., Cook, H., Heller, D., Walter, M. C., Rattei, T., Mende, D. R., Sunagawa, S., Kuhn, M., Jensen, L. J., von Mering, C., & Bork, P. (2016). eggNOG 4.5: A hierarchical orthology framework with improved functional annotations for eukaryotic, prokaryotic and viral sequences. *Nucleic Acids Research*, *44*(D1), D286–D293. https://doi.org/10.1093/nar/gkv1248

Khan, A., Carey, S. B., Serrano, A., Zhang, H., Hargarten, H., Hale, H., Harkess, A., & Honaas, L. (2022). A phased, chromosome-scale genome of ‘Honeycrisp’ apple (Malus domestica). *Gigabyte*, *2022*, 1–15. https://doi.org/10.46471/gigabyte.69

Könyves, K., Mian, S., Johns, J., Royal Botanic Garden Edinburgh Genome Acquisition Lab, Royal Botanic Gardens Kew Genome Acquisition Lab, Darwin Tree of Life Barcoding collective, Wellcome Sanger Institute Tree of Life programme, Wellcome Sanger Institute Scientific Operations: DNA Pipelines collective, Tree of Life Core Informatics collective, Ruhsam, M., Leitch, I. J., & Darwin Tree of Life Consortium. (2022). The genome sequence of the apple, Malus domestica (Suckow) Borkh., 1803. *Wellcome Open Research*, *7*, 297. https://doi.org/10.12688/wellcomeopenres.18646.1

Li, W., Chu, C., Li, H., Zhang, H., Sun, H., Wang, S., Wang, Z., Li, Y., Foster, T. M., López-Girona, E., Yu, J., Li, Y., Ma, Y., Zhang, K., Han, Y., Zhou, B., Fan, X., Xiong, Y., Deng, C. H., … Han, Z. (2024). Near-gapless and haplotype-resolved apple genomes provide insights into the genetic basis of rootstock-induced dwarfing. *Nature Genetics*, *56*(3), 505–516. https://doi.org/10.1038/s41588-024-01657-2

Montanari, S., Bianco, L., Allen, B. J., Martínez-García, P. J., Bassil, N. V., Postman, J., Knäbel, M., Kitson, B., Deng, C. H., Chagné, D., Crepeau, M. W., Langley, C. H., Evans, K., Dhingra, A., Troggio, M., & Neale, D. B. (2019). Development of a highly efficient Axiom^TM^ 70 K SNP array for Pyrus and evaluation for high-density mapping and germplasm characterization. *BMC Genomics*, *20*(1), 331. https://doi.org/10.1186/s12864-019-5712-3

Neph, S., Kuehn, M. S., Reynolds, A. P., Haugen, E., Thurman, R. E., Johnson, A. K., Rynes, E., Maurano, M. T., Vierstra, J., Thomas, S., Sandstrom, R., Humbert, R., & Stamatoyannopoulos, J. A. (2012). BEDOPS: High-performance genomic feature operations. *Bioinformatics*, *28*(14), 1919–1920. https://doi.org/10.1093/bioinformatics/bts277

Ou, S., Chen, J., & Jiang, N. (2018). Assessing genome assembly quality using the LTR Assembly Index (LAI). *Nucleic Acids Research*. https://doi.org/10.1093/nar/gky730

Ou, S., & Jiang, N. (2018). LTR_retriever: A Highly Accurate and Sensitive Program for Identification of Long Terminal Repeat Retrotransposons. *Plant Physiology*, *176*(2), 1410–1422. https://doi.org/10.1104/pp.17.01310

Pertea, G., & Pertea, M. (2020). GFF Utilities: GffRead and GffCompare. *F1000Research*, *9*, 304. https://doi.org/10.12688/f1000research.23297.2

Qin, S., Xu, G., He, J., Li, L., Ma, H., & Lyu, D. (2023). A chromosome-scale genome assembly of Malus domestica, a multi-stress resistant apple variety. *Genomics*, *115*(3), 110627. https://doi.org/10.1016/j.ygeno.2023.110627

Quesneville, H., Bergman, C. M., Andrieu, O., Autard, D., Nouaud, D., Ashburner, M., & Anxolabehere, D. (2005). Combined Evidence Annotation of Transposable Elements in Genome Sequences. *PLoS Computational Biology*, *1*(2), e22. https://doi.org/10.1371/journal.pcbi.0010022

Quinlan, A. R., & Hall, I. M. (2010). BEDTools: A flexible suite of utilities for comparing genomic features. *Bioinformatics*, *26*(6), 841–842. https://doi.org/10.1093/bioinformatics/btq033

Ricemeyer, E. S., Carroll, R. A., & Warren, W. C. (2025). Agptools: A utility suite for editing genome assemblies. *Bioinformatics*, *41*(7), btaf388. https://doi.org/10.1093/bioinformatics/btaf388

Ruhsam, M., Bell, D., Hart, M., Hollingsworth, P., Royal Botanic Garden Edinburgh Genome Acquisition Lab, Darwin Tree of Life Barcoding collective, Wellcome Sanger Institute Tree of Life programme, Wellcome Sanger Institute Scientific Operations: DNA Pipelines collective, Tree of Life Core Informatics collective, & Darwin Tree of Life Consortium. (2022). The genome sequence of the European crab apple, Malus sylvestris (L.) Mill., 1768. *Wellcome Open Research*, *7*, 296. https://doi.org/10.12688/wellcomeopenres.18645.1

Stiehler, F., Steinborn, M., Scholz, S., Dey, D., Weber, A. P. M., & Denton, A. K. (2021). Helixer: Cross-species gene annotation of large eukaryotic genomes using deep learning. *Bioinformatics*, *36*(22–23), 5291–5298. https://doi.org/10.1093/bioinformatics/btaa1044

Sun, X., Jiao, C., Schwaninger, H., Chao, C. T., Ma, Y., Duan, N., Khan, A., Ban, S., Xu, K., Cheng, L., Zhong, G.-Y., & Fei, Z. (2020). Phased diploid genome assemblies and pan-genomes provide insights into the genetic history of apple domestication. *Nature Genetics*, *52*(12), 1423–1432. https://doi.org/10.1038/s41588-020-00723-9

Tang, H., Zhang, X., Miao, C., Zhang, J., Ming, R., Schnable, J. C., Schnable, P. S., Lyons, E., & Lu, J. (2015). ALLMAPS: Robust scaffold ordering based on multiple maps. *Genome Biology*, *16*(1), 3. https://doi.org/10.1186/s13059-014-0573-1
