## Supplementary tables and figures for "Innovation within stability: balancing selection preserves multigenic S-haplotype architecture in apple"

### **Supplementary Table 1: *Malus* assemblies used in this study**

| **Haplotype** | **Species** | **Cultivar / Origin** | **Ploidy** | **Sequence length** | **Scaffold N50 (Mb)** | **Gene count** | **Protein coding genes (%)** | **Repeats (%)** | **Busco (%)** | **Chromosome 17 length (Mb)** | **References** | **Rq** |
| --- | --- | --- | --- | --- | --- | --- | --- | --- | --- | --- | --- | --- |
| h1 | *M. domestica* | cult. Golden delicious | Doubled-haploid | 656,833,032 | 37.632 | 49,762 | 51.6 | 65.4 | 96.9 | 34.749 | (Daccord et al., 2017) | Kept for subsequent analysis |
| h2 | *M. domestica* | cult. Gala | Diploid | 657,696,253 | 37.966 | 51,889 | 93.6 | 74.85 | 95.7 | 32.702 | (Sun et al., 2020) | Kept for subsequent analysis |
| h3 | *M. domestica* | cult. Gala | Diploid | 577,209,808 | 36.439 | 45,344 | 93.3 | 74.79 | 86.6 | 30.262 | (Sun et al., 2020) | Kept for subsequent analysis |
| h4 | *M. domestica* | RGTH1 | Haploid | 754,684,250 | 35.935 |  |  |  | 96.7 |  | (Tian et al., 2022) | Not kept for subsequent analysis |
| h6 | *M. domestica* | cult. Fuji | Diploid | 648,363,766 | 36.861 | 56,85 | 94.1 | 73.75 | 97.7 | 34.006 | (Li et al., 2024) | Kept for subsequent analysis |
| h7 | *M. domestica* | cult. Fuji | Diploid | 647,946,317 | 37.293 | 52,523 | 93.8 | 74.15 | 97.6 | 34.212 | (Li et al., 2024) | Kept for subsequent analysis |
| h8 | *M. domestica* | cult. Red delicious | Diploid | 625,992,033 | 37.632 | 49,554 | 93.4 | 74.16 | 97.4 | 33.496 | (Han et al., 2017) | Kept for subsequent analysis |
| h9 | *M. domestica* | cult. Honeycrisp | Diploid |  | 31.600 |  |  |  | 1.9 |  | (Khan et al., 2022) | Not kept for subsequent analysis |
| h10 | *M. domestica* | cult. Honeycrisp | Diploid | 648,227,795 | 32.8 | 51,523 | 93.6 | 74.6 | 97.6 | 33.450 | (Khan et al., 2022) | Kept for subsequent analysis |
| h11 | *M. domestica* | cult. Hanfu | Tri-  haploid | 652,502,298 | 37.139 | 51,127 | 93.6 | 75.26 | 97.6 | 33.999 | (Qin et al., 2023) | Kept for subsequent analysis |
| h12 | *M. domestica* | cult. Brown Snout | Diploid | 645,430,038 | 37.911 | 50,416 | 93.6 | 75.08 | 97.7 | 34.188 | (Könyves et al., 2022) | Kept for subsequent analysis |
| h13 | *M. domestica* | cult. Costard | Diploid | 645,735,903 | 37.048 | 50,379 | 64.1 | 75.23 | 97.6 | 33.906 | (Könyves et al., 2022) | Kept for subsequent analysis |
| h14 | *M. domestica* | cult. Bardsey Island | Diploid | 650,288,218 | 37.701 | 50,343 | 93.8 | 75.28 | 97.7 | 34.850 | (Könyves et al., 2022) | Kept for subsequent analysis |
| h15 | *M. domestica* | cult. Newtons | Diploid | 638,366,891 | 37.767 | 50,265 | 93.8 | 74.71 | 97.7 | 34.042 | (Könyves et al., 2022) | Kept for subsequent analysis |
| h16 | *M. sieversii* | Central Asia | Diploid |  | 36.622 |  |  |  | 0.7 |  | (Sun et al., 2020) | Not kept for subsequent analysis |
| h17 | *M. sieversii* | Central Asia | Diploid | 612,178,858 | 39.79 | 46,367 | 93.6 | 75.96 | 91.5 | 32.620 | (Sun et al., 2020) | Kept for subsequent analysis |
| h18 | *M. sylvestris* | Germany | Diploid |  | 35.560 |  |  |  | 0.8 |  | (Sun et al., 2020) | Not kept for subsequent analysis |
| h19 | *M. sylvestris* | Germany | Diploid | 601,478,285 | 35.227 | 49,446 | 56.8 | 74.02 | 92.7 | 31.469 | (Sun et al., 2020) | Kept for subsequent analysis |
| h20 | *M. sylvestris* | England | Diploid | 639,737,487 | 36.902 | 50,379 | 93.8 | 74.94 | 97.7 | 33.873 | (Ruhsam et al., 2022) | Kept for subsequent analysis |
| h21 | *M. domestica* | cult. Xuanina | Diploid | 644,465,789 | 29.167 | 50,762 | 57.0 | 74.57 | 97.1 | 33.655 | This study | Kept for subsequent analysis |
| h22 | *M. domestica* | cult. Xuanina | Diploid | 625,304,225 | 9.418 | 48,696 | 93.7 | 74.6 | 95.8 | 32.333 | This study | Kept for subsequent analysis |
| h23 | *M. sylvestris* | Austria | Diploid | 644,132,839 | 24.315 | 54,796 | 93.9 | 73.79 | 97.6 | 35.410 | This study | Kept for subsequent analysis |
| h24 | *M. sylvestris* | Austria | Diploid | 633,775,122 | 18.97 | 50,424 | 93.7 | 74.11 | 97.3 | 33.610 | This study | Kept for subsequent analysis |
| h25 | *M. sylvestris* | Denmark | Diploid | 638,694,078 | 24.423 | 54,188 | 93.9 | 74.06 | 97 | 34.908 | This study | Kept for subsequent analysis |
| h26 | *M. sylvestris* | Denmark | Diploid | 633,317,074 | 30.454 | 50,573 | 93.8 | 74.25 | 97.1 | 32.936 | This study | Kept for subsequent analysis |
| h27 | *M. sylvestris* | France | Diploid | 651,453,350 | 35.932 | 51,039 | 93.8 | 74.19 | 97.5 | 35.550 | This study | Kept for subsequent analysis |
| h28 | *M. sylvestris* | France | Diploid | 634,091,149 | 33.34 | 49,538 | 93.7 | 73.75 | 97.4 | 32.920 | This study | Kept for subsequent analysis |
| h29 | *M. sylvestris* | Romania | Diploid | 649,942,226 | 35.638 | 51,404 | 93.8 | 73.83 | 97.8 | 34.218 | This study | Kept for subsequent analysis |
| h30 | *M. sylvestris* | Romania | Diploid | 643,103,033 | 30.517 | 50,104 | 93.7 | 74.04 | 97.8 | 35.246 | This study | Kept for subsequent analysis |
| h31 | *M. orientalis* | Armenia | Diploid | 653,674,533 | 32.637 | 55,781 | 93.6 | 74.32 | 97.7 | 35.488 | This study | Kept for subsequent analysis |
| h32 | *M. orientalis* | Armenia | Diploid | 652,365,126 | 32.639 | 51,253 | 93.8 | 74.8 | 98 | 33.672 | This study | Kept for subsequent analysis |

### **Supplementary Table 2: S-RNase database, public accessions from *Malus* and *Pyrus* genus**

| **Accession** | **S allele** | **Species** | **Reference** |
| --- | --- | --- | --- |
| ACS94938.4 | EjS6 | *Eriobotrya japonica* | Niska et al., 2010 |
| AB002139.1 | PpS1 | *Pyrus pyrifolia* | Ishimizu et al., 1998 <https://www.ncbi.nlm.nih.gov/pubmed/9700066> |
| AB002140.1 | PpS3 | *Pyrus pyrifolia* | Ishimizu et al., 1998 <https://www.ncbi.nlm.nih.gov/pubmed/9700066> |
| AB002141.1 | PpS5 | *Pyrus pyrifolia* | Ishimizu et al., 1998 <https://www.ncbi.nlm.nih.gov/pubmed/9700066> |
| AB002142.1 | PpS6 | *Pyrus pyrifolia* | Ishimizu et al., 1998 <https://www.ncbi.nlm.nih.gov/pubmed/9700066> |
| AB002143.1 | PpS7 | *Pyrus pyrifolia* | Ishimizu et al., 1998 <https://www.ncbi.nlm.nih.gov/pubmed/9700066> |
| AB019184.1 | MdSg | *Malus domestica* | Kitahara et al., 1999 |
| AB032246.1 | MdSd | *Malus domestica* | Kitahara et al., 2000 |
| AB032247.1 | MdSh | *Malus domestica* | Kitahara et al., 2000 |
| AB035273.1 | MdSe | *Malus domestica* | Matsumoto & Kitahara, 2000 |
| AB035928.1 | MtSt | *Malus transitoria* | Matsumoto et al., 2000 |
| AB052683.1 | MdSi | *Malus domestica* | Kitahara & Matsumoto, 2002 |
| AB062100.1 | MdSz | *Malus domestica* | Kitahara & Matsumoto, 2002 |
| AB096138.1 | MtSg | *Malus transitoria* | Matsumoto et al., 2000 |
| AB104908.1 | PpS8 | *Pyrus pyrifolia* | Castillo et al., 2001 |
| AB104909.1 | PpS9 | *Pyrus pyrifolia* | Castillo et al., 2002 |
| AB236424.2 | PcSq | *Pyrus communis* | Takasaki et al., 2006 <https://www.ncbi.nlm.nih.gov/pubmed/16565843> |
| AB236425.1 | PcS1 | *Pyrus communis* | Takasaki et al., 2006 <https://www.ncbi.nlm.nih.gov/pubmed/16565843> |
| AB236426.2 | PcSr | *Pyrus communis* | Takasaki et al., 2006 <https://www.ncbi.nlm.nih.gov/pubmed/16565843> |
| AB236427.1 | PcSd | *Pyrus communis* | Takasaki et al., 2006 <https://www.ncbi.nlm.nih.gov/pubmed/16565843> |
| AB236428.1 | PcSe | *Pyrus communis* | Takasaki et al., 2006 <https://www.ncbi.nlm.nih.gov/pubmed/16565843> |
| AB236429.1 | PcSb | *Pyrus communis* | Takasaki et al., 2006 <https://www.ncbi.nlm.nih.gov/pubmed/16565843> |
| AB236430.1 | PcSa | *Pyrus communis* | Takasaki et al., 2006 <https://www.ncbi.nlm.nih.gov/pubmed/16565843> |
| AB236431.1 | PcSa | *Pyrus communis* | Takasaki et al., 2006 <https://www.ncbi.nlm.nih.gov/pubmed/16565843> |
| AB236432.1 | PcSk | *Pyrus communis* | Takasaki et al., 2006 <https://www.ncbi.nlm.nih.gov/pubmed/16565843> |
| AB258359.1 | PcSc | *Pyrus communis* | Moriya et al., unpublished |
| AB258360.1 | PcSg | *Pyrus communis* | Moriya et al., unpublished |
| AB258361.1 | PcSi | *Pyrus communis* | Moriya et al., unpublished |
| AB258362.1 | PcSm | *Pyrus communis* | Moriya et al., unpublished |
| AB258363.1 | PcSn | *Pyrus communis* | Moriya et al., unpublished |
| AB258364.1 | PcSp | *Pyrus communis* | Moriya et al., unpublished |
| AB258365.1 | PcSs | *Pyrus communis* | Moriya et al., unpublished |
| AB258366.1 | PcSt | *Pyrus communis* | Moriya et al., unpublished |
| AB284262.1 | PpSk | *Pyrus pyrifolia* | Kim et al., 2007 |
| AB426604.1 | PpS12 | *Pyrus pyrifolia* | Okada et al., unpublished |
| AB426606.1 | PpSk | *Pyrus pyrifolia* | Okada et al., unpublished |
| AB540121.1 | MsS33 | *Malus sieversii* | Matsumoto et al., 2010 |
| AB540122.1 | MdS34 | *Malus domestica* | Matsumoto et al., 2010 |
| AB731592.1 | PcS25 | *Pyrus communis* | Takasaki-Yasuda et al., unpublished |
| AB779646.1 | PcS0 | *Pyrus communis* | Takasaki-Yasuda et al., unpublished |
| AB779647.1 | PcS00 | *Pyrus communis* | Takasaki-Yasuda et al., unpublished |
| AF016918.1 | MdS26 | *Malus domestica* | Verdoodt et al., unpublished |
| AF016919.1 | MdS27 | *Malus domestica* | Verdoodt et al., unpublished |
| AF016920.1 | MdS24 | *Malus domestica* | Verdoodt et al., unpublished |
| AF327221.1 | MdS10 | *Malus domestica* | Van Nerum et al. unpublished |
| AF327222.1 | MdS27b | *Malus domestica* | Van Nerum et al. unpublished |
| AF327223.1 | MdS4 | *Malus domestica* | Van Nerum et al. unpublished |
| AY249428.2 | PpS13 | *Pyrus pyrifolia* | Tan et al., unpublished |
| AY250989.3 | PbS21 | *Pyrus bretschneideri* | Tan et al., unpublished |
| D49527.1 | PpS0000 | *Pyrus pyrifolia* | Norioka et al., 1995 |
| D49528.1 | PpS000 | *Pyrus pyrifolia* | Norioka et al., 1995 |
| D50836.1 | MdSc | *Malus domestica* | Sassa et al., 1996 |
| D50837.1 | MdSf | *Malus domestica* | Sassa et al., 1996 |
| D88282.1 | PsS5 | *Pyrus serotina* | Sassa et Hirano., 1997 |
| DQ135990.1 | MdS31 | *Malus domestica* | Kim et al., unpublished |
| DQ135991.1 | MdS32 | *Malus domestica* | Kim et al., unpublished |
| DQ224344.1 | PbS35 | *Pyrus bretschneideri* | Yang & Tan, unpublished |
| DQ414812.1 | PbS13 | *Pyrus bretschneideri* | Tan et al., unpublished |
| DQ414813.1 | PbS34 | *Pyrus bretschneideri* | Tan et al., unpublished |
| DQ494532.1 | PbS21 | *Pyrus bretschneideri* | Zhang et al., 2006 |
| DQ494676.1 | PbS34 | *Pyrus bretschneideri* | Zhang et al., 2006 |
| DQ839240.1 | PuS35 | *Pyrus ussuriensis* | Wuyun et al., unpublished |
| DQ991388.1 | PbS16 | *Pyrus bretschneideri* | Tan et al., unpublished |
| DQ995285.1 | PbS39 | *Pyrus bretschneideri* | Tan et al., unpublished |
| EF566872.1 | PsiS28 | *Pyrus sinkiangensis* | Zhang et al., unpublished |
| EF643630.1 | PpS15 | *Pyrus pyrifolia* | Zhang et al., unpublished |
| EF643631.2 | PbS38 | *Pyrus bretschneideri* | Tan et al., unpublished |
| EF643635.1 | PbS16 | *Pyrus bretschneideri* | Tan et al., unpublished |
| EF643636.1 | PbS18 | *Pyrus bretschneideri* | Tan et al., unpublished |
| EF643638.1 | PbS19 | *Pyrus bretschneideri* | Tan et al., unpublished |
| EF643639.1 | PsiS22 | *Pyrus sinkiangensis* | Tan et al., unpublished |
| EF643640.1 | PbS27 | *Pyrus bretschneideri* | Tan et al., unpublished |
| EF643641.1 | PuS30 | *Pyrus ussuriensis* | Tan et al., unpublished |
| EF689006.1 | PuS42 | *Pyrus ussuriensis* | Tan et al., unpublished |
| EF689007.1 | PbS42 | *Pyrus bretschneideri* | Tan et al., unpublished |
| EF689008.1 | PbS22 | *Pyrus bretschneideri* | Tan et al., unpublished |
| EU081888.1 | PbS26 | *Pyrus bretschneideri* | Tan et al., unpublished |
| EU081889.1 | PbS12 | *Pyrus bretschneideri* | Tan et al., unpublished |
| EU101462.1 | PbS29 | *Pyrus bretschneideri* | Tan et al., unpublished |
| EU101463.1 | PbS26 | *Pyrus bretschneideri* | Tan et al., unpublished |
| EU101464.1 | PuS40 | *Pyrus ussuriensis* | Tan et al., unpublished |
| EU101465.1 | PpS5a | *Pyrus pyrifolia* | Tan et al., unpublished |
| EU101466.1 | PbS17 | *Pyrus bretschneideri* | Tan et al., unpublished |
| EU117115.1 | PpS12 | *Pyrus pyrifolia* | Tan et al., unpublished |
| EU336979.1 | PuS32 | *Pyrus ussuriensis* | Tan et al., unpublished |
| EU336980.1 | PbS39 | *Pyrus bretschneideri* | Tan et al., unpublished |
| EU360894.1 | PbS20 | *Pyrus bretschneideri* | Tan et al., unpublished |
| EU375364.1 | PbS28 | *Pyrus bretschneideri* | Tan et al., unpublished |
| EU477839.1 | PcS21 | *Pyrus communis* | Sanzol, <https://www.ncbi.nlm.nih.gov/pubmed/19096853> |
| EU477840.1 | PcS21 | *Pyrus communis* | Sanzol, <https://www.ncbi.nlm.nih.gov/pubmed/19096853> |
| FJ943264.1 | MspecS1 | *Malus spectabilis* | Zhang et al., unpublished |
| FJ943266.1 | MspecS2 | *Malus spectabilis* | Zhang et al., unpublished |
| FJ943268.1 | MspecS3 | *Malus spectabilis* | Zhang et al., unpublished |
| FJ943270.1 | MspecS4 | *Malus spectabilis* | Zhang et al., unpublished |
| FJ943272.1 | MspecS5 | *Malus spectabilis* | Zhang et al., unpublished |
| FJ946628.1 | PpS46 | *Pyrus pyrifolia* | Tan et al., unpublished |
| GQ180466.1 | PcS4 | *Pyrus communis* | Sanzol, <https://www.ncbi.nlm.nih.gov/pubmed/20701704> |
| KF468727.1 | MsiS1 | *Malus sikkimensis* | Gu, unpublished |
| KF468730.1 | MsiS7 | *Malus sikkimensis* | Gu, unpublished |
| KF588567.1 | PcS126 | *Pyrus communis* | Nikzad Gharehaghaji et al., unpublished |
| U12199.1 | MdS2 | *Malus domestica* | Broothaerts et al., 1995 |
| U12200.1 | MdS3 | *Malus domestica* | Broothaerts et al., 1995 |
| U19793.1 | MdS9 | *Malus domestica* | Broothaerts et al., 1995 |

### **Supplementary Table 3: S-loci coordinates, predicted SFBB number and S-alleles in studied haplotypes**

| **Haplotype** | **Species** | **Cultivar/ origin** | **S-loci start on chr17** | **S-loci end on chr17** | **S-loci**  **length** | **Predicted S-loci gene number** | **Predicted SFBBs number (complete sequences)** | **S-Rnase ID** | **S-RNase length** | **Predicted S-RNase allele** |
| --- | --- | --- | --- | --- | --- | --- | --- | --- | --- | --- |
| h14 | *M. domestica* | Bardsey Island | 30701244 | 32097319 | 1396076 | 39 | 17 | h14_chr17_3.1 | 834 | MdS2 |
| h12 | *M. domestica* | Brown snout | 29604507 | 31323943 | 1719437 | 43 | 18 | h12_chr17_001110.1 | 1154 | MspecS4 |
| h13 | *M. domestica* | Costard | 29699373 | 31095434 | 1396062 | 38 | 17 | h13_chr17_3.1 | 834 | MdS2 |
| h7 | *M. domestica* | Fuji | 30197526 | 31518020 | 1320495 | 40 | 17 | h7_chr17_6.1 | 830 | MdS9 |
| h6 | *M. domestica* | Fuji | 29893470 | 31195680 | 1302211 | 48 | 17 | h6_chr17_001448.1 | 1295 | MdSf |
| h3 | *M. domestica* | Gala | 26135286 | 27539957 | 1404672 | 51 | 17 | h3_chr17_3.1 | 834 | MdS2 |
| h2 | *M. domestica* | Gala | 28665960 | 29985017 | 1319058 | 49 | 17 | h2_chr17_001456.1 | 2061 | MdS5 |
| h1 | *M. domestica* | Golden delicious double haploid (GDDH13) | 30032275 | 31732190 | 1699916 | 33 | 16 | h1_chr17_001308.1 | 2322 | MdS3 |
| h11 | *M. domestica* | Hanfu | 29830584 | 31150956 | 1320373 | 42 | 18 | h11_chr17_8.1 | 831 | MdS9 |
| h10 | *M. domestica* | Honeycrisp | 29310889 | 30706999 | 1396111 | 39 | 17 | h10_chr17_4.1 | 834 | MdS2 |
| h15 | *M. domestica* | Newtons | 29920233 | 31222329 | 1302097 | 43 | 18 | h15_chr17_001441.1 | 1309 | MdSf |
| h8 | *M. domestica* | Red delicious | 29883495 | 30810778 | 927284 | 48 | 17 | h8_chr17_3.1 | 805 | MdSe |
| h21 | *M. domestica* | Xuanina | 29544619 | 30982290 | 1437672 | 42 | 18 | h21_chr17_001396.1 | 1172 | MdS26 |
| h22 | *M. domestica* | Xuanina | 28259688 | 29525297 | 1265610 | 40 | 18 | h22_chr17_001349.1 | 1037 | MdS4 |
| h32 | *M. orientalis* | Armenia | 29847786 | 30941904 | 1094119 | 36 | 19 | h32_chr17_001379.1 | 1059 | MdS9 |
| h31 | *M. orientalis* | Armenia | 31699461 | 32754781 | 1055321 | 45 | 19 | h31_chr17_5.1 | 848 | NewC |
| h17 | *M. sieversii* | Kazakstan | 28636332 | 29834722 | 1198391 | 36 | 17 | h17_chr17_3.1 | 834 | PbS13 |
| h20 | *M. sylvestris* |  | 30157545 | 31217444 | 1059900 | 37 | 16 | h20_chr17_001396.1 | 1063 | MdS4 |
| h19 | *M. sylvestris* |  | 27161358 | 28640677 | 1479320 | 39 | 20 | h19_chr17_4.1 | 633 | PpS1 |
| h23 | *M. sylvestris* | Austria | 30930378 | 32432552 | 1502175 | 47 | 16 | h23_chr17_2.1 | 834 | MdS2 |
| h24 | *M. sylvestris* | Austria | 29911385 | 30929757 | 1018373 | 40 | 14 | h24_chr17_001105.1 | 1131 | NewA |
| h26 | *M. sylvestris* | Denmark | 29352325 | 30455556 | 1103232 | 38 | 17 | h26_chr17_001363.1 | 1967 | MdS5 |
| h25 | *M. sylvestris* | Denmark | 30768546 | 32202276 | 1433731 | 42 | 18 | h25_chr17_2.1 | 845 | NewD |
| h28 | *M. sylvestris* | France | 28763989 | 30262854 | 1498866 | 37 | 17 | h28_chr17_001351.1 | 1508 | PbS42 |
| h27 | *M. sylvestris* | France | 31333056 | 32800349 | 1467294 | 41 | 19 | h27_chr17_3.1 | 853 | MdSe |
| h29 | *M. sylvestris* | Romania | 29817762 | 31260259 | 1442498 | 49 | 19 | h29_chr17_4.1 | 1830 | NewE |
| h30 | *M. sylvestris* | Romania | 30593311 | 32469098 | 1875788 | 47 | 17 | h30_chr17_2.1 | 846 | MspecS1 |

**Supplementary Table 4: Pairwise dN, dS and ω (dN/dS) among 27 S-RNase haplotype CDS sequences (chr17), same-allele vs different-allele comparisons.** Pairwise dN, dS and ω = dN/dS were estimated with yn00 (Yang & Nielsen 2000, PAML v4.9) for all pairwise comparisons among 27 SRNase haplotype coding sequences. pair_type = intra: both haplotypes share the same S-allele (group column: MdS2, MdS4, MdS5, MdS9, MdSe, MdSf); pair_type = inter: different S-alleles (group = NA). Intra-allelic pairs were aligned at the amino-acid level (MUSCLE) and back-translated to codons (PAL2NAL) to properly handle real in-frame indels before dN/dS estimation. omega = 99.0 (with dN = dS ≈ 0) marks pairs with no detected substitutions, where the ratio is undefined.

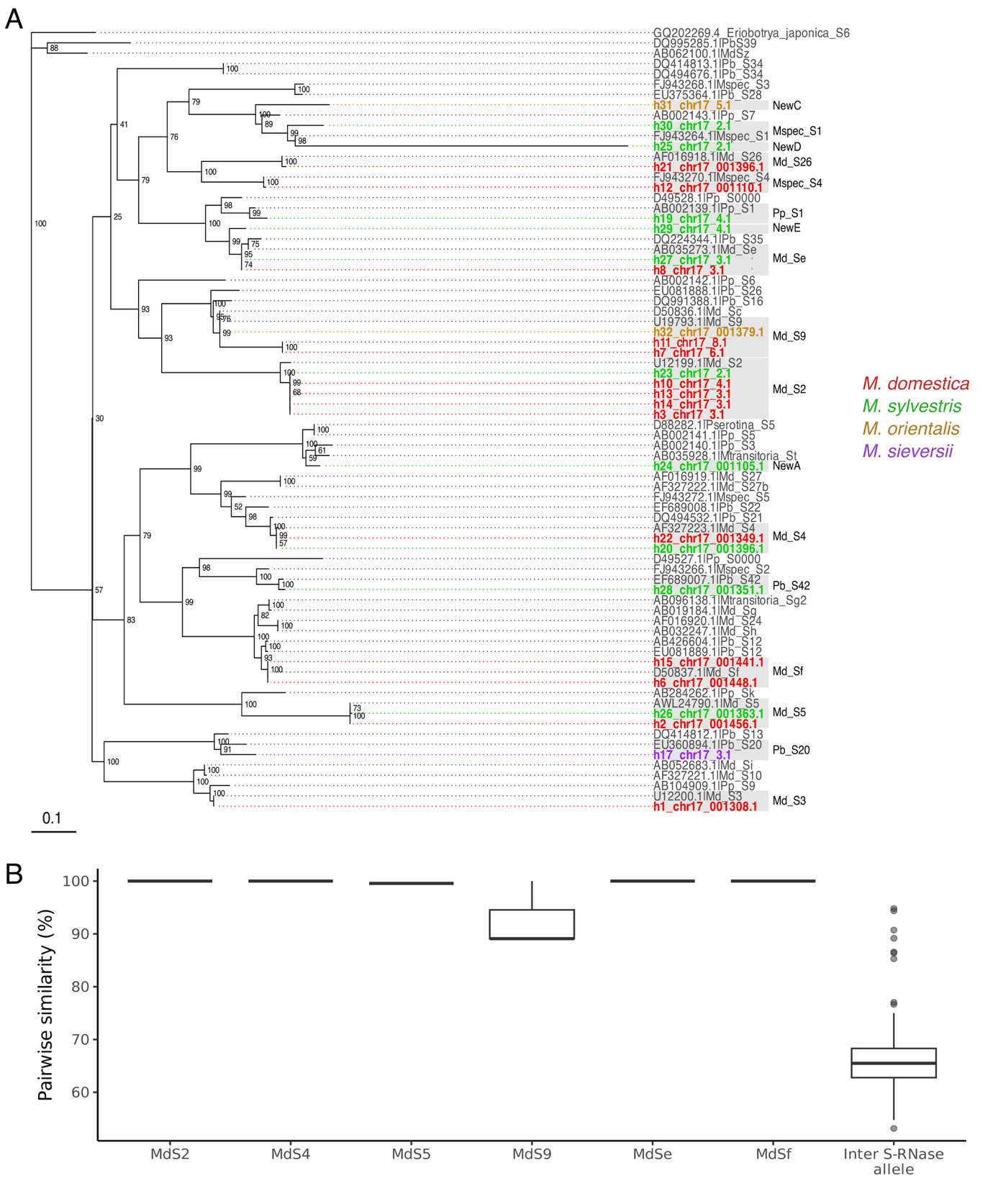

**Supplementary Fig. 1: Phylogeny and similarity patterns of *Malus* S-RNases.** (A) Maximum-likelihood phylogeny of S-RNase protein sequences including both annotated haplotype sequences from the present dataset and publicly available S-RNases. Species of origin are indicated: *Md* = *Malus domestica*; *Mspec* = *M. spectabilis*; *Msiev* = *M. sieversii*; *Pb* = *Pyrus breitschneideri*; *Pc* = *P. communis*; *Pp* = *P. pyrifolia*; *Pu* = *P. ussuriensis*. (B) Distribution of pairwise protein sequence similarities among S-RNases, computed in R based on amino acid identity (1 – distance). Boxplots show similarity values for S-RNases belonging to the same S-allele (intra-allelic comparisons) versus those belonging to different alleles (inter-allelic comparisons).

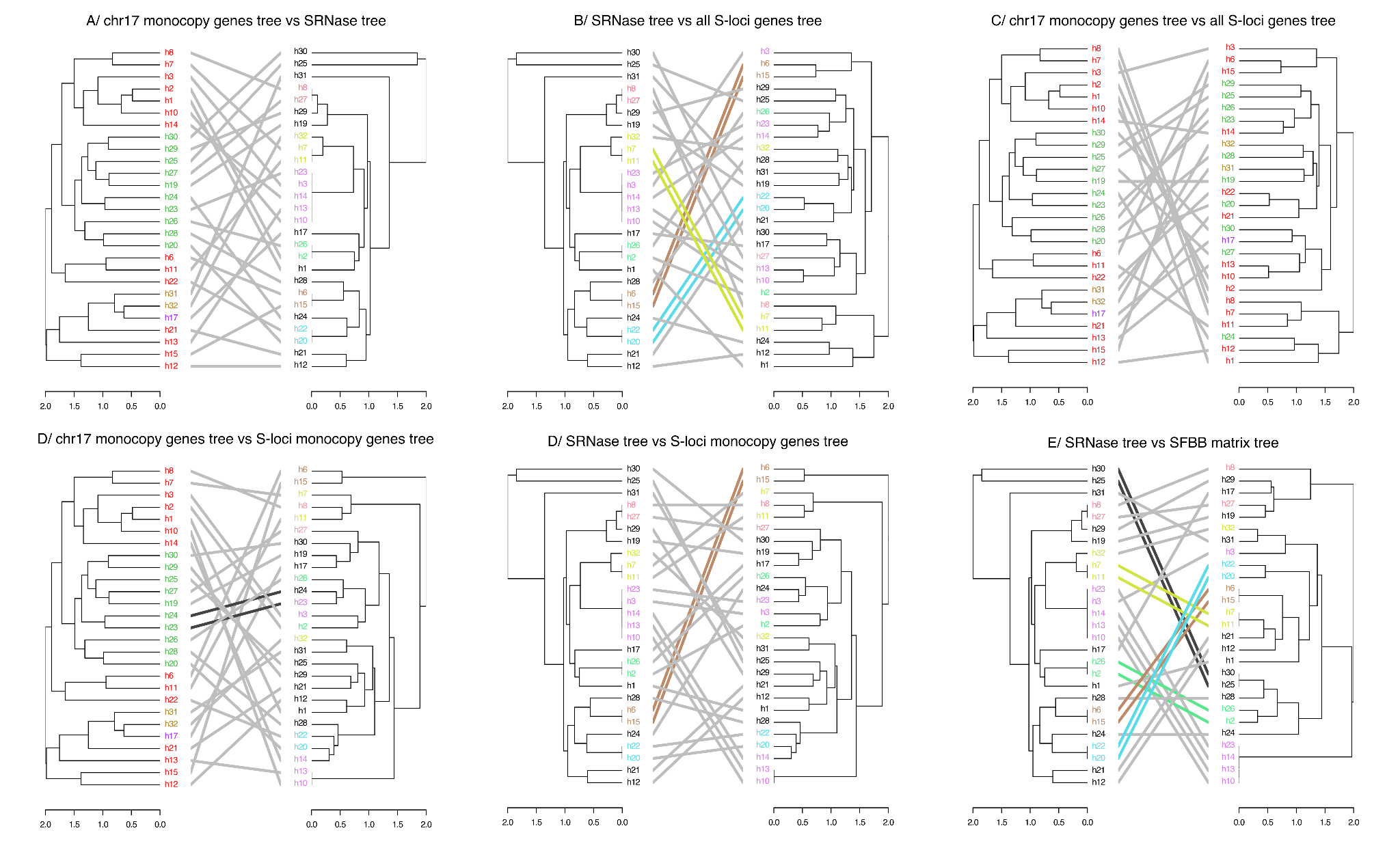

### **Supplementary Fig. 2: Tanglegram comparisons of phylogenetic trees related to S-locus components in apple trees.** Tanglegrams illustrate topological comparisons between hierarchical clustering trees derived from different genomic features: **(A)** Species tree vs. *S-RNase* gene tree ; **(B)** *S-RNase* gene tree vs. tree based on all S-locus genes (including mono- and multi-copy genes) ; **(C)** Species tree vs. all S-loci gene tree **(D)** Species tree vs. tree based on S-locus single-copy genes only ; **(E)** *S-RNase* gene tree vs. S-locus single-copy genes tree ; **(F)** *S-RNase* gene tree vs. tree derived from presence/absence matrix of SFBB genes. Each line links identical haplotypes (h1 to h32) between the trees being compared. Colored connectors highlight haplotypes exhibiting strong or consistent topological congruence across comparisons. Horizontal axes indicate phylogenetic distances. Tree topologies were computed using hierarchical clustering based on pairwise genetic distances. Two independent color schemes are used to facilitate interpretation: 1. Species colors: Red = *M. domestica*; Green = *M. sylvestris*; Yellow = *M. orientalis*; Purple = *M. sieversii*. 2. S-allele colors: Colors correspond to specific S-alleles: Pink: MdS2; Brown: MdSf; Light green: MdS5; Blue: MdS4; Light yellow: MdS9; Salmon: MdSe.

### **Supplementary Table 5: SFBB database, public accessions from *Malus* and *Pyrus* (excel file).**

### **Supplementary Table 6: Expected and observed S-RNase alleles in *M. domestica* S-haplotypes from our dataset**

| **Cultivar** | **Expected S allele (references)** | **Observed S alleles** |
| --- | --- | --- |
| Golden delicious | MdS2/MdS3 (Schneider et al., 2001) | h1 (MdS3) |
| Fuji | MdS1/MdS9 (Kasajima et al., 2017) or MdSf (Broothaerts, 2003; Matsumoto et al., 1999) or MdSc/MdSf (Sassa et al., 1994) | h6 (MdSf) / h7 (MdS9) |
| Honeycrisp | MdS2/MdS24 (Sakurai et al., 2000) | h10 (MdS2) |
| Gala | MdS2/MdS5 (Janssens et al., 1995) | h2 (?) / h3 (MdS2) |
| Red delicious | MdS9/MdS28 (Matsumoto et al., 2003) MdSe = MdS28 (Kasajima et al., 2017) | h8 (?) |
| Newtons | Unknown | h15 (MdSf) |
| Brown snout | Unknown | h12 (MspecS4) |
| Hanfu | Unknown | h11 (MdS9) |
| Costard | Unknown | h13 (MdS3) |
| Bardsey Island | Unknown | h14 (MdS3) |

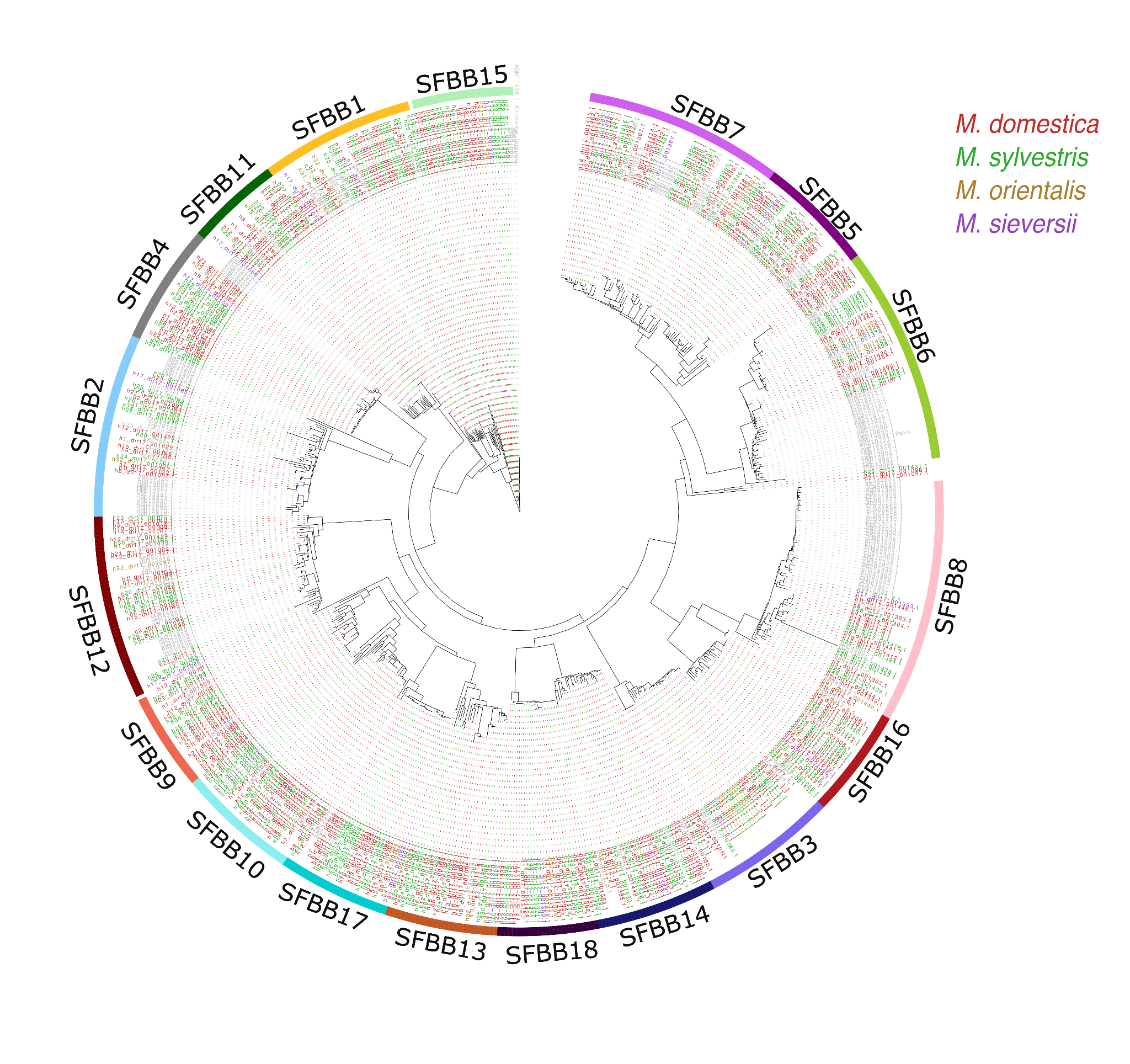

### **Supplementary Fig. 3: SFBB genes phylogeny and families determined using a combination of classification methods.** Phylogenetic tree based on the alignment of SFBB protein sequences from *Malus* S-haplotypes included in this study, along with publicly available SFBB sequences. Protein sequences were aligned using Clustal Omega (Sievers & Higgins, 2014), and the tree was inferred with IQ-TREE (Minh et al., 2020) using 1000 bootstrap replicates. Hierarchical clustering using the Ward method was combined with Orthogroups identification using Orthofinder (Emms & Kelly, 2019) to determine SFBB families, which were then compared with those from publicly available SFBB sequences. Tip labels are color-coded according to the species origin of the S-haplotype from which each SFBB sequence was derived.

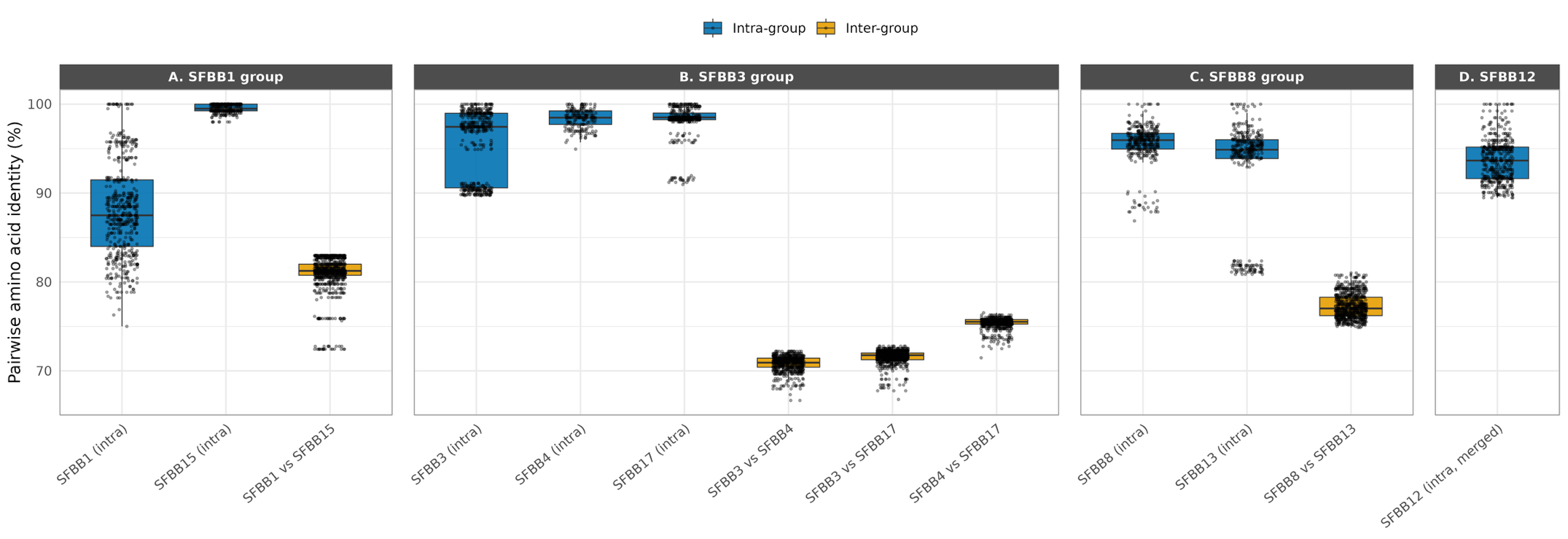
**Supplementary Fig. 4: Pairwise similarity distribution of SFBB gene groups before and after proposed reclassification.** Boxplots display the pairwise similarity values among sequences within and between different SFBB groups. (A) The SFBB1 group and its proposed subdivision into SFBB1 and SFBB15. (B) The SFBB3 group and its proposed subdivision into SFBB3 and SFBB4. (C) The SFBB8 group and its proposed subdivision into SFBB8 and SFBB13. (D) SFBB4 group, resulting from the merging of two previously separate families (former SFBB4 and SFBB12), now SFBB12. For (D), the high pairwise similarity observed among sequences supports the merging of these two families into a single family. See Supplementary Note 1 for methods.

### **Supplementary Table 7: Original and updated SFBB families, based on Ward hierarchical clustering, Orthofinder results, and phylogenetic relationships among SFBB protein sequences.**

| **Original SFBB families** | **Updated SFBB families proposition** | **Supported by** |
| --- | --- | --- |
| SFBB1 | SFBB1 | Ward, Orthofinder, phylogeny, identity percentages |
|  | SFBB15 | Ward, Orthofinder, phylogeny, identity percentages |
| SFBB2 | SFBB2 | Ward, Orthofinder, phylogeny, identity percentages |
| SFBB3 | SFBB3 | Ward, Orthofinder, phylogeny, identity percentages |
|  | SFBB4 | Ward, Orthofinder, phylogeny, identity percentages |
|  | SFBB17 | Ward, Orthofinder, phylogeny, identity percentages |
| SFBB8 | SFBB8 | Ward, phylogeny, identity percentages |
|  | SFBB13 | Ward, phylogeny, identity percentages |
| SFBB4 | SFBB12 | Ward, phylogeny, identity percentages |
| SFBB12 |  |  |
| SFBB9 | ~SFBB9 | Ward, Orthofinder, phylogeny, identity percentages |
| SFBB10 | ~SFBB10 | Ward, Orthofinder, phylogeny, identity percentages |
| SFBB11 | SFBB11 | Ward, Orthofinder, phylogeny, identity percentages |
| SFBB5 | SFBB5 | Ward, phylogeny, identity percentages |
| SFBB6 | SFBB6 | Ward, phylogeny, identity percentages |
| SFBB7 | SFBB7 | Ward, phylogeny, identity percentages |
| – | SFBB14 | Ward, Orthofinder, phylogeny, identity percentages |
| – | SFBB16 | Ward, Orthofinder, phylogeny, identity percentages |
| – | SFBB18 | Ward, Orthofinder, phylogeny, identity percentages |

### **Supplementary Table 8: SFBB from assembly set, coordinates and families (excel file).**

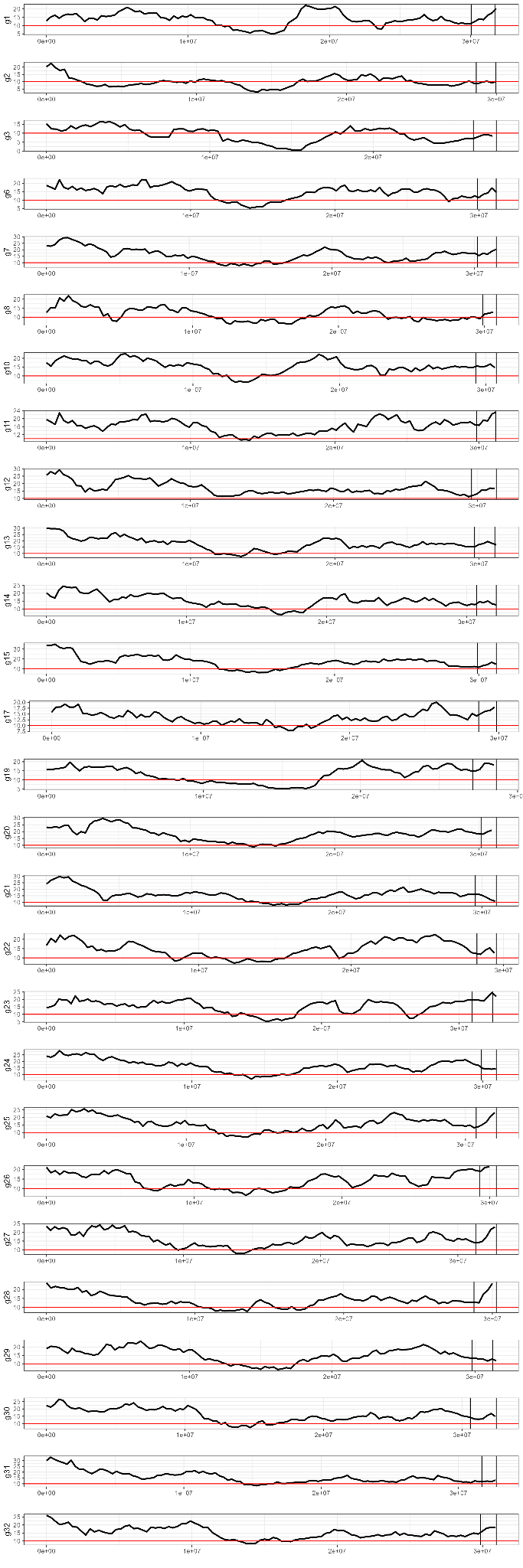

**Supplementary Fig. 5: Variation of the LTR Assembly Index (LAI) along chromosome 17 for different *Malus* assemblies.** Each panel corresponds to a distinct assembly. Black lines represent local LAI values, while the horizontal red line indicates the quality threshold. The S-locus region is delimited by the two vertical bars at the right end of each plot.

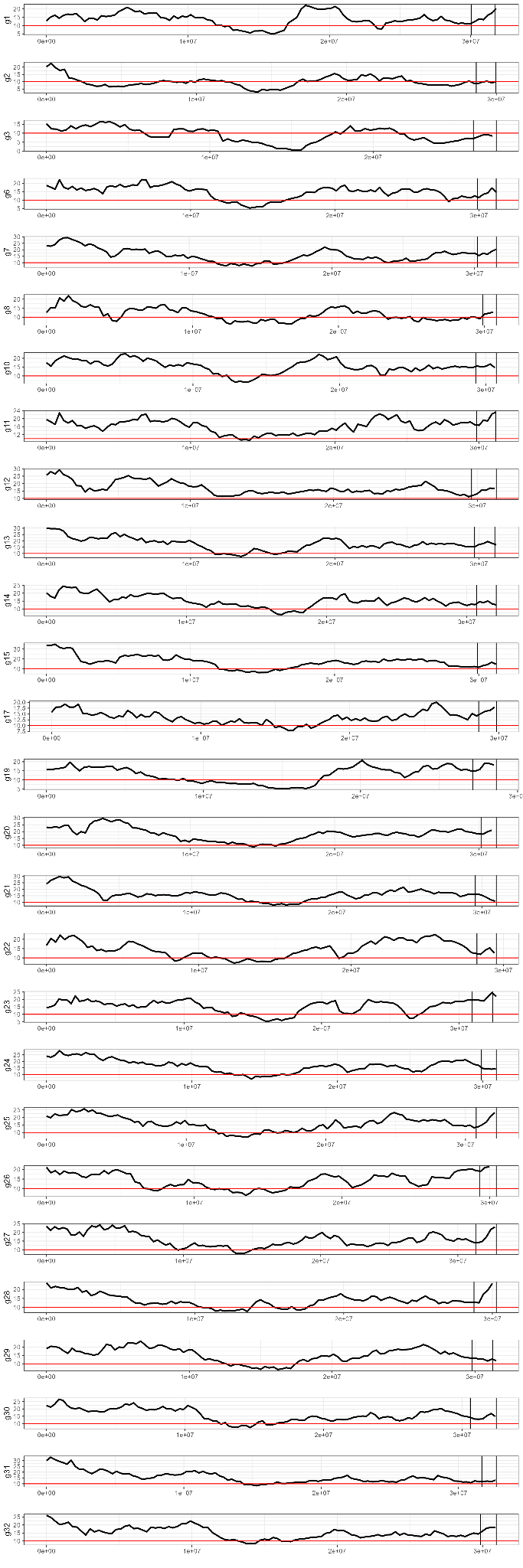

**Supplementary Fig. 5 (continued)**

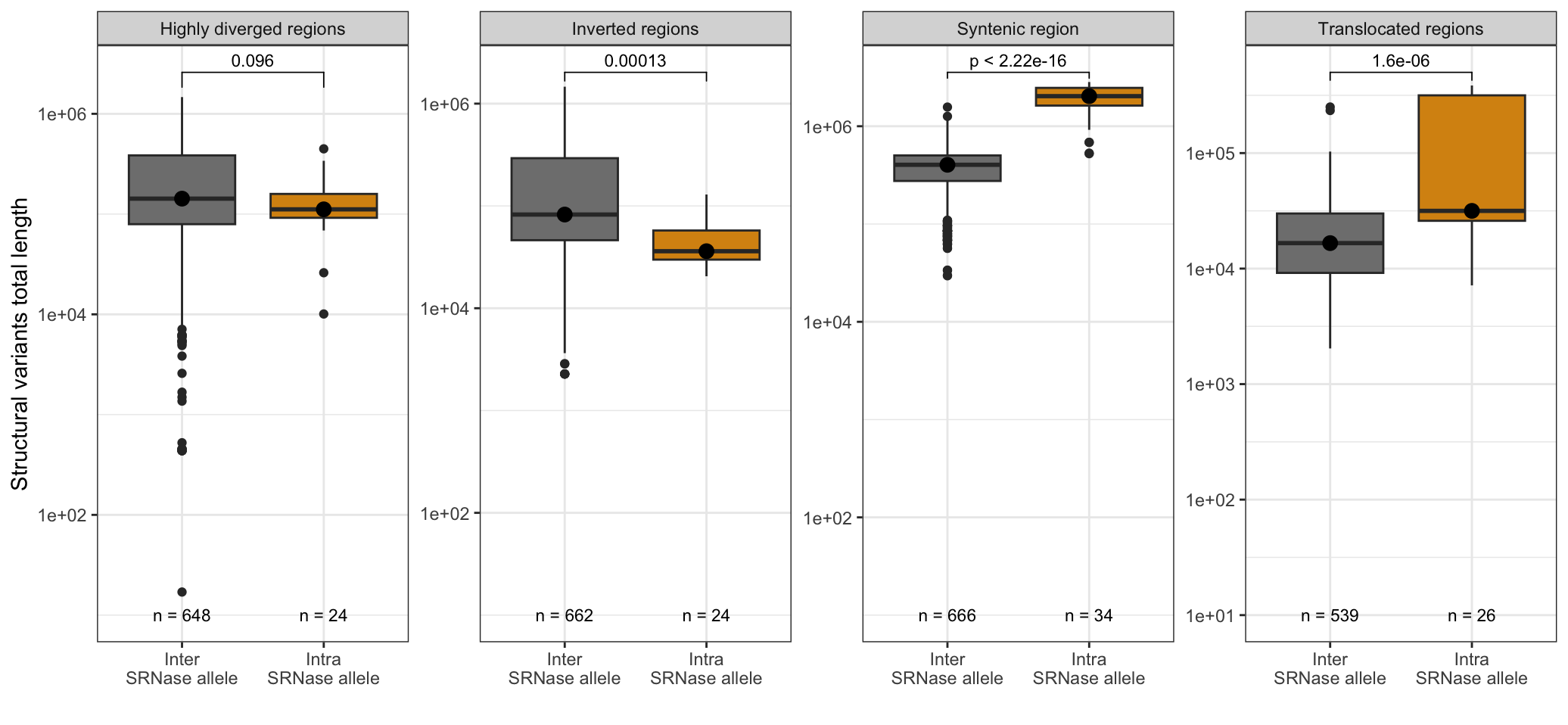

**Supplementary Fig. 6: Structural variant and syntenic regions size distribution between S-locus haplotypes sharing the same or different S-RNase alleles.** Boxplots showing the distribution of region sizes (in bp, log scale) for different types of structural variants detected in pairwise alignments of S-locus haplotypes, categorized by whether the two haplotypes share the same S-RNase allele (*intra S-RNase allele*, orange) or not (*inter S-RNase allele*, gray). Structural variant classes include inverted, translocated, and highly diverged regions, as defined by SyRI. For each category, the median region size is indicated along with results from a Wilcoxon–Mann–Whitney test comparing intra vs. inter S-RNase groups. *W* statistics, *p*-values are reported above each plot. Syntenic regions are significantly larger between haplotypes with the same S-RNase allele (*p-value* < 2.2e−16), whereas inverted regions are significantly larger (*p-value* = 1.3e-4) in comparisons between different S-RNases, suggesting increased structural divergence between non-homologous S-haplotypes.

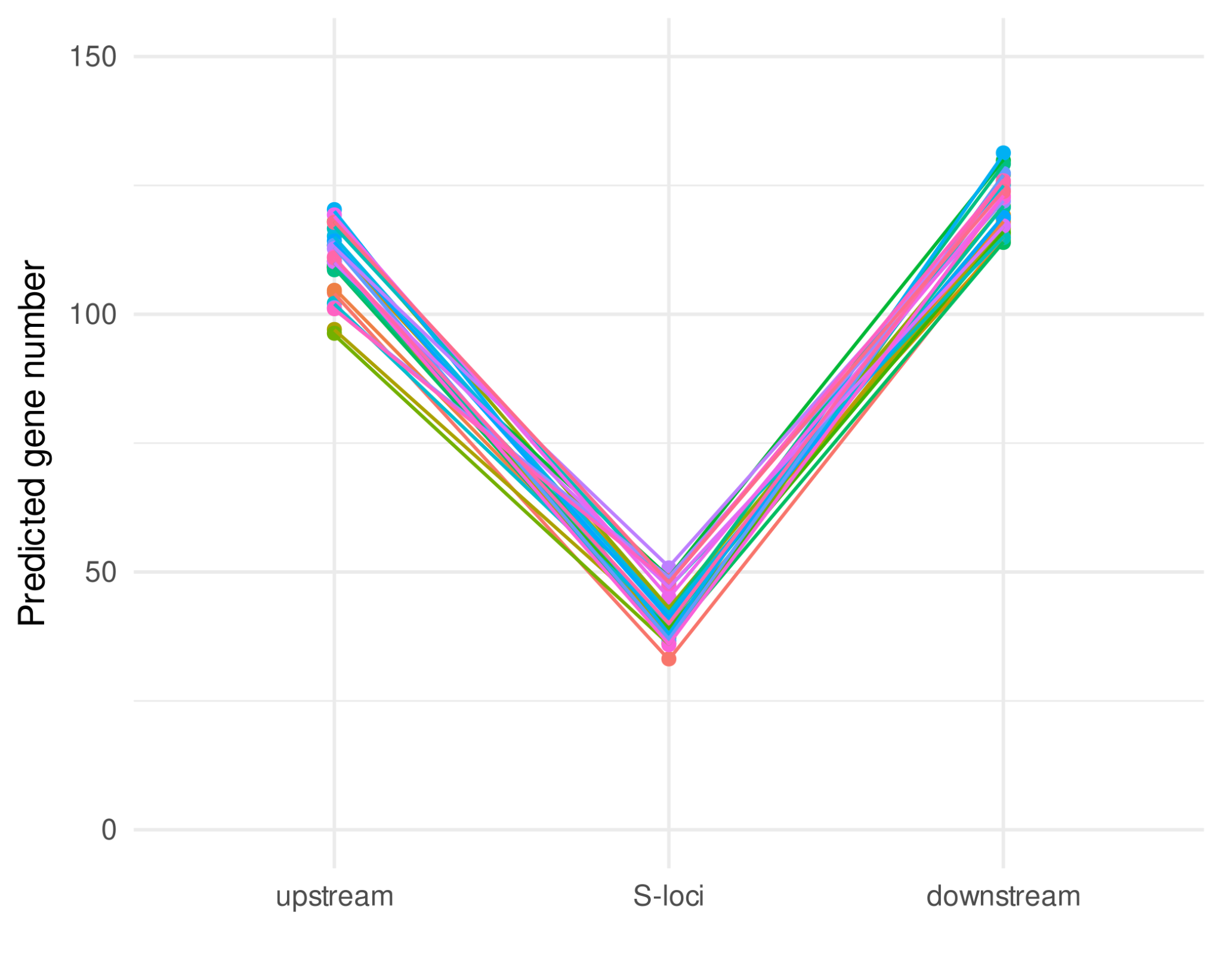

**Supplementary Fig. 7: Predicted gene number inside S-loci and in regions of the same length (1.35 Mb) upstream and downstream.** Each point is one haplotype region, and regions from the same haplotypes have the same color and are linked by the same color line.

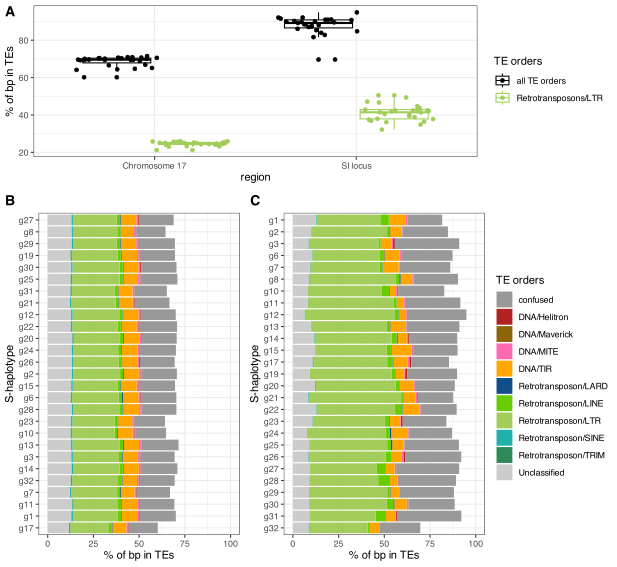

### **Supplementary Fig. 8: Chromosome 17 and S-loci TEs content in *Malus* haplotypes.** Base percentage in TE appears higher in S-loci than in chromosome 17 (A, black points), and the difference is more pronounced for LTR retrotransposons (A, green points). TEs families proportion appears conserved in different chromosome 17 (B) and less conserved across SI loci, even with the same S allele (C). The confused category corresponds to TEs for which the classification assigned multiple possible families, with no clear preference for one family over another.

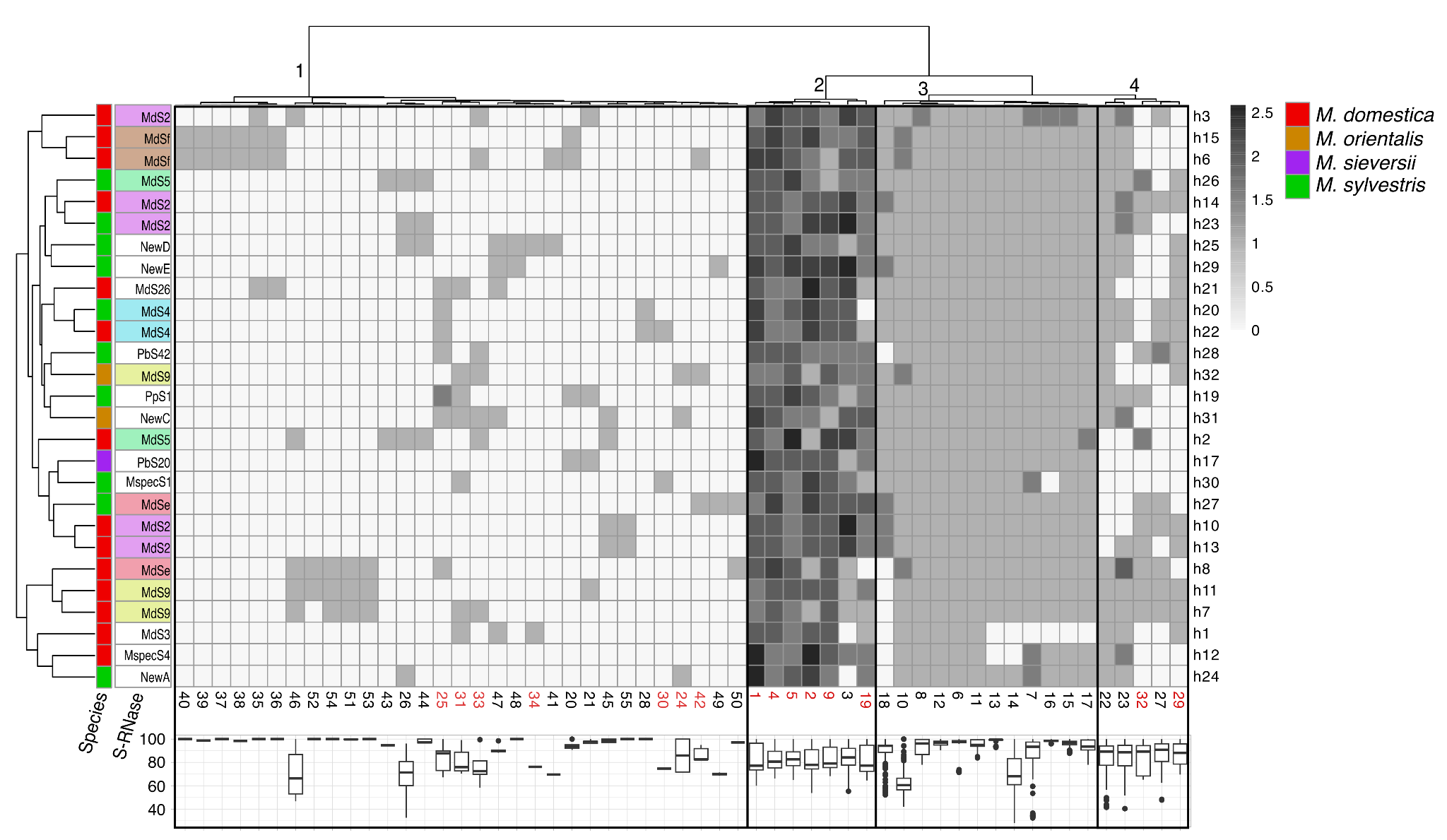

**Supplementary Fig. 9: Heatmap showing the number of genes from each synteny cluster present in individual S-haplotypes.** Each column represents a gene cluster inferred by syntenet (Almeida-Silva et al., 2023) from the *S*-locus region. Each row corresponds to an *S*-haplotype from one of four *Malus* species (*M. domestica*, *M. sylvestris*, *M. sieversii*, *M. orientalis*), annotated by species (left color strip) and *S*-allele (right color strip). The grayscale indicates the log-transformed gene count (log₂(n + 1)) for each cluster within each *S*-haplotype. Cluster IDs are shown below each column; those highlighted in red contain known SFBB genes, which are involved in the self-incompatibility system. Both *S*-haplotypes and gene clusters were hierarchically clustered based on gene content similarity. Based on their distribution across S-haplotypes, gene clusters were grouped into four categories. Group 2 includes clusters 19, 3, 9, 2, 4, 1, and 5, which are present in all S-haplotypes in multiple copies. This group is composed mostly of SFBB gene clusters, except for cluster 3, which contains genes encoding a serine/threonine protein kinase and transcription factors. Group 3 contains clusters 15, 8, 16, 17, 7, 14, 13, 11, 6, 12, 10, and 18, which typically include a single gene copy per haplotype. These include the S-RNase gene (cluster 10), as well as genes encoding a DNA ligase, a C2H2-type zinc finger, and a coatomer subunit. Groups 1 and 4 consist of clusters that are not consistently present across all haplotypes. Clusters in group 1 are rarer than those in group 4. Notably, some SFBB gene clusters are also found in these two groups, indicating variable copy number and presence across haplotypes. The full gene composition of each cluster is provided in Supplementary Table 8.

**Supplementary Table 9:** **Gene content of S-locus clusters across the 27 Malus haplotypes.** For each cluster group and individual cluster, the table lists the gene identifiers present, their putative functions (e.g., SFBB family membership or other annotated domains), and the corresponding S-haplotypes when available (Excel file).

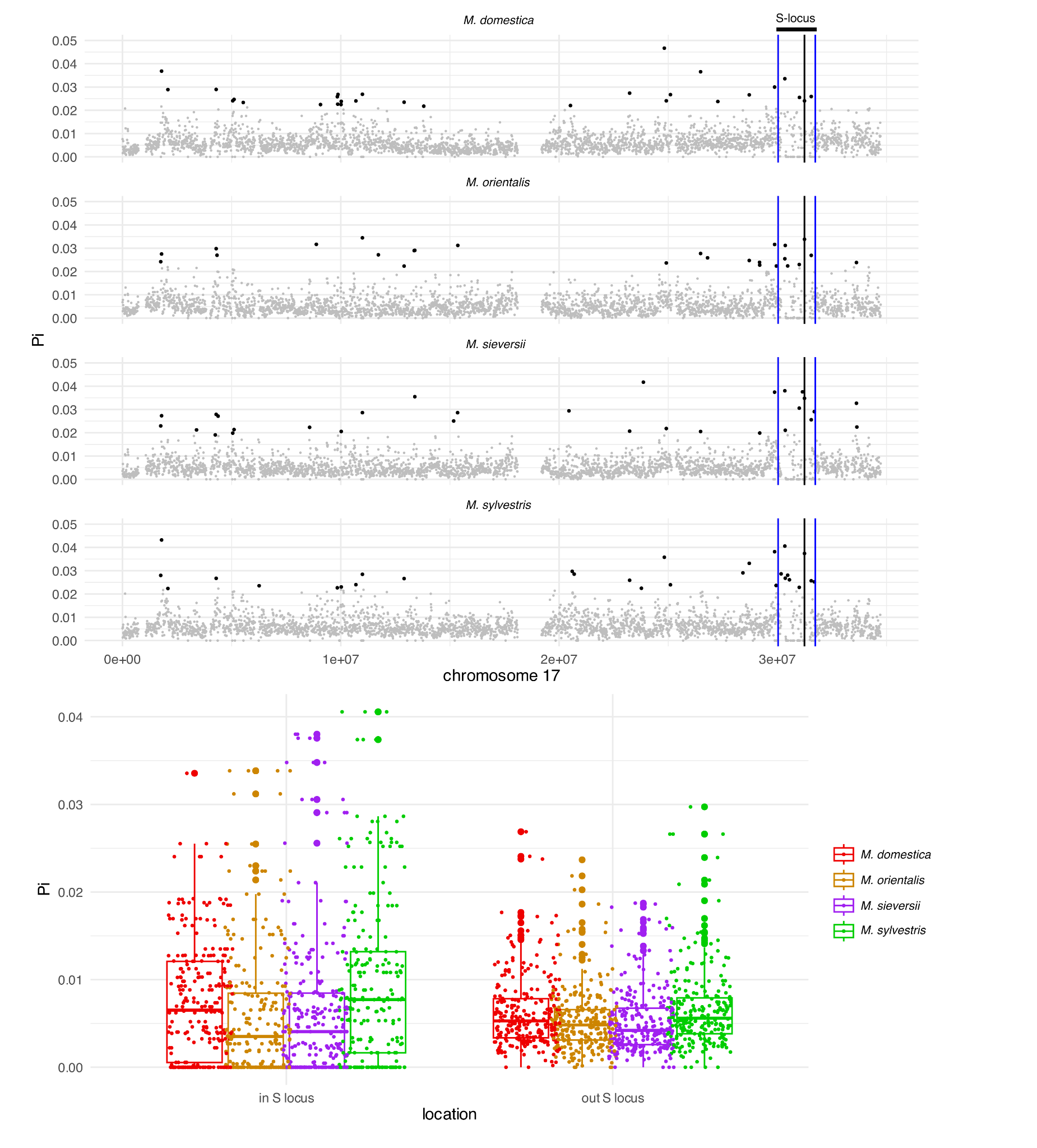

**Supplementary Fig. 10: Nucleotide diversity (π) on chromosome 17 across four *Malus* species.** (Top) Genome-wide distribution of nucleotide diversity on chromosome 17 in *M. domestica*, *M. orientalis*, *M. sieversii,* and *M. sylvestris*, calculated in 10 kb non-overlapping windows using Pixy. Black dots represent the top 1% windows in terms of π, calculated separately for each species. The region corresponding to the S-locus (as annotated in GDDH13; Daccord et al., 2017; haplotype h1 in this study) is highlighted in blue. In all four species, the number of top 1% windows falling within the S-locus was significantly higher than expected by chance based on 10,000 random permutations (*M. domestica*: p-value = 1.07e-2, *M. orientalis*: 3e-4, *M. sieversii*: 1e-4, *M. sylvestris*: 1e-4). (Bottom) Boxplots of nucleotide diversity for 1,000 randomly sampled windows located inside vs. outside the S-locus, for each species. Each dot represents a 10 kb window. Colors correspond to species.

### **Supplementary Table 10: Detection of positive selection on SFBB genes in each S-haplotype using site models implemented in PAML / CodeML.** Results of site models (M0, M1a, M2a, M3, M7, M8) fitted to codon alignments of SFBB genes from haplotypes h1 to h32. For each haplotype, the number of SFBB sequences included (**N**) and the total alignment length in codons (**S**) are indicated. **M0** assumes a single ω ratio for all sites and branches.**M1a** (neutral) and **M2a** (selection) compare models with and without a class of positively selected sites (ω > 1). **M3** allows discrete site classes with different ω values. **M7** (beta distribution) and **M8** (beta + ω > 1) are alternative site models for detecting selection. Model parameters shown include the log-likelihood (lnL), site class proportions (p₀, p₁, p₂), and dN/dS ratios (ω, ω₀, ω₁, ω₂) (Excel file).

### **Supplementary Table 11: Likelihood ratio tests (LRTs) for detection of positive selection on SFBB genes in individual S-haplotypes.** Results of model comparisons conducted using codeml (PAML) on codon alignments of SFBB genes from haplotypes h1 to h32. For each haplotype, three likelihood ratio tests were performed to evaluate the presence of **codons under positive selection**: **M0 vs M3**: compares a one-ratio model (M0) to a discrete model allowing for multiple site classes (M3), testing for variable selection pressure across sites. **M1a vs M2a**: tests for the presence of positively selected codons (ω > 1) by comparing a neutral model (M1a) to a selection model (M2a). **M7 vs M8**: compares a beta-distributed ω model (M7, without selection) to a model allowing an extra site class with ω > 1 (M8). For each comparison, the table reports log-likelihoods (lnL) of both models, the test statistic -2Δℓ, degrees of freedom (**K**), significance level (**alpha**), associated *p*-value, and the chi-squared critical value for 5% significance (Χ²(α)) (Excel file).

### **Supplementary Table 12: Amino acids observed at positively selected sites across SFBB families in 27 S-haplotypes.** For each of the 27 S-haplotypes, the table lists the amino acid residues found at the codon positions most frequently inferred to be under positive selection (posterior probability > 0.95 in PAML analyses). Each cell contains the amino acids present in the given haplotype for a given SFBB family; “absent” indicates that the SFBB gene was not detected in that haplotype (Excel file).

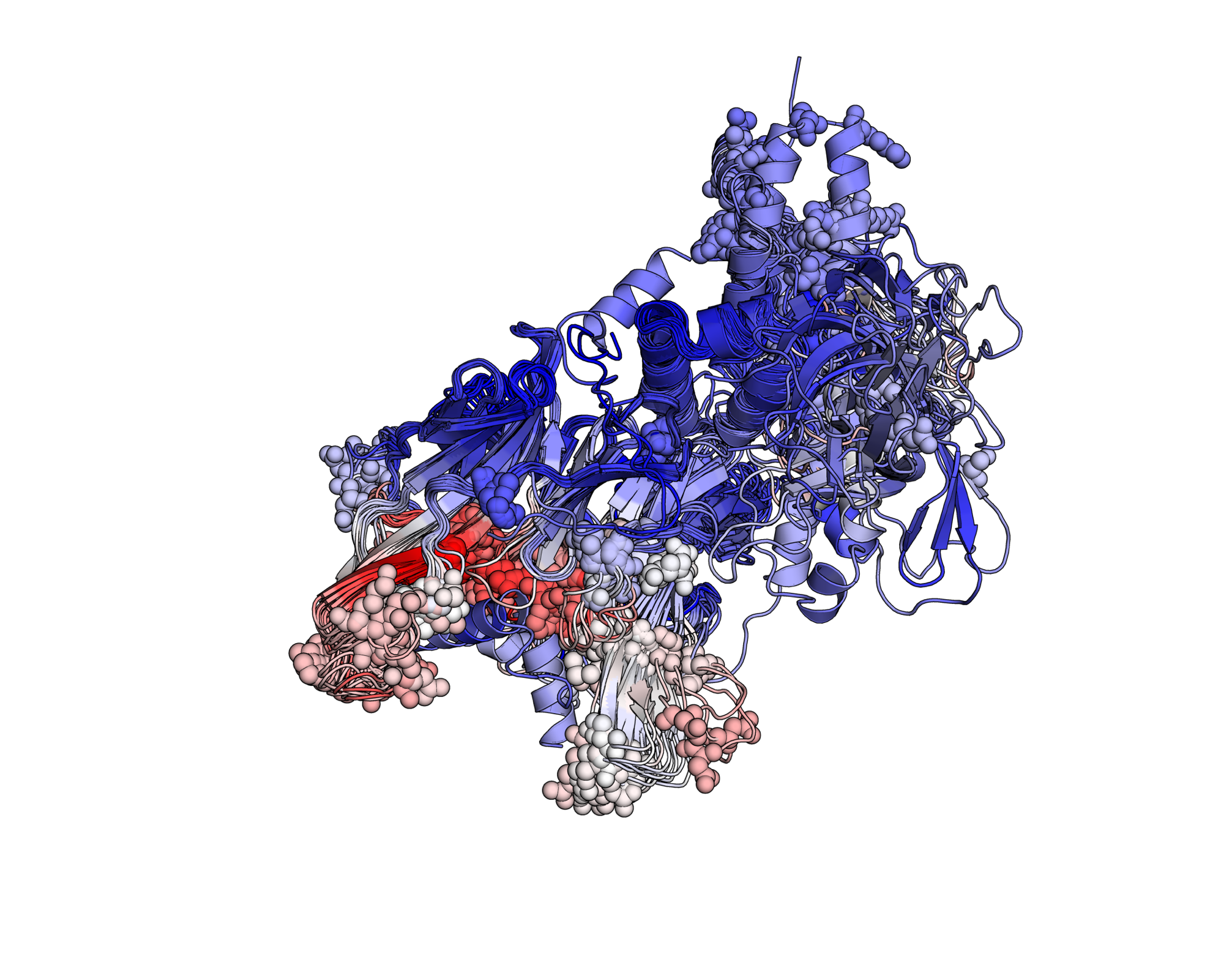

**Supplementary Fig. 11:** **Structural representation of *Malus* SFBBs highlighting positively selected sites.** Ribbon diagram of SFBB structures predicted by Boltz-1, showing the typical α/β fold. The longest sequence for each S-allele was chosen as representative, totalling 17 sequences, and the resulting structures were aligned in PyMOL to minimize pairwise root mean square deviation of atomic positions. The spheres indicate amino acid residues inferred to be under positive selection across SFBB families (posterior probability > 0.95, Bayes Empirical Bayes analysis, CodeML). These selected sites are broadly distributed on the surface of the protein, suggesting their potential involvement in allele-specific recognition or interaction with pistil determinants. Structural elements are colored according to MuLAN attention scores. Red-to-white colors correspond to high while blue tones to low scores.

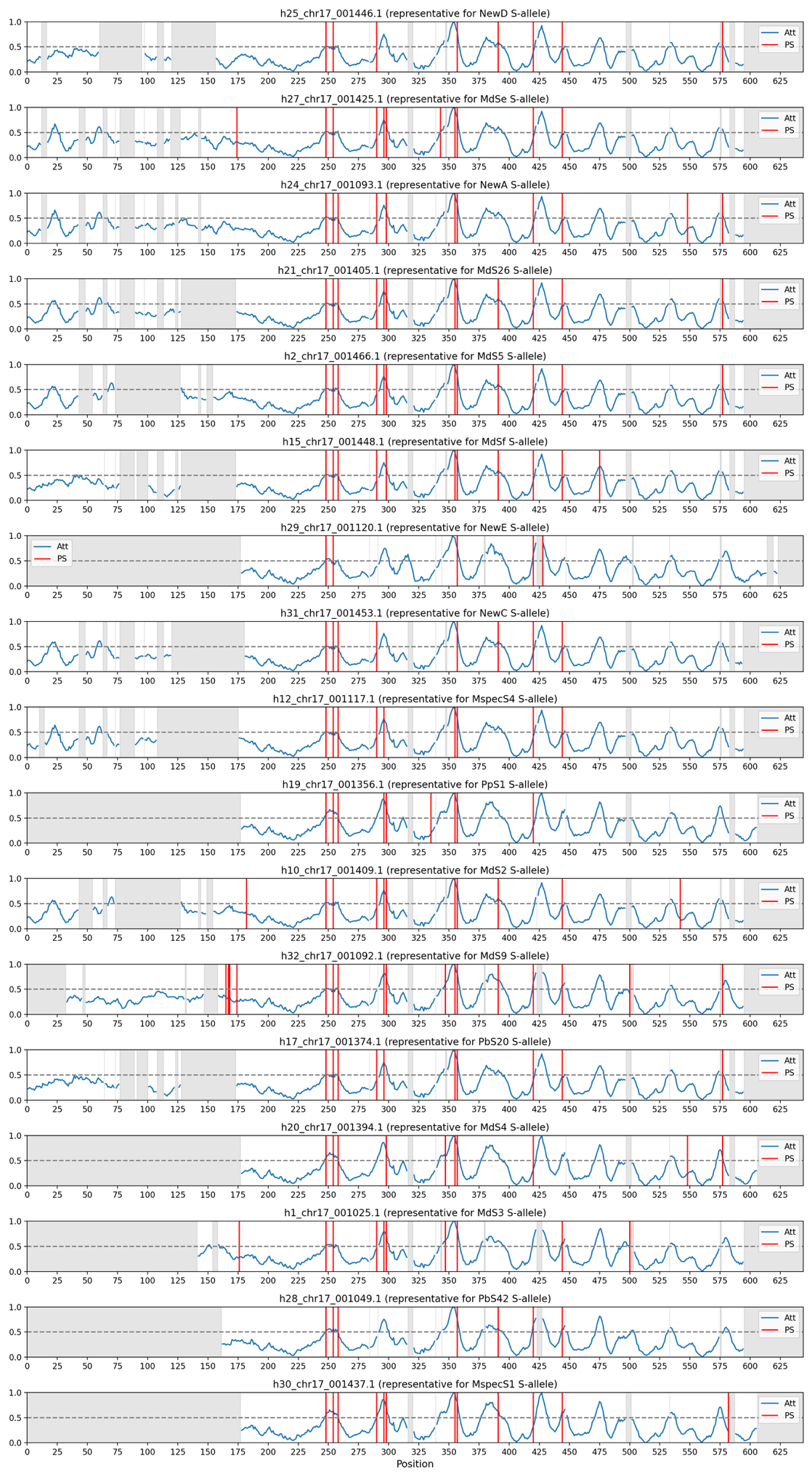

**Supplementary Fig. 12: Alignment of MuLAN attention scores and positively selected sites across *Malus* SFBB proteins.** For SFBB genes associated with a distinct S-allele, we plotted MuLAN attention scores (blue) along the amino acid sequence to estimate the predicted functional relevance of each residue. Vertical red lines indicate sites inferred to be under positive selection. Several of these sites co-localize with peaks in attention scores, suggesting that positively selected residues tend to coincide with positions predicted to have high functional impact. Gray-shaded regions represent alignment gaps or poorly aligned regions excluded from the analysis.

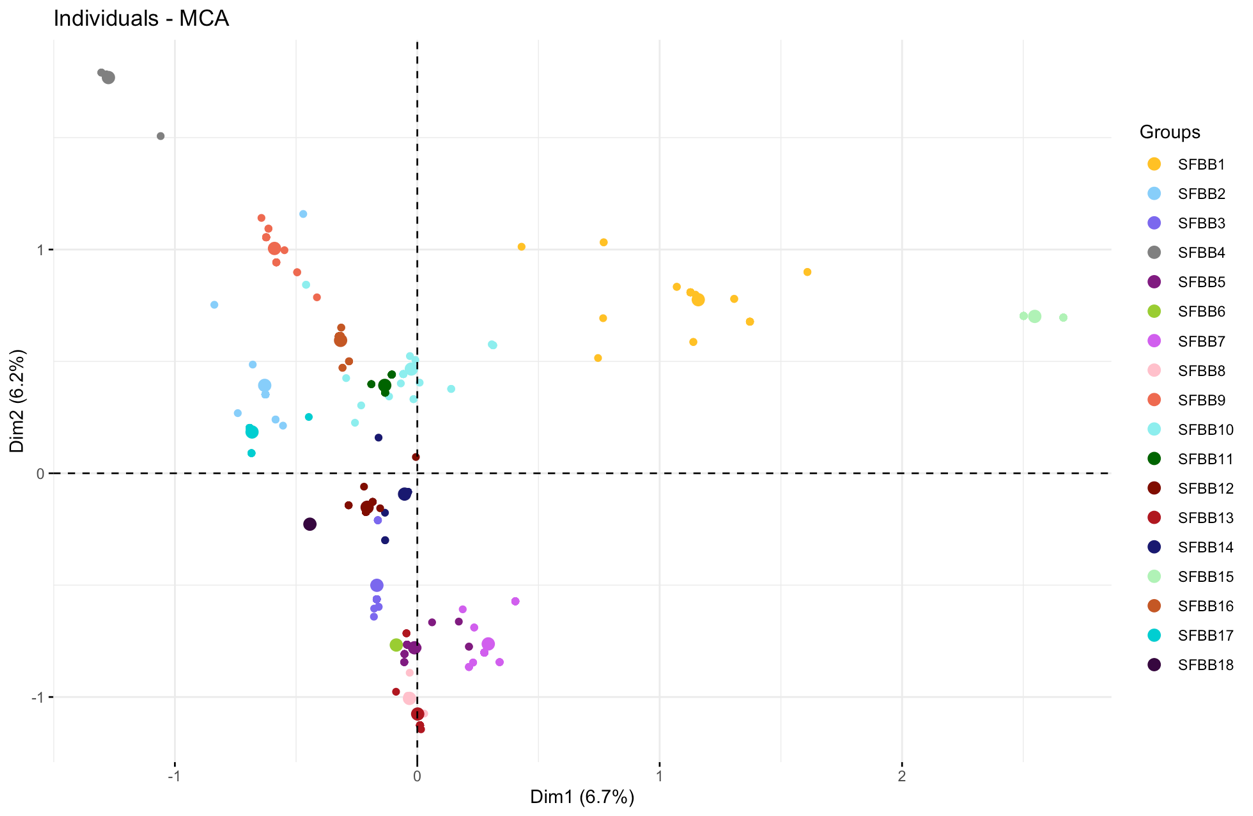

**Supplementary Fig. 13: Clustering of SFBB genes based on amino acid identity at conserved positions**. Multiple correspondence analysis (MCA) was performed using amino acid residues at selected conserved positions across SFBB genes. Each point represents a gene, colored according to its SFBB family. The first two dimensions, which together explain XX% of the total variance (Dim1: 6.7%, Dim2: 6.2%), reveal a structured separation of gene families. Families with similar amino acid profiles tend to cluster together, while others form well-defined and distinct groups in the multivariate space. Ellipses represent 95% confidence intervals for each family.

**Supplementary Table 13: Amino acid composition at conserved codon positions across SFBB families.** Matrix summarizing amino acid (AA) frequencies at ten codon positions previously identified as highly associated with SFBB family identity (see Table 2). Each row corresponds to a codon position, and each column represents one SFBB family. Within each cell, the observed amino acids and their relative frequencies (≥3%) are listed, based on all SFBB gene sequences belonging to the corresponding family. Gray cells indicate positions that are invariant within a family (i.e., a single AA observed at 100% frequency). Rows on the right indicate the number and proportion of SFBB families for which each position is invariant, highlighting positions with family-specific conservation. Similarly, the bottom row shows the number and proportion of invariant positions per SFBB family. These results support the presence of family-specific amino acid signatures likely contributing to functional diversification among SFBB proteins.

| **SFBB family** | **SFBB1** | **SFBB2** | **SFBB3** | **SFBB4** | **SFBB5** | **SFBB6** | **SFBB7** | **SFBB8** | **SFBB9** | **SFBB10** | **SFBB 11** | **SFBB 12** | **SFBB 13** | **SFBB 14** | **SFBB 15** | **SFBB 16** | **SFBB 17** | **SFBB 18** | **Number of invariant family** | **Percentage** |
| --- | --- | --- | --- | --- | --- | --- | --- | --- | --- | --- | --- | --- | --- | --- | --- | --- | --- | --- | --- | --- |
| **position 297** | F:1 | H:0.82, L:0.14, N:0.05 | P:0.74, H:0.26 | P:1 | N:1 | N:1 | N:1 | E:0.96, G:0.04 | F:1 | F:1 | P:1 | H:0.97, Q:0.03 | E:0.89, D:0.11 | P:1 | F:1 | F:1 | Y:1 | P:1 | 13 | 72% |
| **position 299** | Q:0.61, E:0.33, H:0.06 | S:0.95, W:0.05 | Q:0.81, H:0.19 | V:1 | E:1 | Q:1 | E:0.9, D:0.1 | E:1 | L:1 | L:0.93, M:0.03, Q:0.03 | Q:1 | L:0.84, I:0.16 | E:1 | Q:1 | E:1 | Q:1 | L:1 | L:1 | 12 | 67% |
| **position 336** | H:0.48, P:0.42, C:0.03, D:0.03, R:0.03 | D:0.86, N:0.09, S:0.05 | N:0.81, D:0.19 | D:1 | D:0.92, N:0.08 | D:1 | N:0.67, D:0.23, Y:0.1 | D:1 | D:0.65, N:0.26, V:0.09 | D:0.63, H:0.17, A:0.07, N:0.07, V:0.07 | D:0.5, A:0.27, S:0.18, T:0.05 | D:1 | N:0.85, D:0.11, S:0.04 | D:1 | H:1 | D:0.9, N:0.1 | S:1 | D:1 | 8 | 44% |
| **position 356** | I:0.97, V:0.03 | T:0.91, N:0.05, S:0.05 | T:1 | S:1 | V:0.8, I:0.12, T:0.08 | L:1 | T:0.82, I:0.18 | T:1 | S:0.83, T:0.13, N:0.04 | T:1 | I:1 | D:1 | T:1 | I:1 | I:1 | T:1 | V:0.96, I:0.04 | T:1 | 12 | 67% |
| **position 358** | G:0.85, A:0.06, I:0.06, V:0.03 | Q:1 | H:1 | G:1 | K:1 | K:1 | K:1 | K:1 | Q:1 | Q:0.97, L:0.03 | E:1 | G:1 | K:1 | G:0.9, R:0.1 | T:1 | E:1 | R:1 | R:1 | 15 | 83% |
| **position 421** | E:0.82, D:0.06, G:0.06, K:0.03, R:0.03 | D:0.95, Y:0.05 | S:0.81, T:0.19 | K:1 | D:1 | D:1 | E:1 | K:0.96, I:0.04 | T:0.96, R:0.04 | Q:0.7, D:0.1, E:0.07, N:0.07, K:0.03, T:0.03 | K:1 | K:1 | K:1 | K:1 | E:1 | T:0.79, N:0.14, K:0.07 | K:1 | K:1 | 11 | 61% |
| **position 429** | H:0.88, D:0.06, Y:0.06 | N:0.91, H:0.05, S:0.05 | P:1 | L:1 | P:1 | P:1 | P:1 | P:1 | S:1 | H:0.83, S:0.13, T:0.03 | Q:1 | P:1 | P:1 | P:1 | C:1 | S:1 | S:0.96, T:0.04 | P:1 | 14 | 78% |
| **position 476** | E:0.88, K:0.12 | L:0.95, T:0.05 | N:0.81, K:0.19 | Q:1 | K:0.96, E:0.04 | K:1 | K:0.95, N:0.05 | K:1 | R:1 | T:0.97, K:0.03 | K:1 | E:1 | K:1 | K:0.97, R:0.03 | K:1 | T:1 | K:1 | K:1 | 11 | 61% |
| **position 543** | T:0.85, S:0.15 | S:1 | C:0.94, F:0.03, Y:0.03 | S:1 | C:0.64, Y:0.2, F:0.16 | C:1 | C:0.97, S:0.03 | R:0.74, H:0.19, C:0.07 | S:0.78, N:0.17, D:0.04 | C:0.9, S:0.07, Y:0.03 | S:1 | C:0.9, G:0.1 | H:0.85, C:0.11, Q:0.04 | C:0.93, H:0.07 | T:0.72, I:0.28 | S:0.93, N:0.07 | S:1 | G:1 | 6 | 33% |
| **position 549** | V:0.91, F:0.06, A:0.03 | V:0.95, I:0.05 | V:1 | A:0.8, T:0.15, V:0.05 | L:1 | L:1 | L:1 | I:1 | L:1 | L:0.9, M:0.07, V:0.03 | V:1 | I:0.77, L:0.19, F:0.03 | I:0.89, L:0.11 | L:1 | E:1 | L:1 | L:0.82, I:0.18 | L:1 | 11 | 61% |
| **Number of invariant positions** | 1 | 2 | 4 | 9 | 6 | 10 | 5 | 7 | 6 | 2 | 9 | 6 | 6 | 7 | 9 | 7 | 7 | 10 | 113 | 63% |
| **Percentage** | 10% | 20% | 40% | 90% | 60% | 100% | 50% | 70% | 60% | 20% | 90% | 60% | 60% | 70% | 90% | 70% | 70% | 100% |  |  |

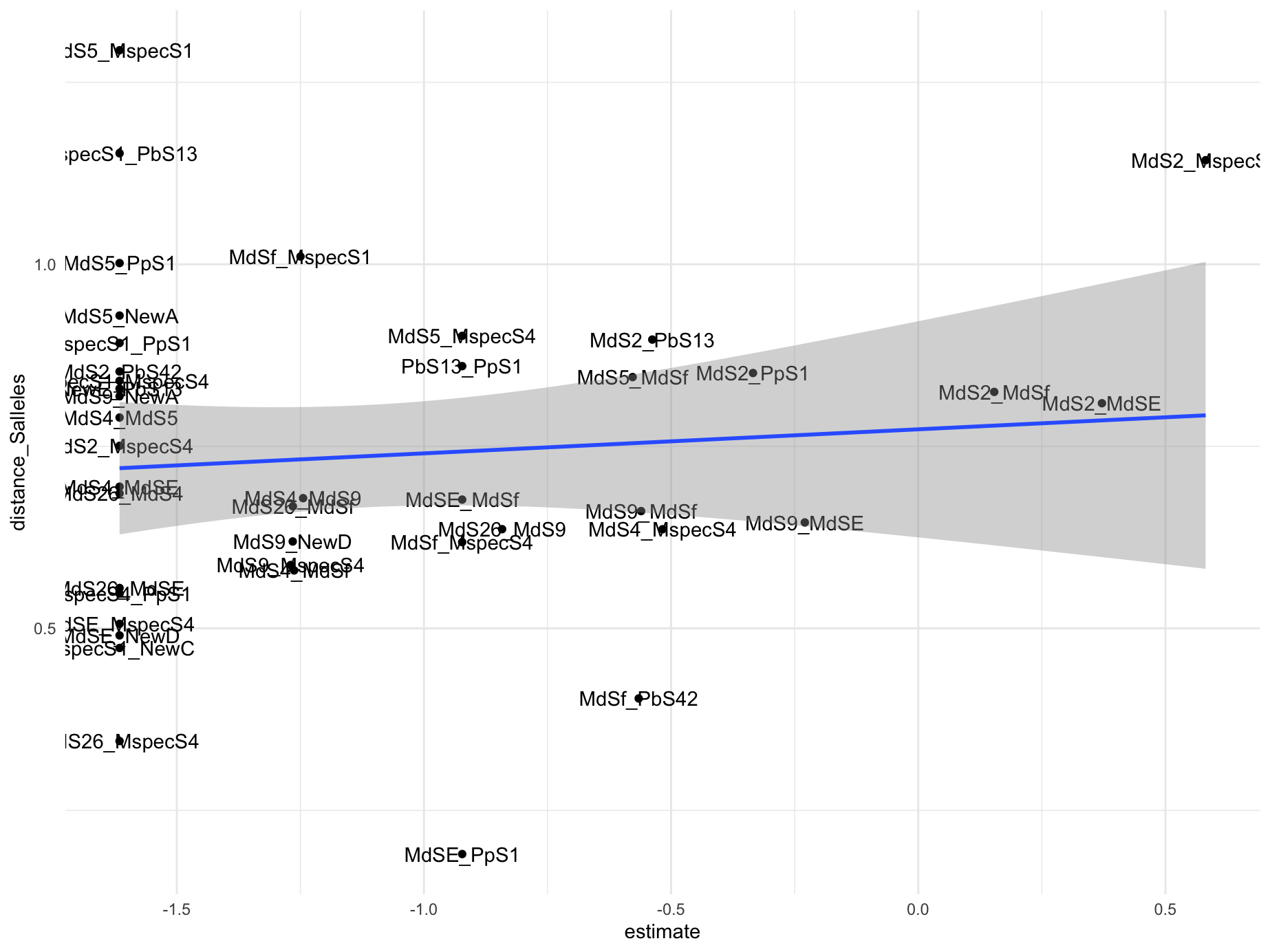

**Supplementary Fig. 14: Relationship between phylogenetic distance of S-RNase alleles and frequency of gene conversion between S-haplotypes.** Each point represents a pair of S-haplotypes, for which gene conversion events were inferred. The x-axis shows the estimate of gene conversion frequency (from the quasi-Poisson model), and the y-axis represents the cophenetic distance between the corresponding S-RNase alleles, extracted from the phylogenetic tree (Supplementary Fig. 1). The blue line shows the linear regression fit with 95% confidence interval (grey area). No significant correlation was detected (Pearson’s r = 0.09, p = 0.57), suggesting that phylogenetic relatedness of S-RNases does not predict gene conversion intensity between their associated SFBB genes.

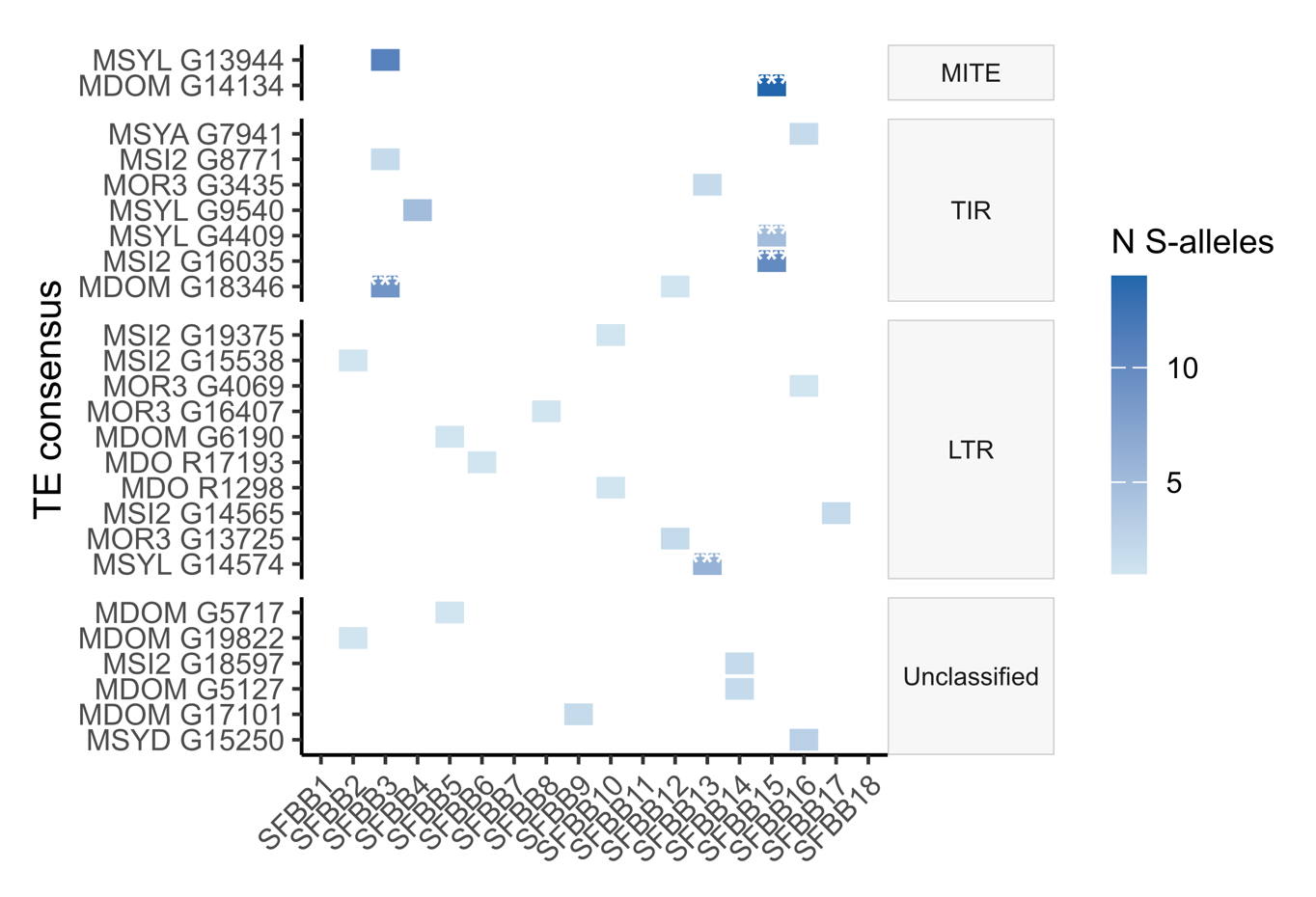

**Supplementary Fig. 15:** **Heatmap of inter-alleleic associations between TE consensus sequences** (rows) **and SFBB gene families** (SFBB1-18, columns), groupes by TE order (MITE, TIR, LTR, Unclassified). Cell sharing indicates the number of S-alleles in which the TE-SFBB association was detected. Asterisks indicate statistically significant associations (* p < 0.05, ** p < 0.01, *** p < 0.001).

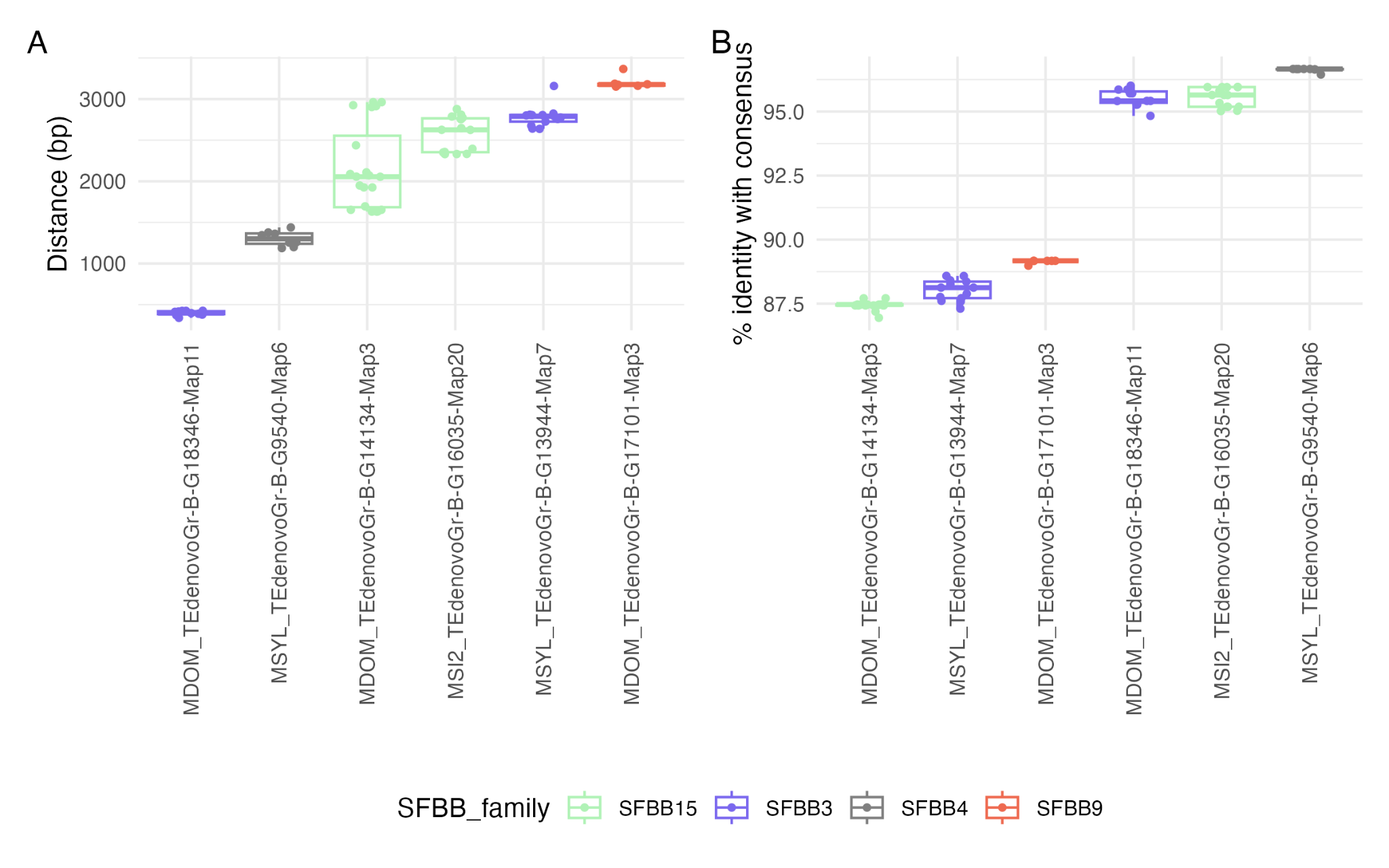

**Supplementary Fig. 16: Distance and sequence identity of TE copies linked to SFBB gene families.** (A) Distribution of distances (in base pairs) between SFBB genes and their associated TE copies, shown separately for each SFBB family. (B) Sequence identity (%) between TE copies associated with SFBB genes and the corresponding TE family consensus sequence.

### **Supplementary Table 14: Contextual information for TE–SFBB associations in the S-locus of *Malus*.** This table reports only TE–SFBB pairs identified as significant in the permutation test. Each row corresponds to a TE copy located within 4 kb of an SFBB gene. For each TE, we report its consensus family, TE order, identity to the consensus, and haplotype of origin, along with the nearest upstream and downstream SFBB families and their respective distances. The relative position of the SFBB gene with respect to the TE (upstream, downstream, overlapping, or inside) is indicated. Haplotype metadata include species of origin and associated S-RNase allele. “Enriched” TE consensus and SFBB families pairs (Excel file).

**Supplementary Table 15: Transposable elements (TEs) detected in association with SFBB genes in the *Malus* S-locus.** Each row represents a TE consensus identified in proximity (< 4kb) to SFBB gene families across the 27 haplotypes analyzed. The columns indicate the class (I = retrotransposon, II = DNA transposon), order (e.g., LTR, TIR, MITE), superfamily (e.g., Ty/Gypsy, hAT, MuDR), and the closest match in RepBase (if any), which provides a putative annotation based on sequence similarity. “unclassified” indicates elements that could not be confidently assigned to a known TE category, and “–” marks missing or uncertain information.

| **TE REPET Consensus ID** | **Consensus size** | **class** | **order** | **superfamily** | **Match on RepBase** |
| --- | --- | --- | --- | --- | --- |
| MSYL_TEdenovoGr-B-G9540-Map6 | 1818 | II | TIR | hAT | HAT-4_Mad |
| MDOM_TEdenovoGr-B-G18346-Map11 | 691 | II | TIR | hAT | HAT-10_Mad |
| MSI2_TEdenovoGr-B-G16035-Map20 | 649 | II | TIR | MuDR | MuDR-4_Mad |
| MSYL_TEdenovoGr-B-G13944-Map7 | 574 | I | LTR | Ty3/Gypsy | Gypsy-38_Mad |
| MDOM_TEdenovoGr-B-G14134-Map3 | 371 | II | MITE | - | - |
| MDOM_TEdenovoGr-B-G17101-Map3 | 484 | unclassified | unclassified | unclassified | - |

**Supplementary Table 16: Gene conversion events detected between SFBB alleles using GENECONV (Excel file).** Pairwise gene conversion events among SFBB coding sequences were identified with GENECONV. Only significant events (pairwise BC_KA p-value < 0.05) between sequences of the same SFBB family but distinct S-alleles are reported, restricting the dataset to interallelic conversion events. For each event, the table reports the two sequences involved (seqA, seqB), the global (Sim_pvalue) and pairwise (BC_KA_pvalue) significance of the conversion signal, the alignment coordinates and length of the converted fragment (Aligned_begin, Aligned_end, Offset_length), the number of polymorphic sites (nb_poly), observed differences within the fragment (nb_difs) and total pairwise differences outside the fragment (tot_difs), and the SFBB family (SFBBfamA/SFBBfamB), species (speciesA/speciesB) and S-allele (S_alleleA/S_alleleB) identity of each sequence. Combined pair-level identifiers (pair_sp, pair_Sallele, pair_SFBB) were used to summarize events by species, S-allele, and gene family combination in Figure 3D.

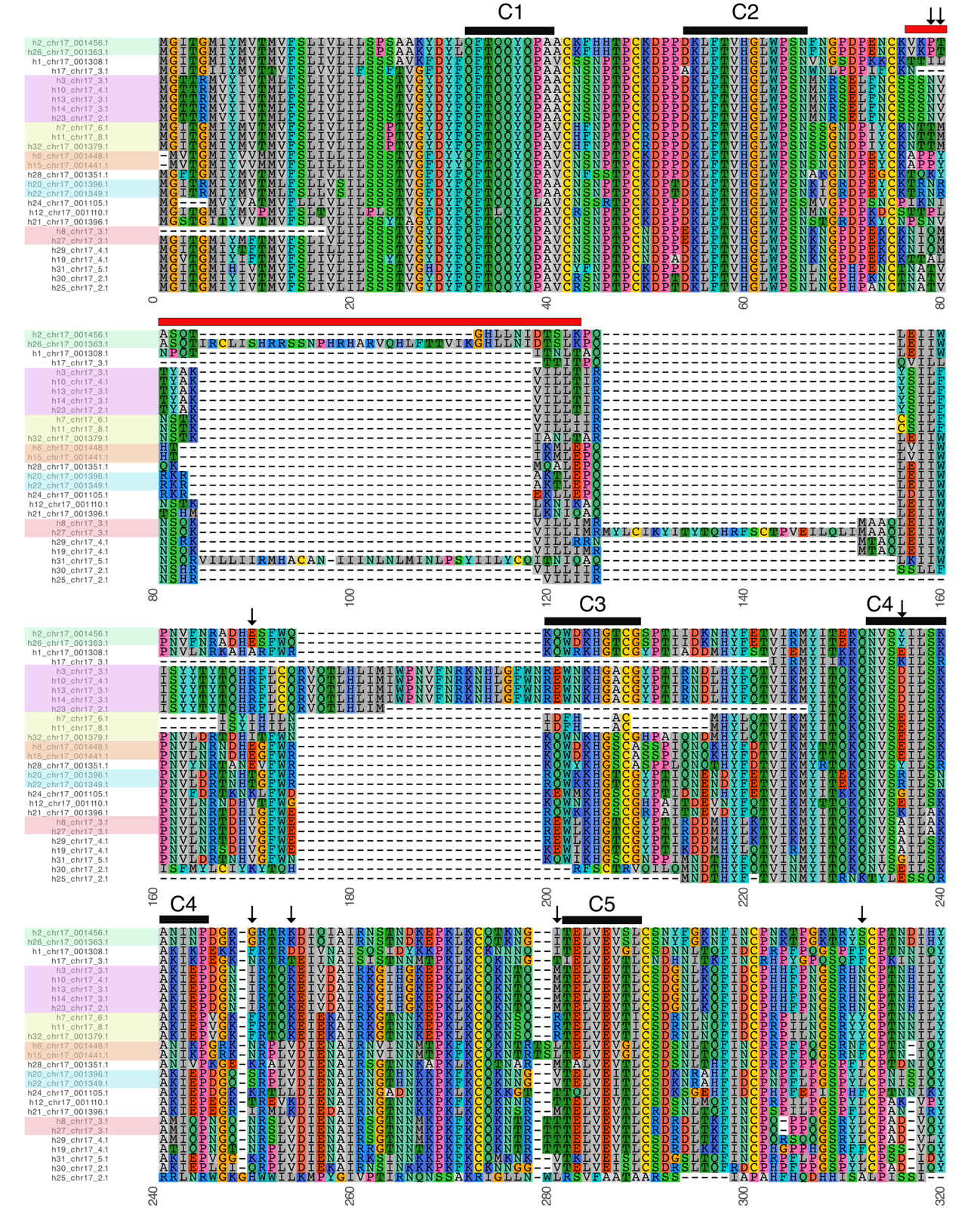

**Supplementary Fig. 17: Multiple *Malus* S-RNase amino-acids sequence alignment.** The background color of sequence names reflects their allele, with identical colors indicating the same S-RNase specificity. Each amino acid is represented by its one-letter code. Background colors indicate amino acid chemical properties, with similar colors assigned to residues of similar physicochemical characteristics (e.g., hydrophobic in grey, aromatic in cyan, acids in red, basics in blue, polar in green, specials in yellow, orange and pink). The alignment is divided into consecutive blocks of 80 amino acids for readability. Positions under diversifying selection are marked with an arrow (↓) above the alignment columns. Gaps introduced during the alignment process are shown in white. The putative hypervariable region (in red) defined by Ishimizu et al. (1998) and Ushijima et al. (1998) and five conserved regions (C1, C2, C3, C4 and C5) are underlined.

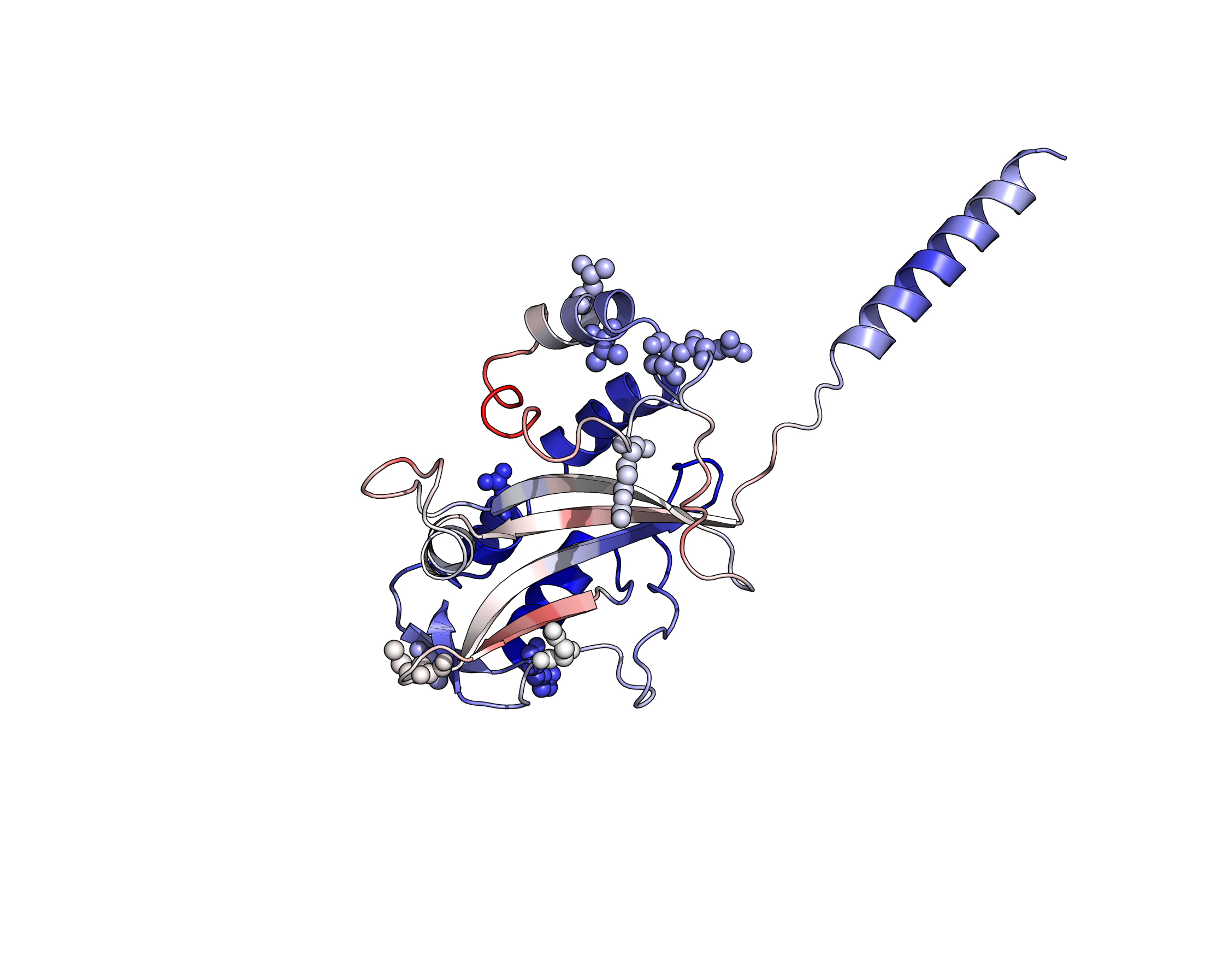

**Supplementary Fig. 18:** **Structural representation of a *Malus* S-RNase highlighting positively selected sites.** Ribbon diagram of a representative S-RNase protein structure (Accession MdS2_h23_chr17_2.1), showing the typical α/β fold. The spheres indicate amino acid residues inferred to be under positive selection across S-haplotypes (posterior probability > 0.95, Bayes Empirical Bayes analysis, CodeML). These selected sites are broadly distributed on the surface of the protein, suggesting their potential involvement in allele-specific recognition or interaction with pollen determinants. Structural elements are colored according to MuLAN attention scores. Red-to-white colors correspond to high while blue tones to low scores.

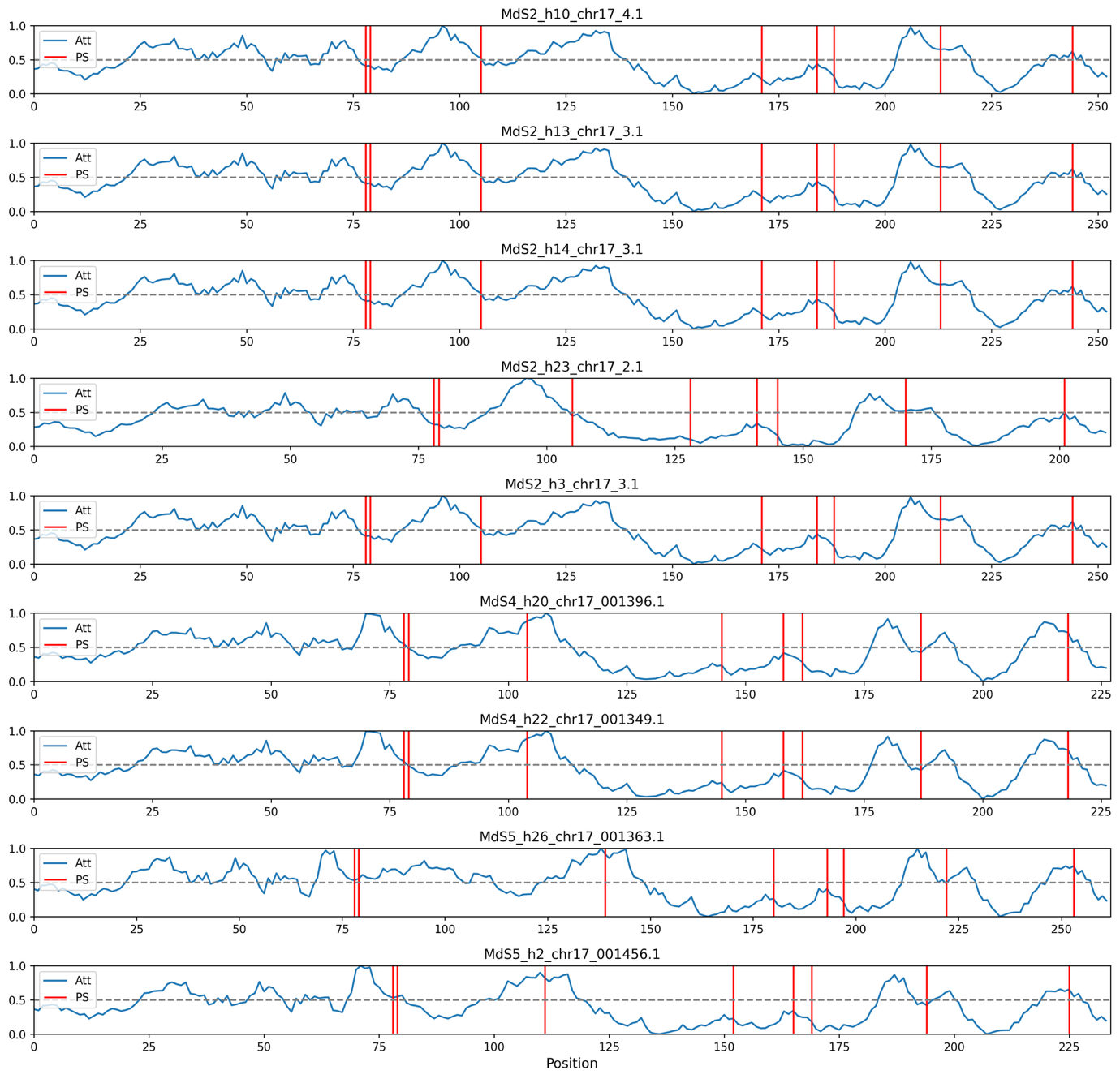
**Supplementary Fig. 19: Comparison of MuLAN attention scores and positively selected sites across *Malus* S-RNase sequences.** MuLAN attention scores (blue) are shown along the amino acid sequences of several S-RNases (MdS2, MdS4, MdS5). Vertical red lines mark positions inferred to be under positive selection (posterior probability > 0.95, CodeML analysis). While some positively selected sites coincide with peaks in attention scores, others appear in regions with low model-derived attention, suggesting that not all adaptively evolving residues are predicted to be functionally critical by the sequence-based model.

**Supplementary Table 17: Putative targeting SFBB families determined for each S-RNases in the *Malus* dataset.** Conclusions are based on the absence of SFBB families or on SFBB proteins mutations in S alleles. Sites located in the FBA domain, involved in the interaction with S-RNase are in bold characters.

| **S allele (S-RNase)** | **Putative targeting SFBB family (supported by)** |
| --- | --- |
| MdS2 (5 haplotypes) | SFBB2 (absence from all haplotypes)  SFBB3 (consistent AA variations on sites 299, **336**, **421** and **476**) |
| MdS4 (2 haplotypes) | SFBB11 (absence from all haplotypes) |
| MdS5 (2 haplotypes) | SFBB11 (absence from all haplotypes) |
| MdS9 (3 haplotypes) | SFBB12 (consistent AA variation on site 543),  SFBB13 (consistent AA variation on sites 297, **336**, 543, 549),  SFBB16 (consistent AA variation on site **336**) |
| MdSf (2 haplotypes) | SFBB5 (consistent AA variation on site **336**),  SFBB7 (consistent AA variation on site **476**) |
| MdSe (2 haplotypes) | SFBB9 (absence from all haplotypes) |

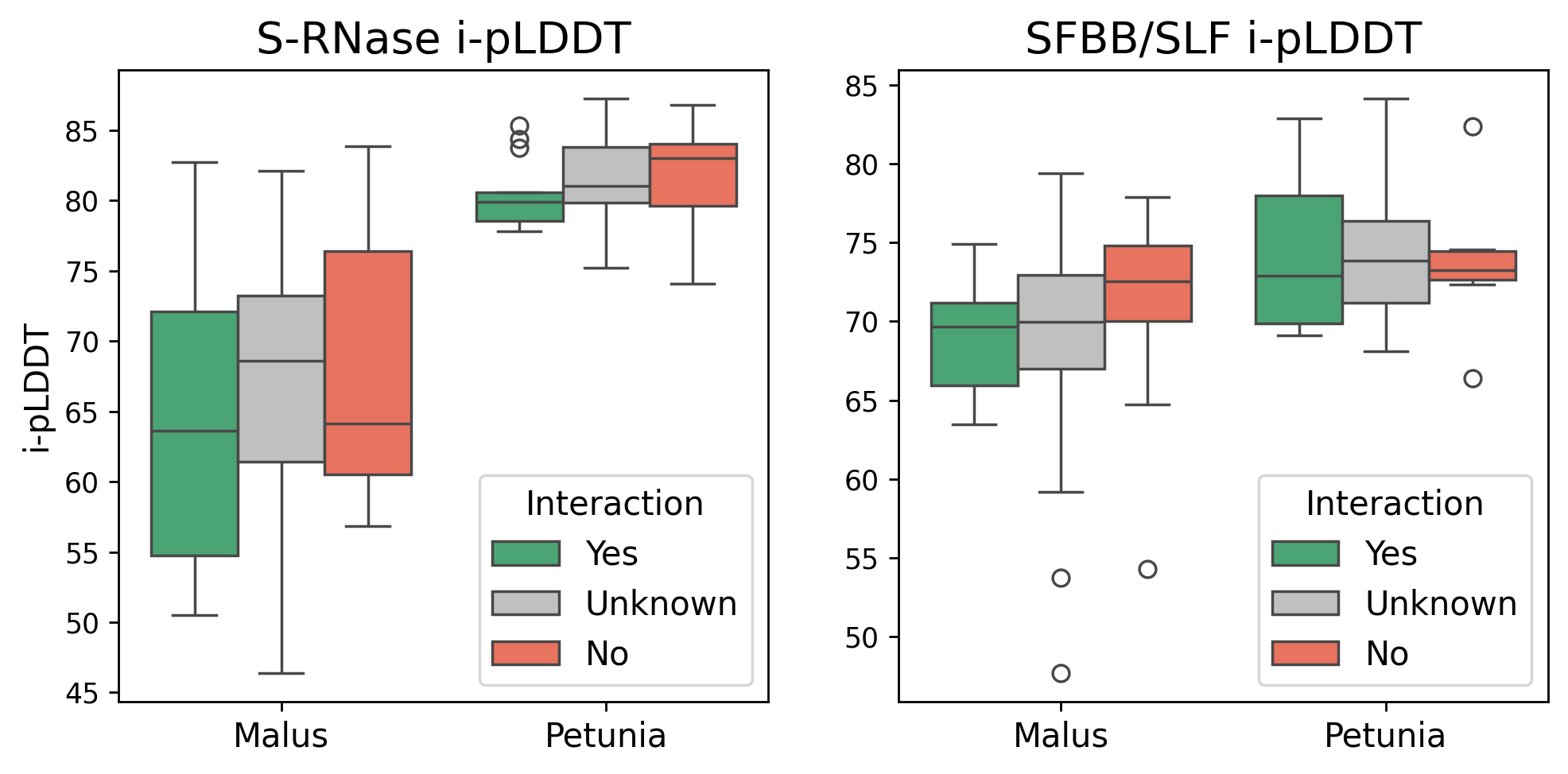

**Supplementary Fig. 20: i-pLDDT confidence scores for individual S-RNase and SFBB/SLF protein models.** i-pLDDT scores for isolated protein chains (i.e., outside the docking context) in *Malus* and *Petunia*. Chain-level models exhibit higher and more consistent confidence than complex-level models, indicating that the folded structures themselves are robustly predicted, whereas the inferred interfaces contribute most of the uncertainty. Although *Petunia* proteins show generally higher confidence values, these do not reliably distinguish known interacting pairs from non-interacting combinations.

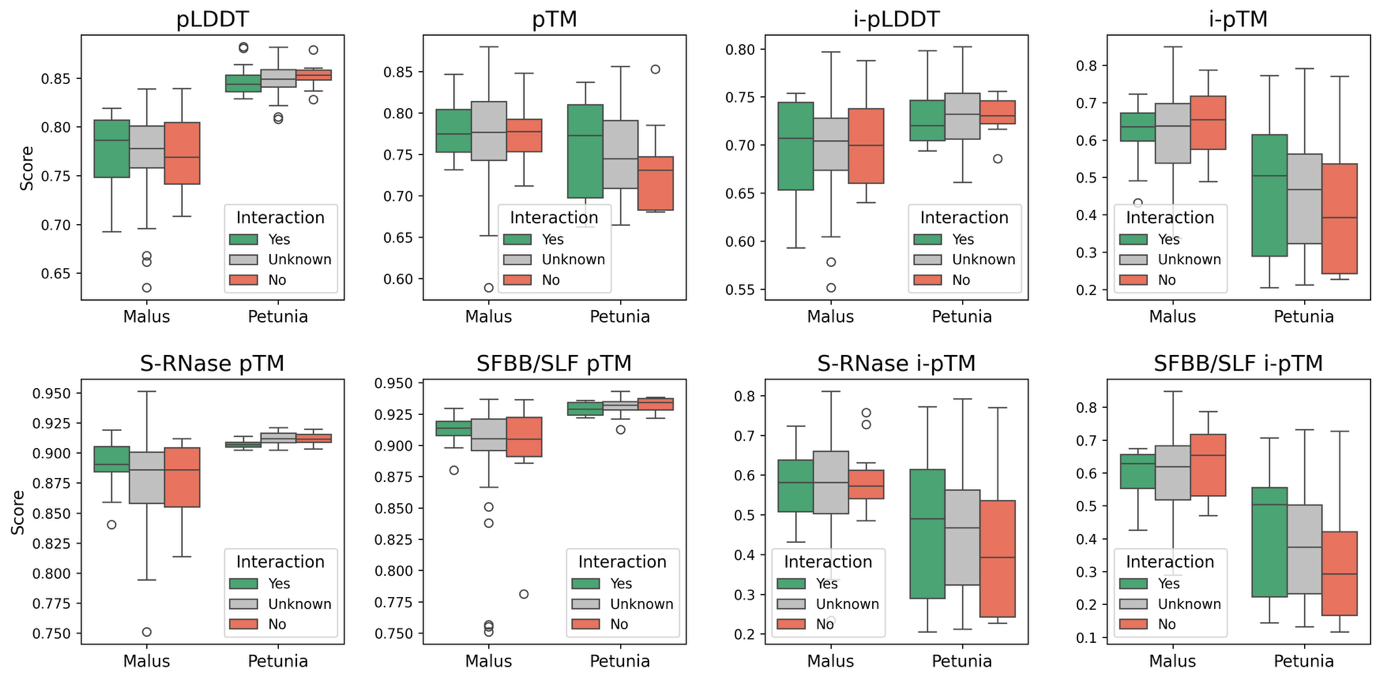

**Supplementary Fig. 21: Structural confidence scores for S-RNase and SFBB/SLF models and predicted complexes in *Malus* and *Petunia*.** Distributions of pLDDT, pTM, i-pLDDT and i-pTM scores for individual proteins and predicted SFBB/SLF–S-RNase complexes. Interaction categories correspond to expected (Yes, in green), unsupported (No, in red), or unknown (Unknown, in grey) functional relationships. In *Malus*, score distributions overlap substantially between categories, indicating that structural confidence metrics alone do not discriminate interacting from non-interacting pairs. In *Petunia*, mean scores are generally higher, but the separation between known and unknown interactions remains incomplete.

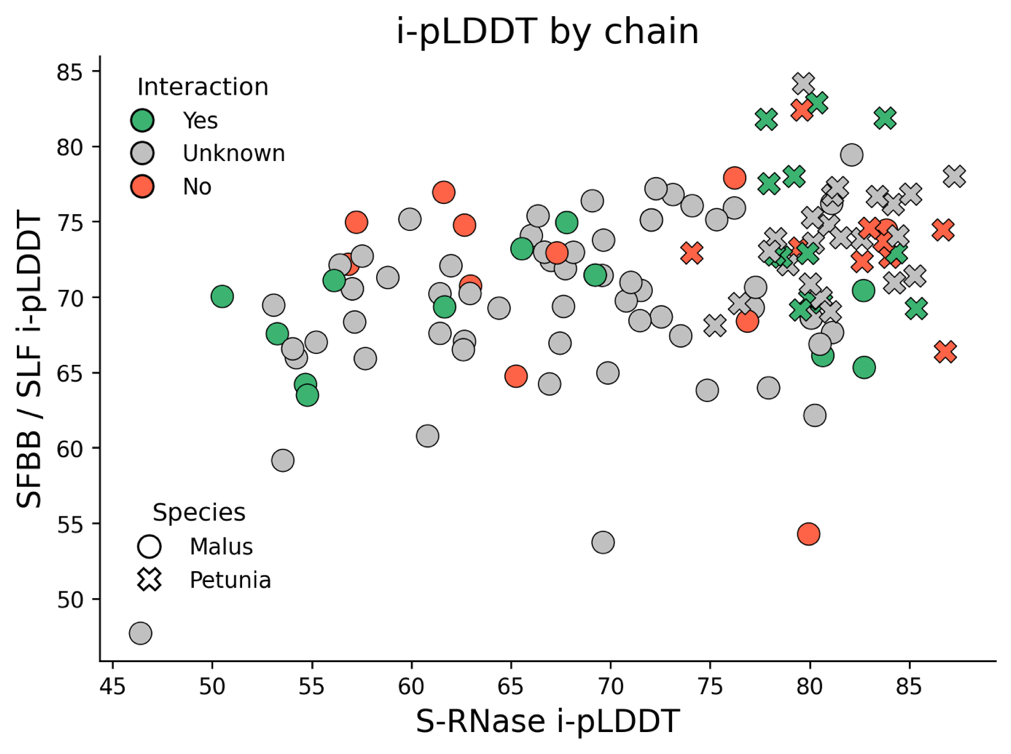

**Supplementary Fig. 22: Relationship between S-RNase and SFBB/SLF interaction confidence across predicted complexes.** Each point represents a predicted S-RNase–SFBB/SLF complex, plotted by the i-pLDDT score of each partner. Colors indicate expected interaction status, and symbol shapes distinguish *Malus* (circles) and *Petunia* (crosses). Although *Petunia* complexes tend to show higher i-pLDDT values overall, the ranges overlap across interaction categories, including several experimentally validated pairs. This indicates limited predictive resolution for interaction specificity based on Boltz-1 docking confidence alone.

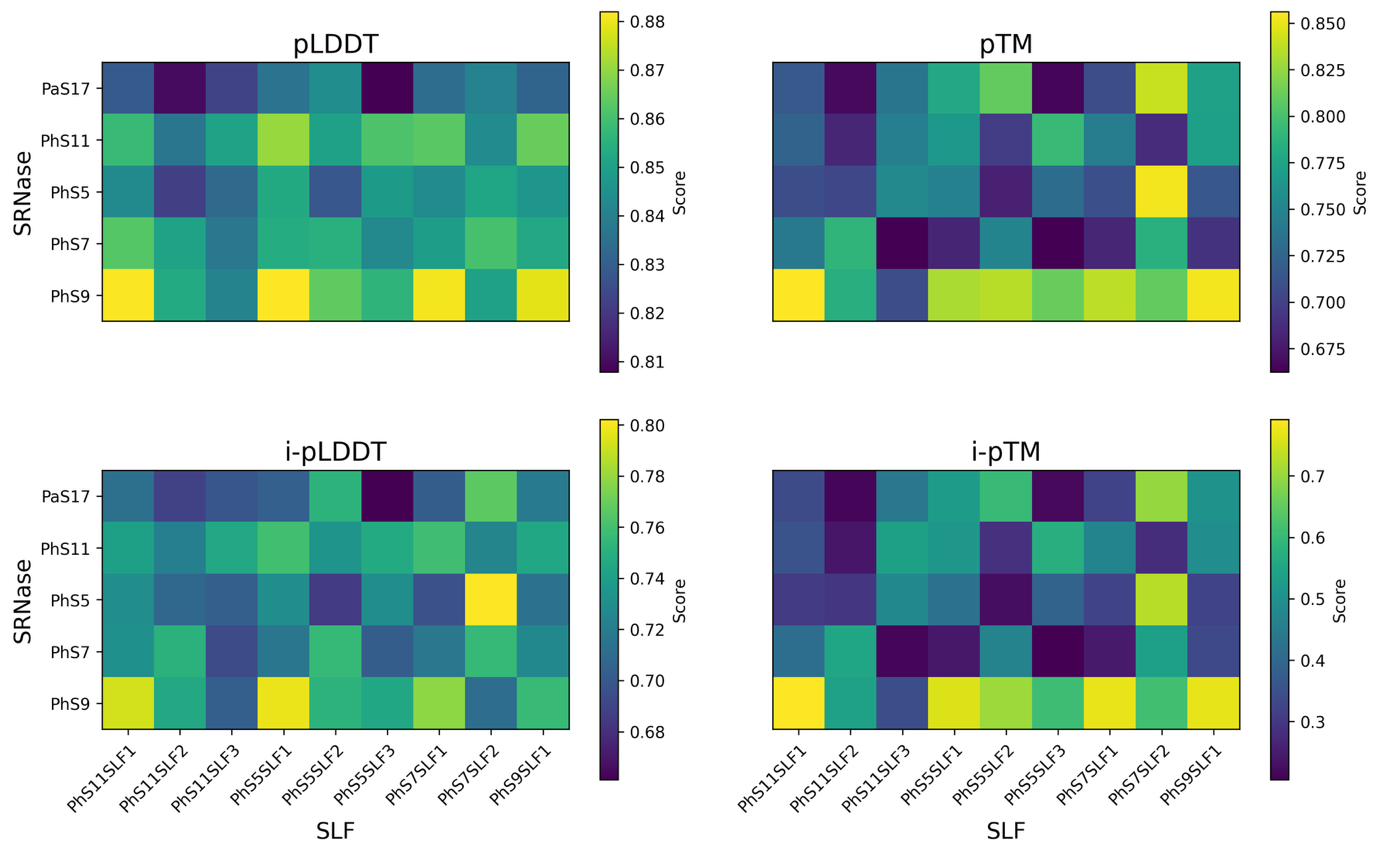

**Supplementary Fig. 23:** **Predicted interaction scores between *Petunia* SLF proteins and S-RNases.** Interaction predictions were computed using the Boltz-1 model for all pairwise combinations of 9 SLF proteins (x-axis) and five S-RNase proteins (y-axis). Four different metrics are shown: pLDDT (predicted local distance difference test, top left), pTM (predicted template modeling score, top right), i-pLDDT (interface pLDDT, bottom left), and i-pTM (interface pTM, bottom right), each representing different confidence measures of the predicted protein–protein interaction. Lighter colors indicate higher predicted interaction scores. SLF and S-RNase identifiers include haplotype and gene id.

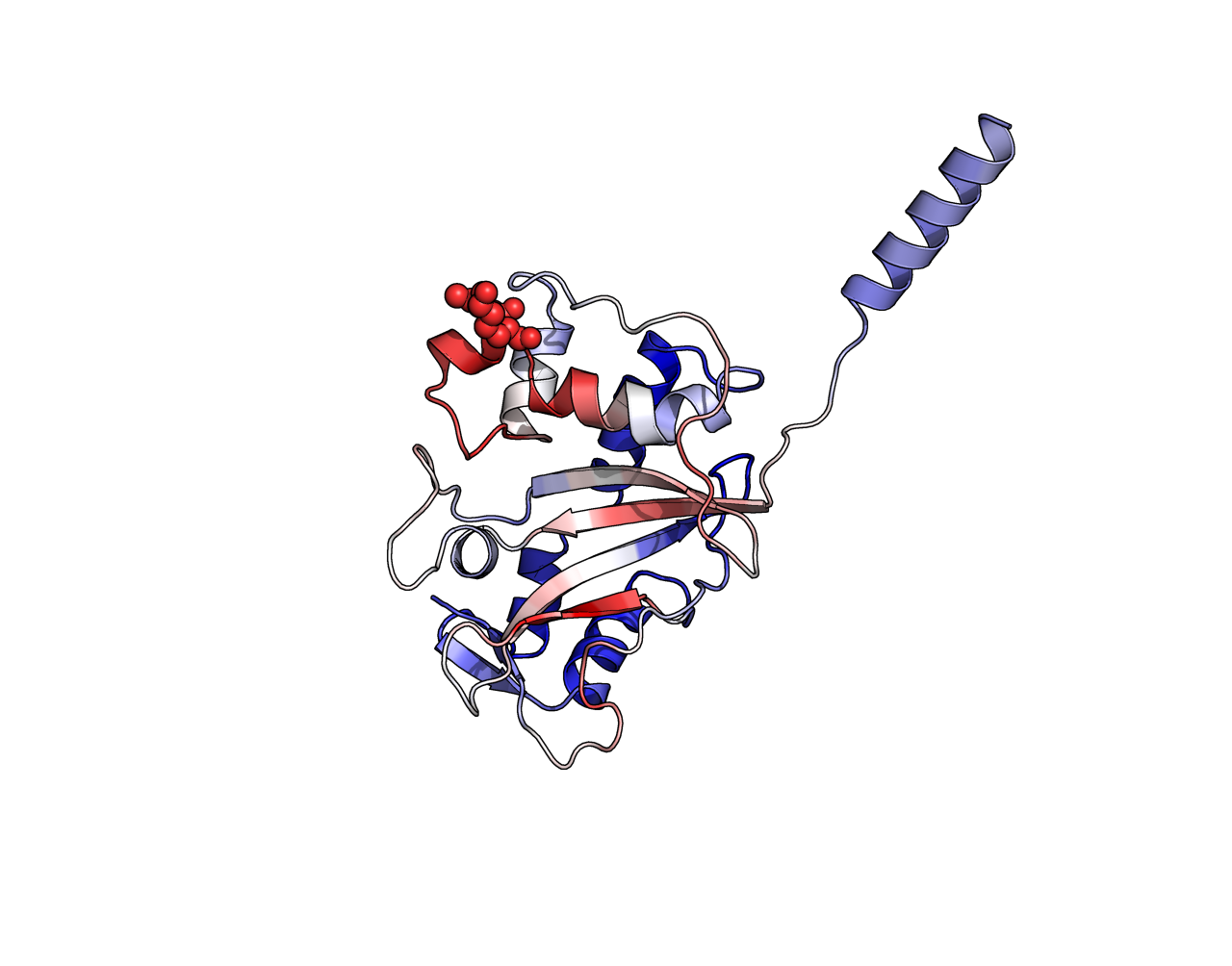

**Supplementary Fig. 24: Structural representation of a *Petunia* S-RNase highlighting positively selected sites.** Ribbon diagram of a representative S-RNase protein structure (Accession LC819199), showing the typical α/β fold. The spheres indicate amino acid residues inferred to be under positive selection across S-haplotypes (posterior probability > 0.95, Bayes Empirical Bayes analysis, CodeML). These selected sites are broadly distributed on the surface of the protein, suggesting their potential involvement in allele-specific recognition or interaction with pollen determinants. Structural elements are colored according to MuLAN attention score. Red-to-white colors correspond to high while blue tones to low scores.

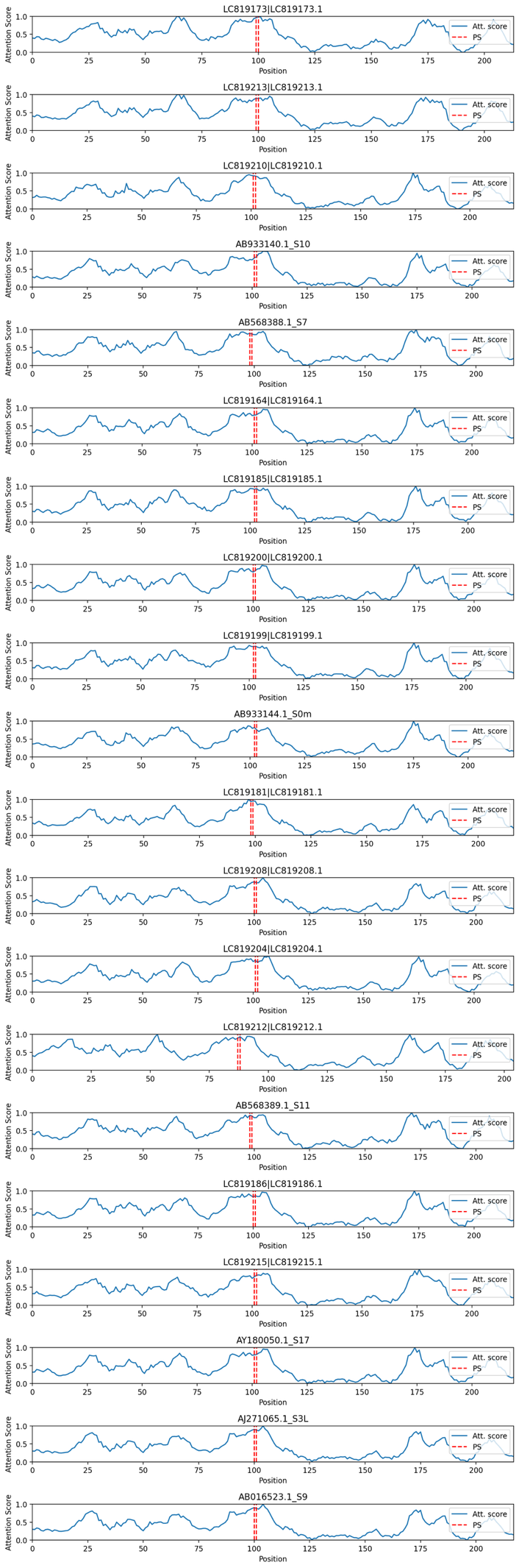

**Supplementary Fig. 25: Comparison of MuLAN attention scores and positively selected sites across *Petunia* S-RNase sequences.** MuLAN attention scores (blue) are shown along the amino acid sequences of several S-RNases. Vertical red lines mark positions inferred to be under positive selection (posterior probability > 0.95, CodeML analysis). Positively selected sites coincide with peaks in attention scores.

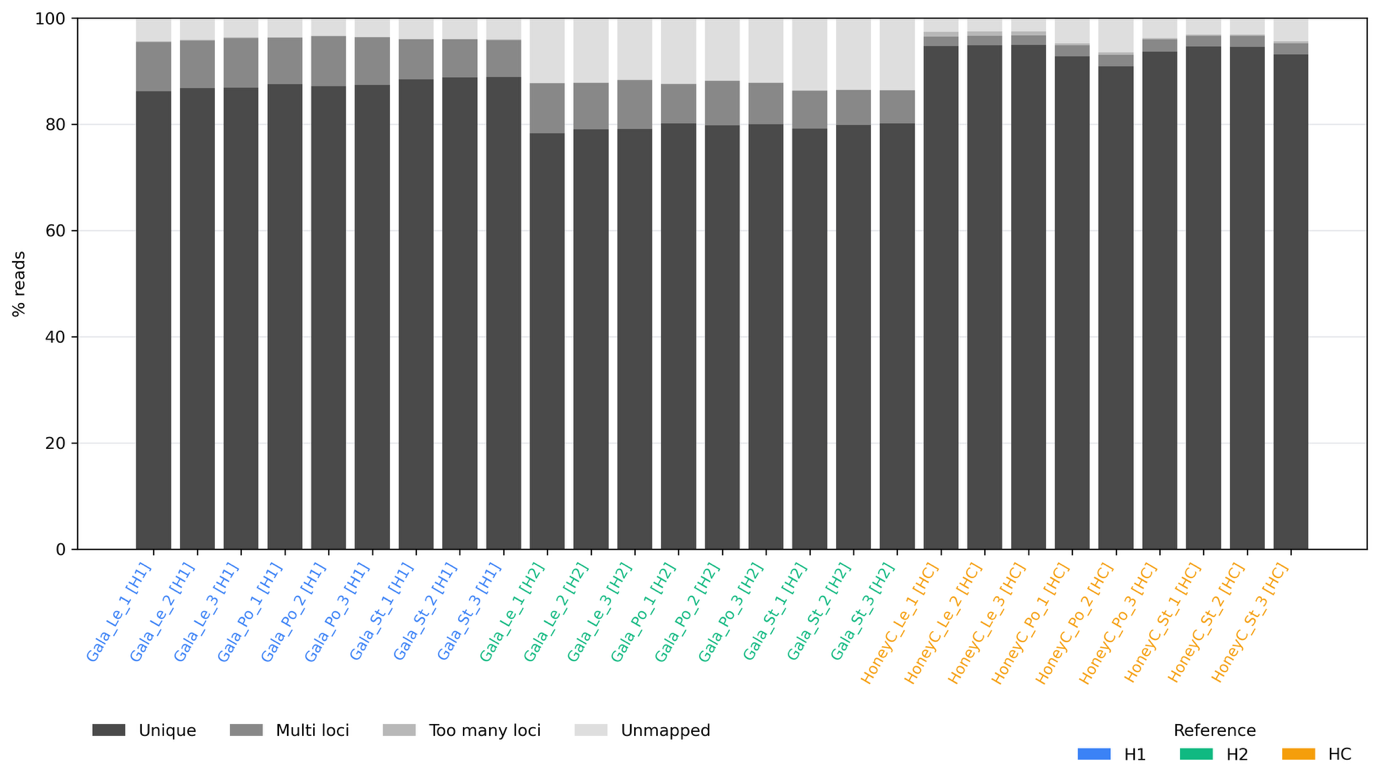

**Supplementary Fig. 26: RNA-seq alignment composition across 27 samples aligned to three *Malus* reference genomes.** Stacked bars show the proportion of uniquely mapped, multi-mapped, too-many-loci, and unmapped reads per sample for nine RNA-seq samples (three biological replicates each of leaf, pollen and pistil) aligned to the Gala haplotype 1 (blue), Gala haplotype 2 (green), and Honeycrisp (orange) reference genomes using STAR. See Supplementary Note 3 for methods.

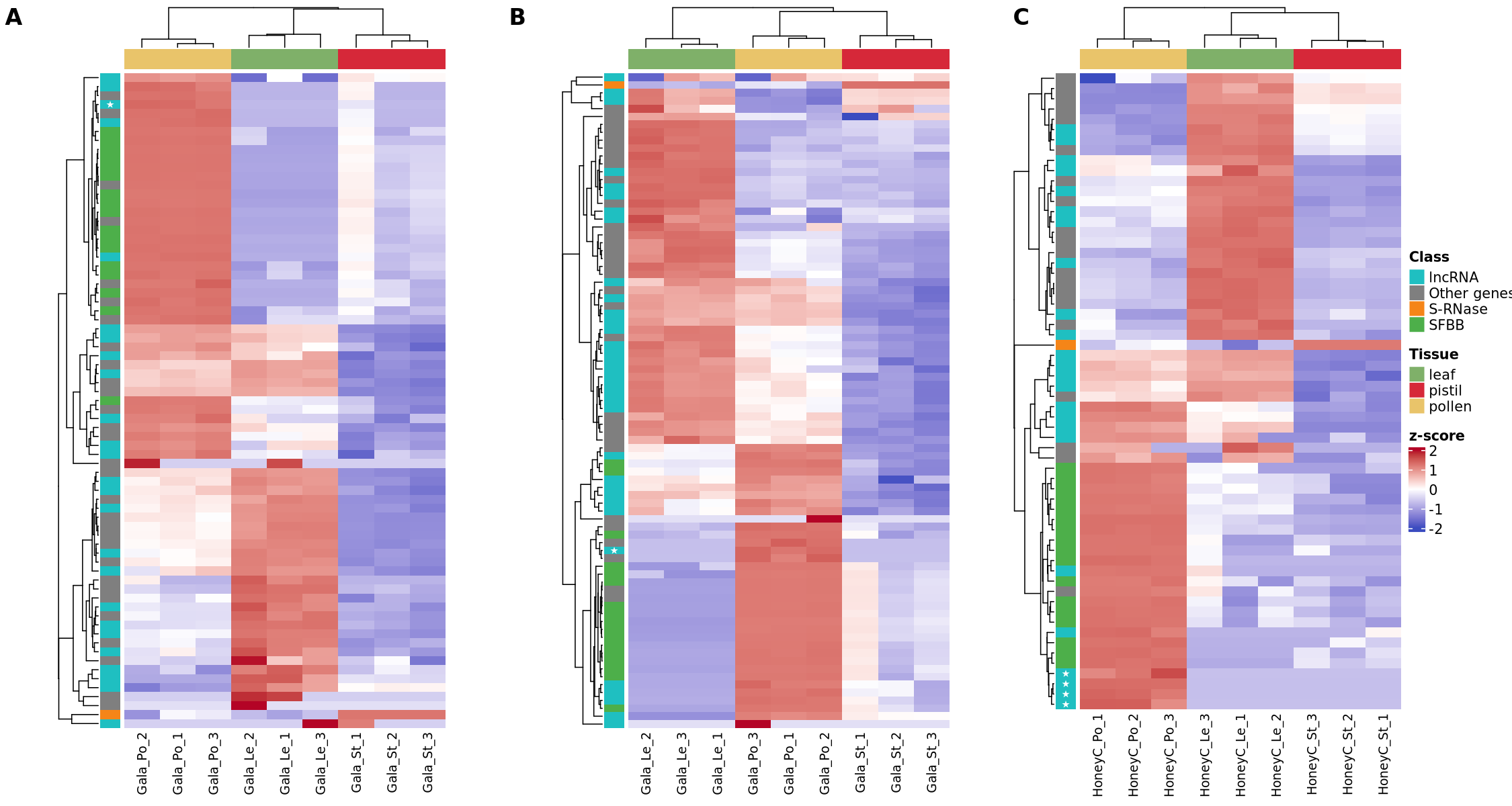

**Supplementary Fig. 27: Expression of S-locus genes across tissues in three apple accessions.** Heatmaps showing the expression of S-locus genes in leaf, pistil and pollen for (A) Gala haplotype 1 (h2), (B) Gala haplotype 2 (h3) and (C) Honeycrisp (h10). Colors represent z-scores of log₂(TPM + 1) values, capped at ±2 (blue: low expression; red: high expression). Only genes with a mean TPM ≥ 1 in at least one tissue are shown. Rows are ordered by hierarchical clustering; columns are grouped by tissue. The left annotation bar indicates gene class: S-RNase (orange), SFBB (green), lncRNA (blue) and other S-locus genes (grey). Stars (★) mark lncRNAs specifically expressed in pollen (mean TPM ≥ 1 in pollen, ≤ 1 in leaf and pistil, in at least two replicates). See Supplementary Note 3 for methods.

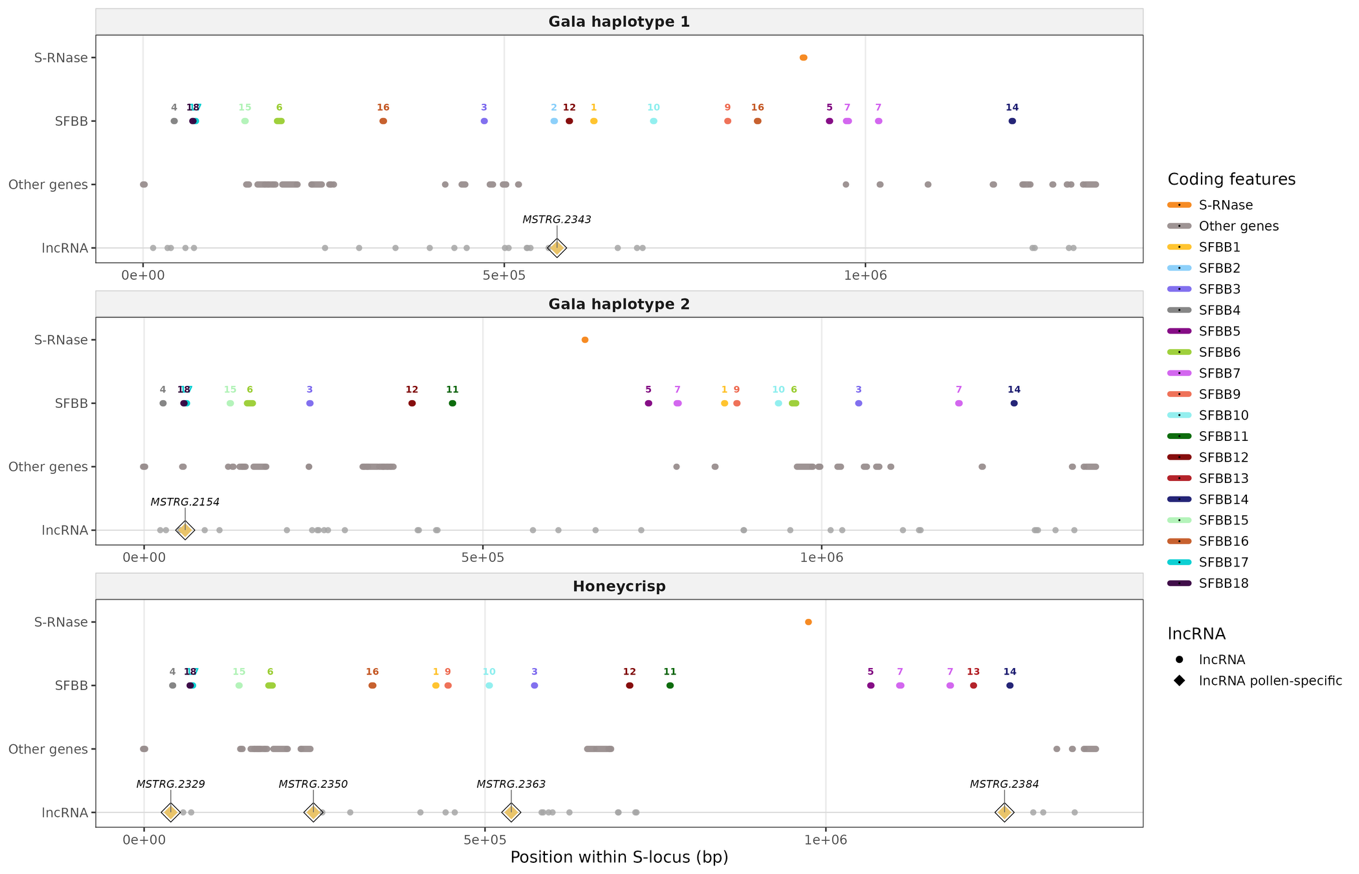

**Supplementary Fig. 28: Distribution of lncRNAs across the S-locus in three apple haplotypes.** Genomic map of the S-locus region (~1.3–1.4 Mb) in Gala haplotype 1, Gala haplotype 2, and Honeycrisp, showing the positions of coding genes and candidate lncRNAs identified by RNA-seq. Each panel represents one haplotype. Colored segments on the SFBB track indicate individual S-locus F-box (SFBB) genes, color-coded by family (SFBB1–SFBB18); family numbers are indicated above each gene. The S-RNase gene is shown in orange. Grey segments indicate other protein-coding genes within the S-locus window. Grey circles on the lncRNA track represent all 83 candidate lncRNAs identified across the three haplotypes (25, 34, and 24 respectively). Pollen-specific lncRNAs — defined as significantly upregulated in pollen relative to both leaf and pistil (DESeq2, adjusted p < 0.05) — are highlighted as yellow diamonds and labelled with their transcript identifier. Six pollen-specific lncRNAs were identified in total: one in Gala haplotype 1 (MSTRG.2343), one in Gala haplotype 2 (MSTRG.2154), and four in Honeycrisp (MSTRG.2329, MSTRG.2350, MSTRG.2363, MSTRG.2384). See Supplementary Note 2 for methods.

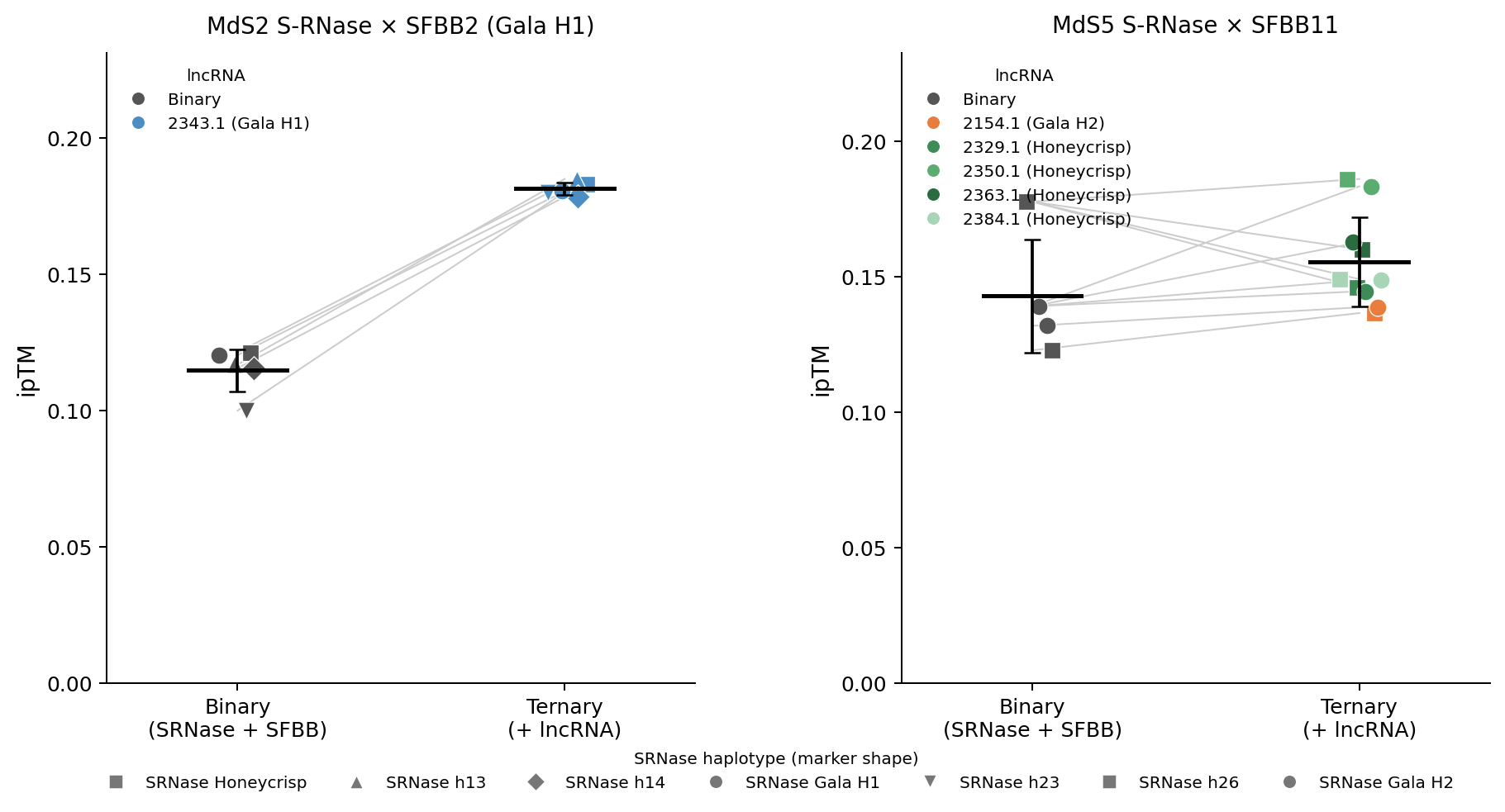

**Supplementary Fig. 29: Effect of pollen-specific lncRNAs on the predicted interaction between S-RNase and SFBB proteins, as assessed by Boltz-2 co-folding.** Each panel shows the interface predicted TM-score (ipTM) for binary complexes (S-RNase + SFBB only) and ternary complexes (S-RNase + SFBB + lncRNA). (A) MdS2 S-RNase copies tested against SFBB2 (Gala Hap1, h2) with the pollen-specific lncRNA MSTRG.2343.1 (Gala Hap1). (B) MdS5 S-RNase copies tested against SFBB11 from Honeycrisp (h10) or Gala Hap2 (h3), with the corresponding haplotype-matched pollen-specific lncRNAs. Each point represents one Boltz-2 prediction; grey lines connect the binary and ternary predictions for the same S-RNase × SFBB pair. Point colour indicates the lncRNA used in the ternary complex (grey: binary, no lncRNA); point shape indicates the S-RNase haplotype. The horizontal bar and error bars indicate the mean ± standard deviation across all predictions in each group. Higher ipTM values indicate greater predicted structural complementarity between chains. See Supplementary Note 2 for methods.

### **Supplementary Table 18: lncRNA annotation summary in the Malus S-locus (Excel file).** Summary of lncRNA discovery, coding-potential filtering and tissue-specificity steps for the three reference haplotypes (Gala haplotype 1, Gala haplotype 2, Honeycrisp), reported as the number of transcripts retained at each step (Gffcompare class assignment, length filter, CPC2/CPAT/PLEK2/RNAplonc coding-potential calls, and final majority-vote lncRNA set). See Supplementary Note 2 for methods.

### **Supplementary Table 19: Differential expression of S-locus genes across tissues (Excel file).** DESeq2 pairwise differential expression results (pistil vs. leaf, pollen vs. leaf, pollen vs. pistil) for S-locus genes (S-RNase, SFBB, lncRNA and other S-locus genes) in the three reference haplotypes (Gala haplotype 1, Gala haplotype 2, Honeycrisp). For each gene and contrast, log₂ fold-change, standard error and test statistic are reported. See Supplementary Note 2 for methods.

### **Supplementary Table 20: Boltz-2 structural prediction design for S-RNase–SFBB–lncRNA complexes (Excel file).** List of the 24 Boltz-2 co-folding predictions performed for the MdS2/SFBB2 and MdS5/SFBB11 S-allele pairs, indicating for each prediction the S-RNase allele and haplotype, the SFBB family and haplotype, the complex type (binary or ternary), and, for ternary complexes, the lncRNA and its haplotype of origin. See Supplementary Note 2 for methods.

### **Supplementary Table 21: RNA-seq mapping rates per sample (Excel file).** STAR alignment statistics for each of the 27 RNA-seq samples (three tissues × three replicates × three reference haplotypes), including total read number and the proportion of uniquely mapped, multi-mapped and total mapped reads. Underlies Supplementary Fig. 26. See Supplementary Note 2 for methods.

### **Supplementary Table 22: Coding-potential predictions for candidate lncRNA transcripts (Excel file).** Per-transcript coding-potential calls (coding/non-coding) from CPC2, CPAT, PLEK2 and RNAplonc for all candidate transcripts retained after the Gffcompare class-code and length filters, together with the majority-vote final classification and Pfam domain flag. See Supplementary Note 2 for methods.

### **Supplementary Table 23: Genomic coordinates and tissue expression of candidate S-locus lncRNAs (Excel file).** Genomic coordinates, strand, length and per-tissue mean TPM values for all candidate S-locus lncRNAs identified in the three reference haplotypes, including flags for pollen-specific and pistil-specific expression. Underlies Supplementary Fig. 27 and Supplementary Fig. 28. See Supplementary Note 2 for methods.

### **Supplementary Table 24: Boltz-2 interface predicted TM-scores (ipTM) for binary and ternary S-RNase–SFBB–lncRNA complexes (Excel file).** ipTM values for each of the 24 Boltz-2 predictions listed in Supplementary Table 20, for binary (S-RNase + SFBB) and ternary (S-RNase + SFBB + lncRNA) complexes across the MdS2/SFBB2 and MdS5/SFBB11 S-allele pairs. Underlies Supplementary Fig. 29. See Supplementary Note 2 for methods.

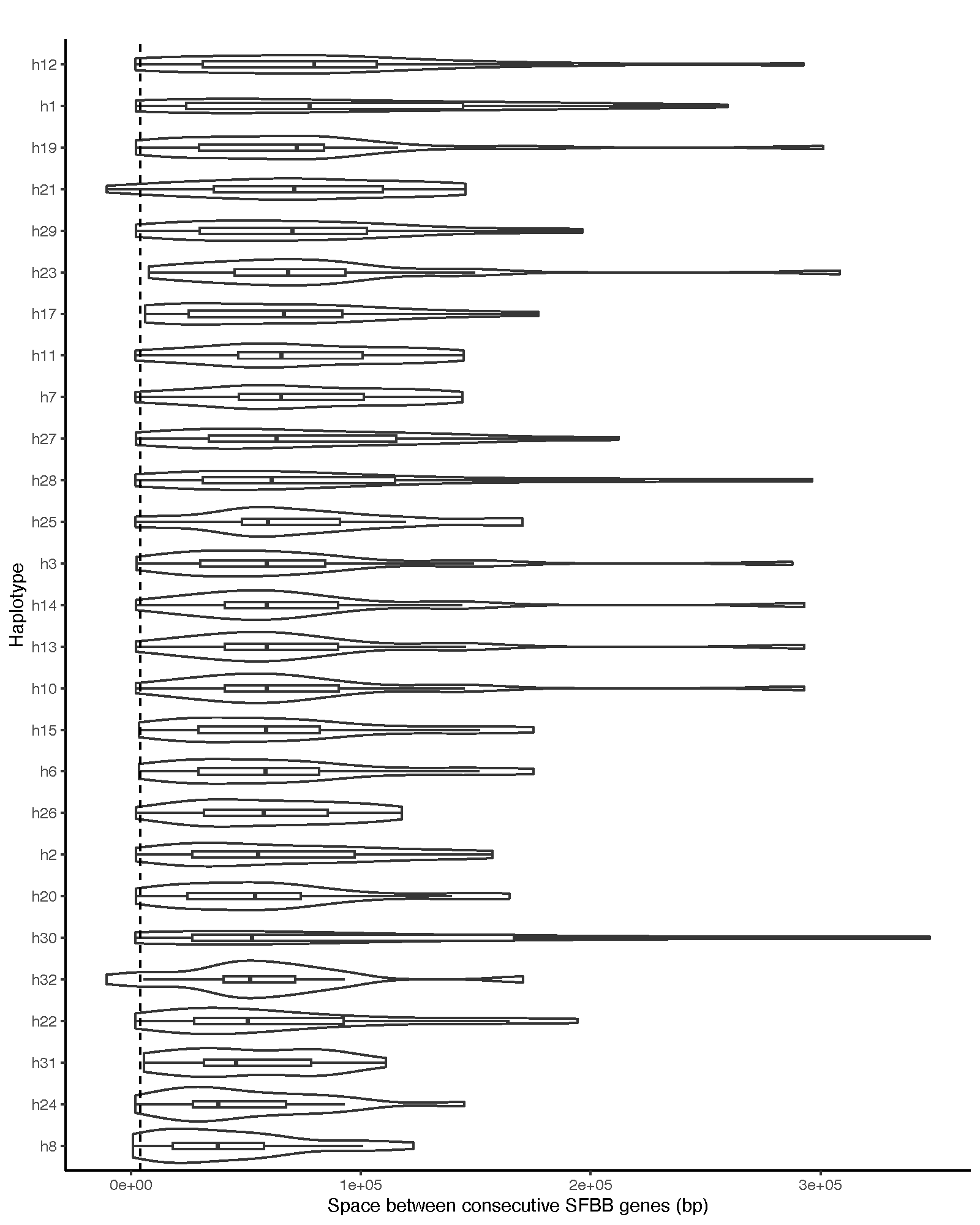

**Supplementary Fig. 30: Distribution of intergenic distances between adjacent SFBB genes across S-locus haplotypes.** Violin plots show the full distribution of distances (in base pairs), while boxplots indicate the median (horizontal line) and interquartile range. Haplotypes are ordered according to their median spacing. The dashed vertical line marks the 4 kb threshold, used as a reference for genomic proximity.

References cited

Almeida-Silva, F., Zhao, T., Ullrich, K. K., Schranz, M. E., & Van De Peer, Y. (2023). syntenet: An R/Bioconductor package for the inference and analysis of synteny networks. *Bioinformatics*, *39*(1). https://doi.org/10.1093/bioinformatics/btac806

Broothaerts, W. (2003). New findings in apple S-genotype analysis resolve previous confusion and request the re-numbering of some S-alleles. *Theoretical and Applied Genetics*, *106*(4), 703–714. https://doi.org/10.1007/s00122-002-1120-0

Daccord, N., Celton, J.-M., Linsmith, G., Becker, C., Choisne, N., Schijlen, E., Van De Geest, H., Bianco, L., Micheletti, D., Velasco, R., Di Pierro, E. A., Gouzy, J., Rees, D. J. G., Guérif, P., Muranty, H., Durel, C.-E., Laurens, F., Lespinasse, Y., Gaillard, S., … Bucher, E. (2017). High-quality de novo assembly of the apple genome and methylome dynamics of early fruit development. *Nature Genetics*, *49*(7), 1099–1106. https://doi.org/10.1038/ng.3886

Emms, D. M., & Kelly, S. (2019). OrthoFinder: Phylogenetic orthology inference for comparative genomics. *Genome Biology*, *20*(1). https://doi.org/10.1186/s13059-019-1832-y

Han, M., Sun, Q., Zhou, J., Qiu, H., Guo, J., Lu, L., Mu, W., & Sun, J. (2017). Insertion of a solo LTR retrotransposon associates with spur mutations in ‘Red Delicious’ apple (Malus × domestica). *Plant Cell Reports*, *36*(9), 1375–1385. https://doi.org/10.1007/s00299-017-2160-x

Janssens, G. A., Goderis, I. J., Broekaert, W. F., & Broothaerts, W. (1995). A molecular method for S-allele identification in apple based on allele-specific PCR. *Theoretical and Applied Genetics*, *91*(4), 691–698. https://doi.org/10.1007/BF00223298

Kasajima, I., Kikuchi, T., & Yoshikawa, N. (2017). Rapid identification of apple (*Malus*×*domestica* Borkh.) *S* alleles using sequencing-based DNA marker *APPLid*. *Plant Biotechnology*, *34*(2), 97–106. https://doi.org/10.5511/plantbiotechnology.17.0503a

Khan, A., Carey, S. B., Serrano, A., Zhang, H., Hargarten, H., Hale, H., Harkess, A., & Honaas, L. (2022). A phased, chromosome-scale genome of ‘Honeycrisp’ apple (Malus domestica). *Gigabyte*, *2022*, 1–15. https://doi.org/10.46471/gigabyte.69

Könyves, K., Mian, S., Johns, J., Royal Botanic Garden Edinburgh Genome Acquisition Lab, Royal Botanic Gardens Kew Genome Acquisition Lab, Darwin Tree of Life Barcoding collective, Wellcome Sanger Institute Tree of Life programme, Wellcome Sanger Institute Scientific Operations: DNA Pipelines collective, Tree of Life Core Informatics collective, Ruhsam, M., Leitch, I. J., & Darwin Tree of Life Consortium. (2022). The genome sequence of the apple, Malus domestica (Suckow) Borkh., 1803. *Wellcome Open Research*, *7*, 297. https://doi.org/10.12688/wellcomeopenres.18646.1

Li, W., Chu, C., Li, H., Zhang, H., Sun, H., Wang, S., Wang, Z., Li, Y., Foster, T. M., López-Girona, E., Yu, J., Li, Y., Ma, Y., Zhang, K., Han, Y., Zhou, B., Fan, X., Xiong, Y., Deng, C. H., … Han, Z. (2024). Near-gapless and haplotype-resolved apple genomes provide insights into the genetic basis of rootstock-induced dwarfing. *Nature Genetics*, *56*(3), 505–516. https://doi.org/10.1038/s41588-024-01657-2

Matsumoto, S., Furusawa, Y., Kitahara, K., & Soejima, J. (2003). Partial Genomic Sequences of S6-, S12-, S13-, S14-, S17-, S19-, and S21-RNases of Apple and Their Allele Designations. *Plant Biotechnology*, *20*(4), 323–329. https://doi.org/10.5511/plantbiotechnology.20.323

Matsumoto, S., Komori, S., Kitahara, K., Imazu, S., & Soejima, J. (1999). S-genotypes of 15 Apple Cultivars and Self-compatibility of “Megumi”. *Engei Gakkai Zasshi*, *68*(2), 236–241. https://doi.org/10.2503/jjshs.68.236

Minh, B. Q., Schmidt, H. A., Chernomor, O., Schrempf, D., Woodhams, M. D., Von Haeseler, A., & Lanfear, R. (2020). Corrigendum to: IQ-TREE 2: New Models and Efficient Methods for Phylogenetic Inference in the Genomic Era. *Molecular Biology and Evolution*, *37*(8), 2461–2461. https://doi.org/10.1093/molbev/msaa131

Qin, S., Xu, G., He, J., Li, L., Ma, H., & Lyu, D. (2023). A chromosome-scale genome assembly of Malus domestica, a multi-stress resistant apple variety. *Genomics*, *115*(3), 110627. https://doi.org/10.1016/j.ygeno.2023.110627

Ruhsam, M., Bell, D., Hart, M., Hollingsworth, P., Royal Botanic Garden Edinburgh Genome Acquisition Lab, Darwin Tree of Life Barcoding collective, Wellcome Sanger Institute Tree of Life programme, Wellcome Sanger Institute Scientific Operations: DNA Pipelines collective, Tree of Life Core Informatics collective, & Darwin Tree of Life Consortium. (2022). The genome sequence of the European crab apple, Malus sylvestris (L.) Mill., 1768. *Wellcome Open Research*, *7*, 296. https://doi.org/10.12688/wellcomeopenres.18645.1

Sakurai, K., Brown, S. K., & Weeden, N. (2000). *Self-incompatibility Alleles of Apple Cultivars and Advanced Selections*. *35*(1), 116–119.

Sassa, H., Mase, N., Hirano, H., & Ikehashi, H. (1994). *Identification of self-incompatibility-related glycoproteins in styles of apple ( Malus x domestica)*. *89*, 201–205.

Schneider, D., Stern, R., Eisikowitch, D., & Goldway, M. (2001). Analysis of S-alleles by PCR for determination of compatibility in the ‘Red Delicious’ apple orchard. *The Journal of Horticultural Science and Biotechnology*, *76*(5), 596–600.

Sievers, F., & Higgins, D. G. (2014). Clustal Omega. *Current Protocols in Bioinformatics*, *48*(1). https://doi.org/10.1002/0471250953.bi0313s48

Sun, X., Jiao, C., Schwaninger, H., Chao, C. T., Ma, Y., Duan, N., Khan, A., Ban, S., Xu, K., Cheng, L., Zhong, G.-Y., & Fei, Z. (2020). Phased diploid genome assemblies and pan-genomes provide insights into the genetic history of apple domestication. *Nature Genetics*, *52*(12), 1423–1432. https://doi.org/10.1038/s41588-020-00723-9

Tian, Y., Thrimawithana, A., Ding, T., Guo, J., Gleave, A., Chagné, D., Ampomah‐Dwamena, C., Ireland, H. S., Schaffer, R. J., Luo, Z., Wang, M., An, X., Wang, D., Gao, Y., Wang, K., Zhang, H., Zhang, R., Zhou, Z., Yan, Z., … Yao, J. (2022). Transposon insertions regulate genome‐wide allele‐specific expression and underpin flower colour variations in apple ( *Malus* spp.). *Plant Biotechnology Journal*, *20*(7), 1285–1297. https://doi.org/10.1111/pbi.13806
